# Rewired type I IFN signaling is linked to age-dependent differences in COVID-19

**DOI:** 10.1101/2024.10.23.619479

**Authors:** Lev Petrov, Sophia Brumhard, Sebastian Wisniewski, Philipp Georg, David Hillus, Anna Hiller, Rosario Astaburuaga-García, Nils Blüthgen, Emanuel Wyler, Katrin Vogt, Hannah-Philine Dey, Saskia von Stillfried, Christina Iwert, Roman D. Bülow, Bruno Märkl, Lukas Maas, Christine Langner, Tim Meyer, Jennifer Loske, Roland Eils, Irina Lehmann, Benjamin Ondruschka, Markus Ralser, Jakob Trimpert, Peter Boor, Sammy Bedoui, Christian Meisel, Marcus A. Mall, Victor M. Corman, Leif Erik Sander, Jobst Röhmel, Birgit Sawitzki

**Author notes:** Senior author (Jobst Röhmel, Birgit Sawitzki).

## Abstract

Advanced age is the most important risk factor for severe disease or death from COVID-19, but a thorough mechanistic understanding of the molecular and cellular underpinnings is lacking. Multi-omics analysis of samples from SARS-CoV-2 infected persons aged 1 to 84 years, revealed a rewiring of type I interferon (IFN) signaling with a gradual shift from signal transducer and activator of transcription 1 (STAT1) to STAT3 activation in monocytes, CD4^+^ T cells and B cells with increasing age. Diversion of interferon IFN signaling was associated with increased expression of inflammatory markers, enhanced release of inflammatory cytokines, and delayed contraction of infection-induced CD4^+^ T cells. A shift from IFN-responsive germinal center B (GCB) cells towards CD69^high^ GCB and atypical B cells corresponded to the formation of IgA in children while complement fixing IgG was dominant in adults. Our data provide a mechanistic basis for inflammation-prone responses to infections and associated pathology during aging.

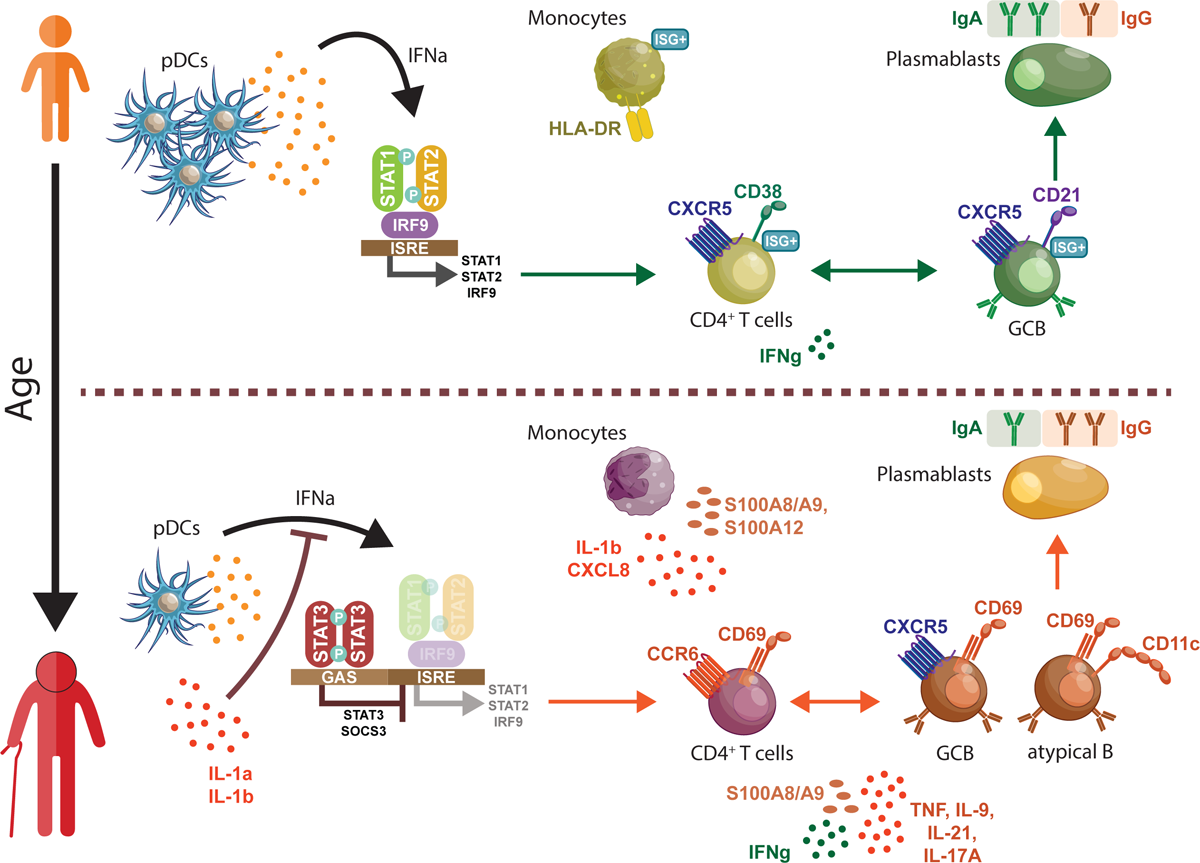

## Introduction

The COVID-19 pandemic has highlighted age-dependent disparities in anti-viral immune responses and susceptibility to severe infections, where advanced age is the most important risk factor for the development of severe COVID-19.^1–3^ Young children were rarely affected by severe lung failure and generally experience mild or asymptomatic disease courses.^4–7^

Our previous investigations revealed more pronounced type I interferon (IFN) responses and interferon-stimulated gene (ISG) signatures in epithelial and innate immune cells of SARS-CoV-2 infected children.^8^ While this points to an earlier and enhanced viral control, available data suggest comparable viral loads and clearance kinetics in children and adults.^8,9^ Thus, the enhanced local innate immunity and better mucosal virus control may at least partly explain the age-dependent differences in disease severity.

The adaptive immune system is crucial for sustained control of SARS-CoV-2 infection.^10^ T cells are required for viral clearance and neutralizing antibodies offer some protection from reinfection.^11^

Mild and asymptomatic courses of COVID-19 leads to seroconversion, the development of neutralizing antibodies, and SARS-CoV-2 specific T cells in the majority of children, but it is rarely associated with systemic inflammation as observed in infected adults, with the exception of PIMS.^10,12–16^

In contrast, severe disease, most frequently observed in the elderly, is driven largely by immune pathology involving dysregulated myeloid cells and highly cytotoxic T cells.^17,18^ This suggests qualitative alterations in immune cell activation upon SARS-CoV-2 infection between children and elderly individuals.

To reveal age-dependent differences in primary SARS-CoV-2 infection we performed a multi-omics-based characterization of immune responses in SARS-CoV-2 infected children and adults sampled in existing cohorts.^1,19,20^ The RECAST cohort allowed us to directly compare immune cell profiles in infected children and their adult family members.^1^ *Ex vivo* profiling was complemented by functional *in vitro* studies using immune cells from uninfected controls.

We found that age was associated with a continuous rewiring in type I IFN signaling from STAT1-dominated to STAT3-driven signal propagation in monocytes, T and B cells. This phenomenon was associated with a gradual change in infection-induced expression of inflammatory markers across multiple leukocyte types at a systemic and local level. These shifts were linked to an increased inflammatory immune response, including the expansion of *S100A8/A9-* and *CXCL8*-expressing HLA-DR^low^ monocytes, increased peripheral CCR6^+^ T helper cells and CD69-expressing germinal center and atypical B cells in older individuals. In contrast, in children we observed a more focused and constrained adaptive response, characterized by the exclusive preference for follicular responses, evidenced by increase of CD38^high^CXCR5^high^ T follicular helper and germinal center B cell differentiation, and a faster contraction of T cell activation.

Overall, our study provides a detailed systematic characterization of innate and adaptive immune responses to SARS-CoV-2 infection across the entire age spectrum. It reveals an age-dependent rewiring of type-I IFN signaling with a gradual shift from antiviral STAT1-signaling towards STAT3-driven inflammation with older age. These findings provide a mechanistic explanation for the often-observed increased susceptibility of older adults to severe viral infections.

## Results

### Up-regulation of HLA-DR versus CD10 distinguishes infection-induced monocytes in children and adults

To study the age-dependent differences in anti-SARS-CoV-2 immune cell responses, we used samples and clinical data from three observational study cohorts (Figure 1A). RECAST allowed us to perform a multi-omics-based profiling of specimens collected from non-hospitalized infected and control families spanning an age range from 1 to 68 years.^1^ Control patients were tested negative for SARS-CoV-2 and had no clinical signs of an infection. Analysis of specimens collected from PaCOVID and prospective vaccination trial EICOV/COVIMMUNIZE studies enabled integration of data from even older and / or hospitalized patients and controls (up to 86 years).^19,20^

**Figure 1.**
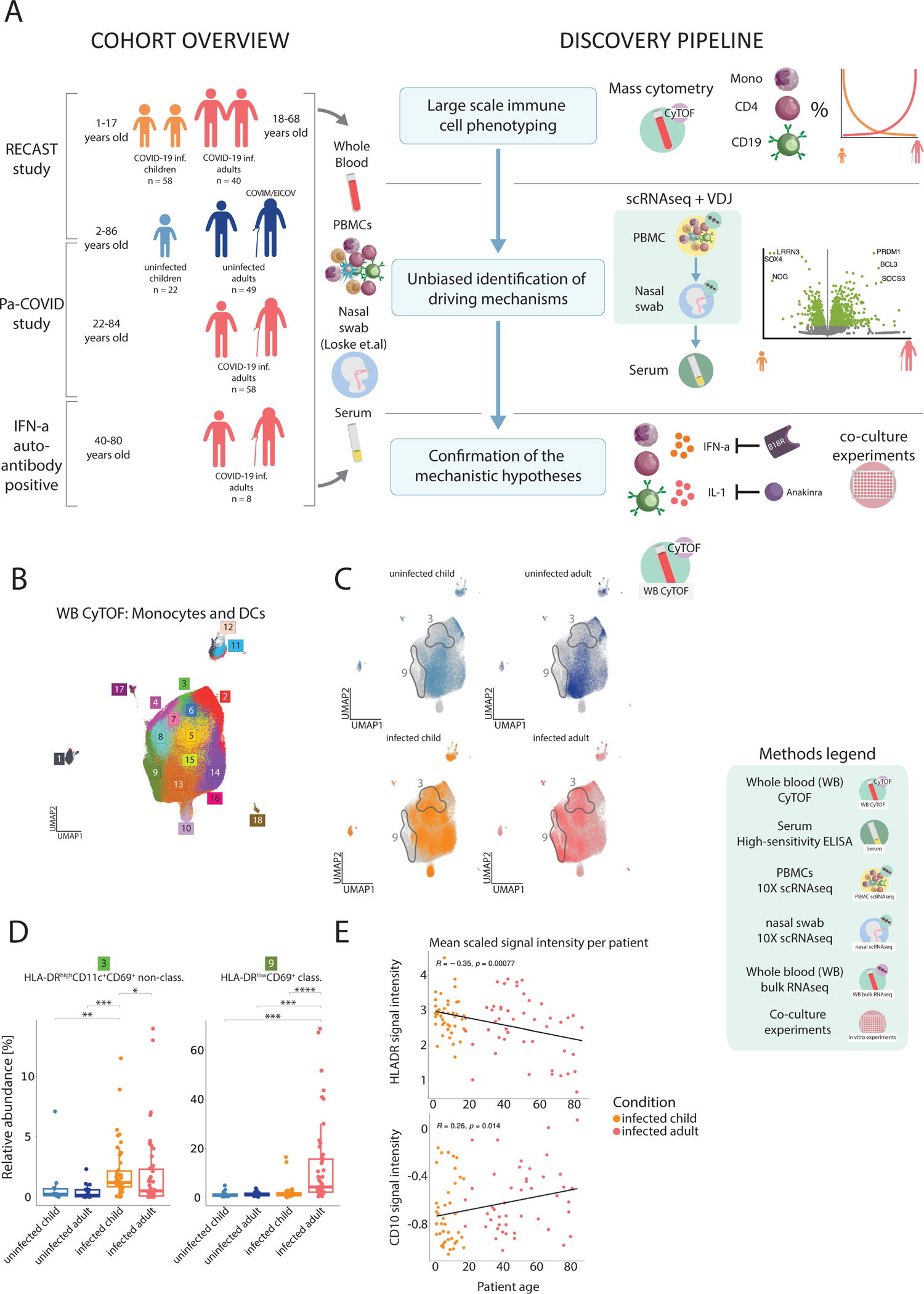
Distinct age-dependent patterns of monocyte activation are characterized by opposite HLA-DR and CD11c versus CD10 expression. (A) Overview of the study cohort and methodological pipeline. Blood samples were collected from SARS-CoV-2 infected children and adults as well as controls spanning an age range from 1 to 86 years. Whole blood CyTOF and scRNA-seq combined with VDJ-seq-based clonotype identification were used to determine age-specific alterations in the monocyte, T and B cell compartment. The obtained results together with serum antibody profiles were used to develop hypotheses on their functional properties, inducing mechanisms and transcriptional control, which were tested in *ex vivo* cultures. Detailed information on samples included in all reported assays can be found in Table S1. Additional cohort summary is included in Table S2. Multiple icons are used throughout the paper to identify data from different experiments (Methods legend in the lower right corner of the figure). (B) UMAP, showing pre-gated monocyte and dendritic cell (DC) populations from the CyTOF data set. 18 clusters have been produced using a semi-supervised approach with FlowSOM algorithm. UMAP presents all the patients included in the dataset and used for clustering: uninfected children (RECAST n = 13; median age = 7), uninfected adults (RECAST n = 25; median age = 48), infected children (RECAST study n = 48; median age = 8; asymptomatic n = 11, symptomatic n = 37) and adults (RECAST n = 21, PA-COVID study n = 30; median age = 47; mild n = 37, severe n = 10, severe with IFN autoantibodies n = 4). Only samples measured during the acute phase of infection (defined as first 14 days after symptom onset) were included. (C) UMAPs, showing the location of cells belonging to the respective group (colored in, whereas grey identifies cells in other groups). Grey outlines indicate cluster regions enriched in infected children or adults. Severe patients with IFN autoantibodies were excluded from the analysis. (D) Box plots, showing relative abundance of infection-induced clusters resulting from the FlowSOM algorithm, calculated per sample within all monocytes and dendritic cells (DC) from the CyTOF data. Severe patients with IFN autoantibodies were excluded from the analysis. Kruskal-Wallis + Wilcoxon; *, p < 0.05; **, p < 0.01; ***, p < 0.001; ****, p < 0.0001. (E) Scatter plots, showing mean normalized HLA-DR and CD10 signals in relationship to patient age for infected patients (linear model fitted to data and Spearman’s rank correlation coefficient shown in black) within monocyte and DC CyTOF data. Severe patients with IFN autoantibodies were excluded from the analysis.

We focused on patient samples collected within the first two weeks post-symptom onset or infection in case of asymptomatic children to reveal immune cell response patterns explaining age-dependent differences in early symptom severity (infected children mean days post-symptom onset - DPSO = 5±3.3 infected adults mean DPSO = 7.7±4.5). For definition of COVID-19 severity we followed the established WHO ordinal scale as previously described.^17,18,21^ Tables S1 and S2 report the maximal WHO ordinary scale, DPSO and comorbidity status per infected patient and patient group, respectively.

Since we had previously identified dysregulated myeloid cells to be a feature of severe COVID-19 we investigated whether monocytes and dendritic cells (DCs) of children and adults responded differently to the viral challenge by performing mass cytometry (CyTOF) on whole blood samples (Figure 1B-E, Figure S1A-C).^17^ We identified plasmacytoid dendritic cells (CD14^-^ CD123^+^CD11c^-^), conventional dendritic cells (cDC) (CD14^-^CD123^-^CD11c^+^), classical monocytes (CD14^high^CD16^-^), intermediate monocytes (CD14^high^CD16^low^, non-classical monocytes (CD14^low^CD16^high^) and Ki67^+^ hematopoietic stem and progenitor cells (HSPC) using FlowSOM clustering of the CD45^+^CD15^-^CD3^-^CD56^-^CD19^-^ pre-gated cells. SARS-CoV-2 infection shifted the composition of the DC and monocyte clusters in infected children and adults (Figure 1C,B, Figure S1A,B). Samples from infected children contained relatively more pDCs and cDCs compared to those from infected adults (Figure S1B). Interestingly, we also detected enhanced proportions of non-classical monocytes characterized by high HLA-DR and CD11c expression (e.g. cluster 3 HLA-DR^high^CD11c^+^CD69^+^ non-class., Figure 1D, Figure S1A,B) in samples from infected children. In contrast, infected adults showed an expansion of HLA-DR^low^, CD10, CD69-expressing classical monocytes (e.g. cluster 9, Figure 1D, S1A,B). Consequently, the whole immune cell space of monocytes and DCs varied between children and adults. Expression of HLA-DR, CD11c, CD123 was increased in children, whereas CD69, CD226, CD10 and CXCR3 were elevated in infected adults (Figure S1C). HLA-DR expression levels across all monocyte populations were negatively correlated with patient age, with a pronounced decline in expression beyond the age of 60 (Figure S1E, upper plot). In contrast, expression of the macrophage marker CD10 linearly increased with age (Figure S1E, lower plot). Thus, monocytes respond differently to SARS-CoV-2 infection, depending on age.

### Gradual age-dependent changes in the phenotype of infection-induced CD4^+^ T cells and B cells

We next determined whether different responses of monocytes in SARS-CoV-2 infected children and adults were associated with differences in the T cell responses. We performed subclustering of CD4^+^ T cells from the CyTOF data (Figure 2A; Figure S2A). This resulted in identification of 19 CD4^+^ T cell clusters, which were grouped based on CD62L and CD45RO expression into naïve (Naïve, CD62L^+^CD45RO^-^), central memory (CM, CD62L^+^, CD45RO^+^), effector memory (EM, CD62L^-^CD45RO^+^) and Effector Memory–Expressing CD45RA / terminally differentiated effector memory T cells (TEMRA, CD62L^-^CD45RO^-^). In accordance with our previous findings we also identified CD4^+^ T cell clusters with high expression of CCR6, CD69 and CD16 (clusters 11 TIGIT^+^CD137^+^HLA-DR^+^CCR6^+^ CM and 15 Lag3^+^TIGIT^+^CD69^+^CCR6^+^CD16^+^ TEMRA).^18^ These T cells were increased in infected adults in comparison to infected children or controls (Figure 2B,C; Figure S2B). In contrast, infected children had higher frequencies of CD38^high^CXCR5^high^ CD4^+^ T cells (cluster 8), indicative of expansion of T follicular helper cells and CD38^+^ naïve CD4^+^ T cells (cluster 12). Consequently, higher signal expression intensity of CD38 across all CD4^+^ T cells was detected in infected children, whereas CD4^+^ T cells from infected adults showed higher CCR6 expression (Figure S2C). CD38 expression was negatively correlated with age and the opposite pattern was observed for CCR6 (Figure 2D). The age-dependent differences in CD38 down-regulation and CCR6 up-regulation followed a linear, gradual trend with patients at the age of 40 showing a balanced increase of both markers (Figure 2D). This age-dependent divergent response pattern of T cells was also observed between children and parents of the same family/household (Figure S2D). This argues against genetic or local environmental covariates as key drivers of these phenotypic changes in T cells although their role cannot be ruled out completely.

**Figure 2.**
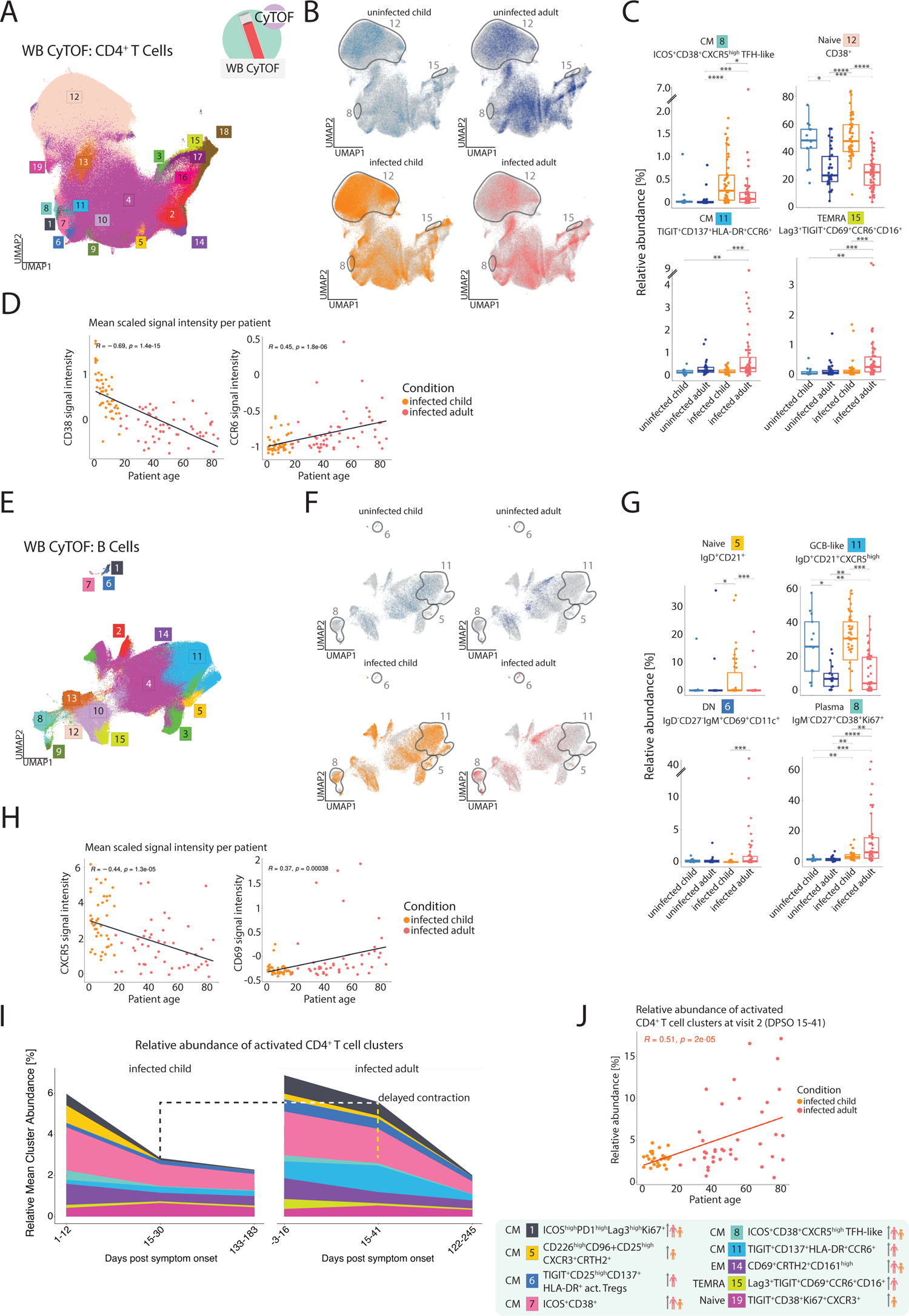
Gradual age-dependent change in the phenotype of infection-induced CD4^+^ T cells and B cells. (A) UMAP, showing pre-gated CD4^+^ T cells from the CyTOF data set. 19 clusters have been produced using a semi-supervised approach with FlowSOM algorithm. UMAP presents all the patients that were part of the dataset and used for clustering, including follow-up measurements of some patients done approximately two weeks and six months after the first, acute infection phase measurement. Patients groups used for clustering were: uninfected children (RECAST n = 13; median age = 7), uninfected adults (RECAST n = 25; median age = 48), infected children (RECAST study n = 53; median age = 8; asymptomatic n = 11, symptomatic n = 42) and adults (RECAST n = 24, PA-COVID study n = 51; median age = 51; mild n = 45, severe n = 24, severe with IFN autoantibodies n = 6). (B) UMAPs, showing the location of cells belonging to the respective group (colored in, whereas grey identifies cells in other groups). Grey outlines indicate cluster regions enriched in infected children or adults. Importantly, only samples measured during the acute phase of infection (defined as first 14 days after symptom onset) are shown. Severe patients with IFN autoantibodies were excluded from the analysis. Group composition is thus as follows: uninfected children (RECAST n = 13; median age = 7), uninfected adults (RECAST n = 25; median age = 48), infected children (RECAST study n = 48; median age = 8; asymptomatic n = 11, symptomatic n = 37) and adults (RECAST n = 21, PA-COVID study n = 26; median age = 45; mild n = 37, severe n = 10). (C) Box plots, showing relative abundance of infection-induced clusters resulting from the FlowSOM algorithm, calculated per sample within all CD4^+^ T cells from the CyTOF data. Only samples measured during the acute phase of infection (defined as first 14 days after symptom onset) are shown. Severe patients with IFN autoantibodies were excluded from the analysis. Group composition is thus as follows: uninfected children (RECAST n = 13; median age = 7), uninfected adults (RECAST n = 25; median age = 48), infected children (RECAST study n = 48; median age = 8; asymptomatic n = 11, symptomatic n = 37) and adults (RECAST n = 21, PA-COVID study n = 26; median age = 45; mild n = 37, severe n = 10). Kruskal-Wallis + Wilcoxon; *, p < 0.05; **, p < 0.01; ***, p < 0.001; ****, p < 0.0001. (D) Scatter plots, showing mean normalized CD38 and CCR6 signals in relationship to patient age for infected patients (linear model fitted to data and Spearman’s rank correlation coefficient shown in black) within CD4^+^ T cell CyTOF data. Only samples measured during the acute phase of infection (defined as first 14 days after symptom onset) are shown. Severe patients with IFN autoantibodies were excluded from the analysis. (E) UMAP, showing pre-gated B cells from the CyTOF data set. 15 clusters have been produced using a semi-supervised approach with FlowSOM algorithm. UMAP presents all the patients included in the dataset and used for clustering: uninfected children (RECAST n = 13; median age = 7), uninfected adults (RECAST n = 25; median age = 48), infected children (RECAST study n = 48; median age = 8; asymptomatic n = 11, symptomatic n = 37) and adults (RECAST n = 21, PA-COVID study n = 30; median age = 47; mild n = 37, severe n = 10, severe with IFN autoantibodies n = 4). Only samples measured during the acute phase of infection (defined as first 14 days after symptom onset) were included.

Taken together, CD4^+^ T cells from children and adults responded differently to the viral infection with a gradual shift from CD38^high^CXCR5^+^ follicular T cells in children to CCR6^+^CD69^+^ peripheral T helper-like cells in adults.

We next studied phenotypical differences in the B cell compartment in SARS-CoV-2 infected children and adults. We performed a subclustering of CD19^+^ B cells (Figure 2E; Figure S3A). This resulted in identification of 15 B cell clusters, which, based on their IgD, IgM, CD27, CD10, CD38 and CXCR5 expression, were grouped into transitional (Transit., CD27^-^CD10^high^CD38^high^), naïve (Naive, IgD^+^IgM^+^CD27^-^), GCB-like cells (GCB-like, CXCR5^high^), double negative-like (DN, IgD^-^CD27^-^), memory (CS mem, CD27^+^CD38^low^) and plasmablasts / antibody secreting cells (Plasma, CD27^+^CD38^high^).

In samples of infected adults, we observed increased proportions of CD69-expressing B cell populations (clusters 1 IgD^-^CD27^-^IgM^+^CD21^-^CD69^+^CD11c^+^ and 6 IgD^-^CD27^-^IgM^+^CD21^-^ CD69^+^CD11c^+^ DN-like B cells, cluster 14 IgD^+^CD21^low^CXCR5^high^CD69^+^ GCB-like naïve B cells; Figure 2F,G, Figure S3B). Cluster 1 and cluster 6 B cells were negative for IgD, CD27, and CD21, characteristic of atypical / extrafollicular B cells.^22^ They also expressed CD11c and thus resembled DN2-like B cells.^22,23^ B cells belonging to cluster 14 however, expressed low levels of IgD and CD21 but high levels of CXCR5 and were annotated as GCB-like naïve B cells. Thus, both follicular and extrafollicular B cell responses were promoted in infected adults, resulting in strong up-regulation of CD69. The induction of both types of B-cell responses was associated with a higher frequency of IgM negative plasmablasts (cluster 8 IgM^-^ CD27^+^CD38^+^Ki67^+^ plasmablasts; Figure S3D-F).

In contrast, in samples from SARS-CoV-2 infected children we detected significantly higher proportions of B cell clusters 3 (CD10^high^CD38^high^TIGIT^-^ transitional-like B cells), 5 (IgD^+^CD21^+^ naïve like B cells) and 11 (IgD^+^CD21^+^CXCR5^high^ GCB-like naïve B cells) suggesting an exclusive induction of follicular B cell responses (Figure 2G; S3B,D). Consequently, higher signal expression intensity of IgD across all B cells was detected in infected children (Figure S3D).

Like for the CD4^+^ T cells, the opposing phenotypes were also apparent between children and their same parents (Figure S3C) and followed a gradual change with increasing age, where average expression of CXCR5 and CD21 was negatively correlated and CCR6 expression positively correlated with age (Figure 2H, Figure S3E).

As the sampling time (DPSO) was not perfectly matched in infected children and adults, we assessed whether this impacted the determined age-specific immune cell phenotypes. However, our cohort contained 11 asymptomatic cases out of 58 infected children.^24–26^ It was not possible to assign a DPSO value to these cases and infection dates could not be determined reliably.

We plotted our CyTOF data against DPSO in infected symptomatic children and adults. None of the clusters that we describe here as being age-related showed consistent correlation with DPSO in adults and children (Mendeley Figure 1).

Additionally, we analyzed samples collected at later timepoints (visit 2: DPSO children = 15-30, adults = 14-41, visit 3: DPSO children = 133-183, adults = 122-245) and did not detect a reversal of infection induced CD4^+^ T cell clusters in neither children nor adults (Figure 2I, J). For example, the children-specific CyTOF clusters 5 (CD226^high^CD96^+^CD25^high^CXCR3^+^CRTH2^+^) and 8 (ICOS^+^CD38^+^CXCR5^high^ TFH-like) did not increase with time in adults, whereas adult-specific clusters 11 (TIGIT^+^CD137^+^HLA-DR^+^CCR6^+^) and 15 (Lag3^+^TIGIT^+^CD69^+^CCR6^+^CD16^+^) remained low in children. Furthermore, although the overall frequency of activated CD4^+^ T cells was similar at the first sampling timepoint, children displayed a faster CD4^+^ T cell contraction compared to adults (Figure 2J).

Plotting average signal intensities per patient of infection-induced surface markers for monocytes, CD4^+^ T cells and B cells (Figure S3F) confirmed that some markers show age-dependent expression patterns across multiple leukocyte types. For example, CD38 and CXCR5 were higher in CD4^+^ T cells and B cells in children, while CD69 expression was higher in adult lymphocytes. CCR6, was higher in monocytes and CD4^+^ T cells of infected adults. This confirmed a systematic shift in infection-induced activation with age.

### Age-dependent shift from type I IFN responsive to inflammatory monocytes

Up-regulation of HLA-DR expression (Figure 1; Figure S1) can be induced by type I and II IFN signaling.^27^ To investigate potential differences in IFN responsiveness between cells from children and adults, we performed scRNAseq of PBMC samples (Figure S4A,B,C) with subsequent subclustering of the monocytes (Figure 4A; Figure S4D).

HLA-DR^low^ classical monocytes were increased in infected adults (clusters 2, 4), whereas IFN stimulated gene (ISG)^+^ intermediate (cluster 1) and non-classical (cluster 7) monocytes were only observed in children (Figure 3A,B; Figure S4D,E). Genes characterizing those clusters included *IFI27*, *IFI6* and *ISG15* but also *SIGLEC1*, which had been previously linked with mild COVID-19.^28^ Importantly, type I IFN serum concentrations were indistinguishable between children and adults, indicating a difference in IFN responsiveness rather than in IFN production. (Figure S4G).

**Figure 3.**
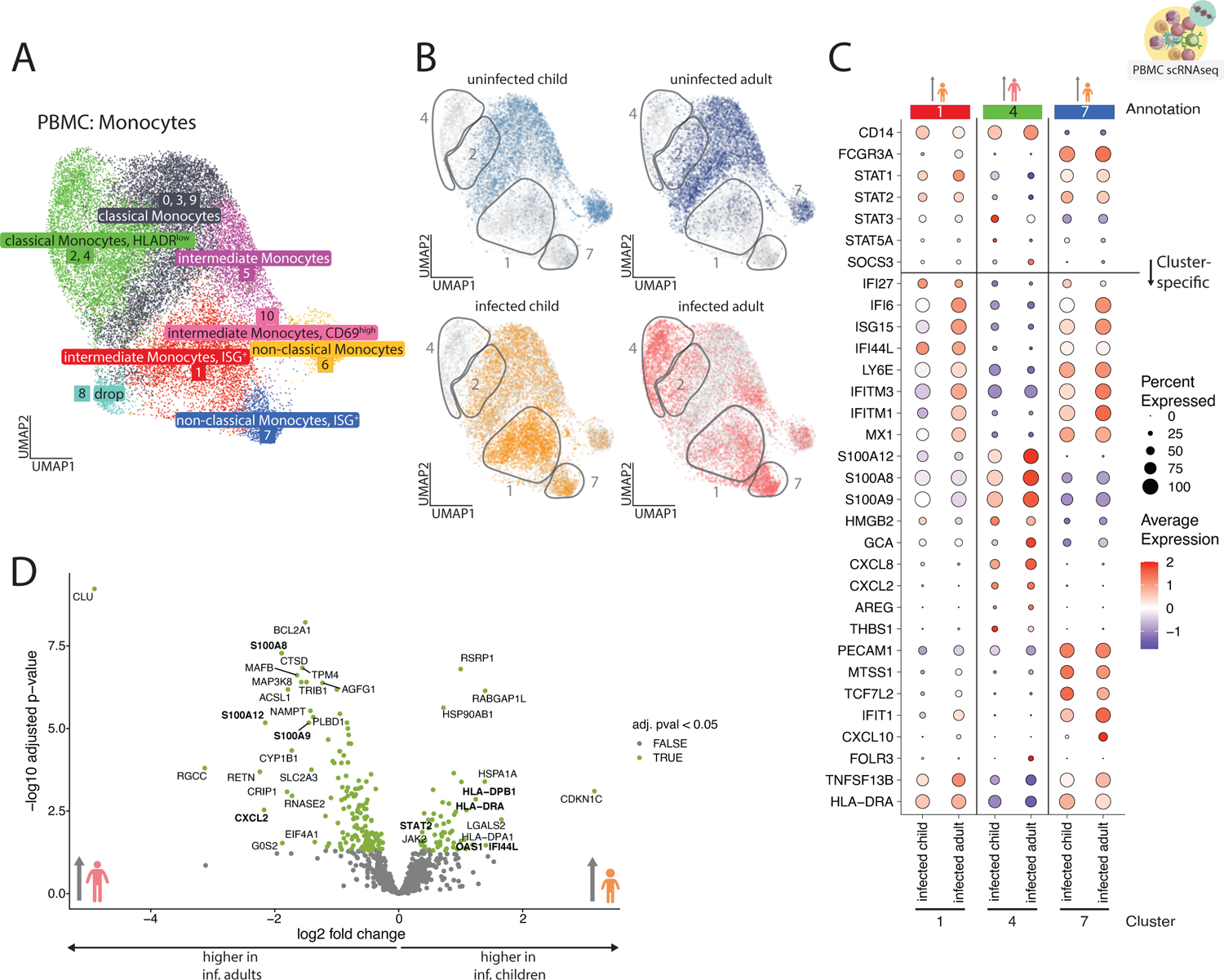
Age-dependent shift from type I IFN responsiveness to increased inflammatory potential. (A) UMAP, showing monocyte cells, subset from the PBMC scRNAseq data. 11 clusters have been produced using a graph-based approach as implemented in Seurat package (KNN graph with Louvain community detection). Some clusters share annotation due to phenotypical similarity and accumulation pattern and are treated as one population for abundance testing in S3E. Cluster annotated as “dropped” was of low quality (as concluded from inspecting number of features, n of counts and percentage of mitochondrial genes as well as other population-specific genes). UMAP presents all the patients included in the dataset and used for clustering: uninfected children (RECAST study n = 5; median age = 7), uninfected adults (RECAST study n = 4, EICOV/COVIMMUNIZE study n = 4; median age = 66), infected children (RECAST study n = 13; median age = 9; asymptomatic n = 5, symptomatic n = 8) and infected adults (RECAST study n = 1; PA-COVID study n = 18; median age = 67; mild n = 5, severe n = 7, severe with IFN autoantibodies n = 7). (B) UMAPs, showing the location of cells belonging to the respective group (colored in, whereas grey identifies cells in other groups). Grey outlines indicate cluster regions enriched in infected children or adults. Severe patients with IFN autoantibodies were excluded from the analysis. Patients with DPSO range from 1 to 19 days were included (median of 8.5, inf. children median = 4.5, inf. adult median = 10). (C) Dotplot, showing scaled average expression of genes in monocyte clusters, increased with infection (1, 4 and 7). Clusters are split between infected children and infected adults. Severe patients with IFN autoantibodies were excluded from the analysis. A horizontal line splits the dotplot in two parts – genes above the line were curated based on the presence of clusters with pronounced ISG signature and include other genes useful for annotation, genes below the line were found to be differentially expressed between the clusters (FindMarkers Seurat function). Patient group color-coded figurines on the top point out cluster accumulation patterns as seen in S4E. (D) Volcano plot, showing the results of differential expression (DE) analysis of monocytes from clusters expanded with infection (1, 4 and 7) using DESeq2. Significantly enriched genes (baseMean counts over 50 and adjusted p value < 0.05, a total of 189 genes), are colored green and 20 significant genes with highest absolute log2 fold change are labeled. Additional significant genes, tangential to other findings, were labelled ad-hoc. Positive log2 fold change values mean an enrichment in pediatric infected patients and negative – in infected adults. Severe patients with IFN autoantibodies were excluded from the analysis.

**Figure 4.**
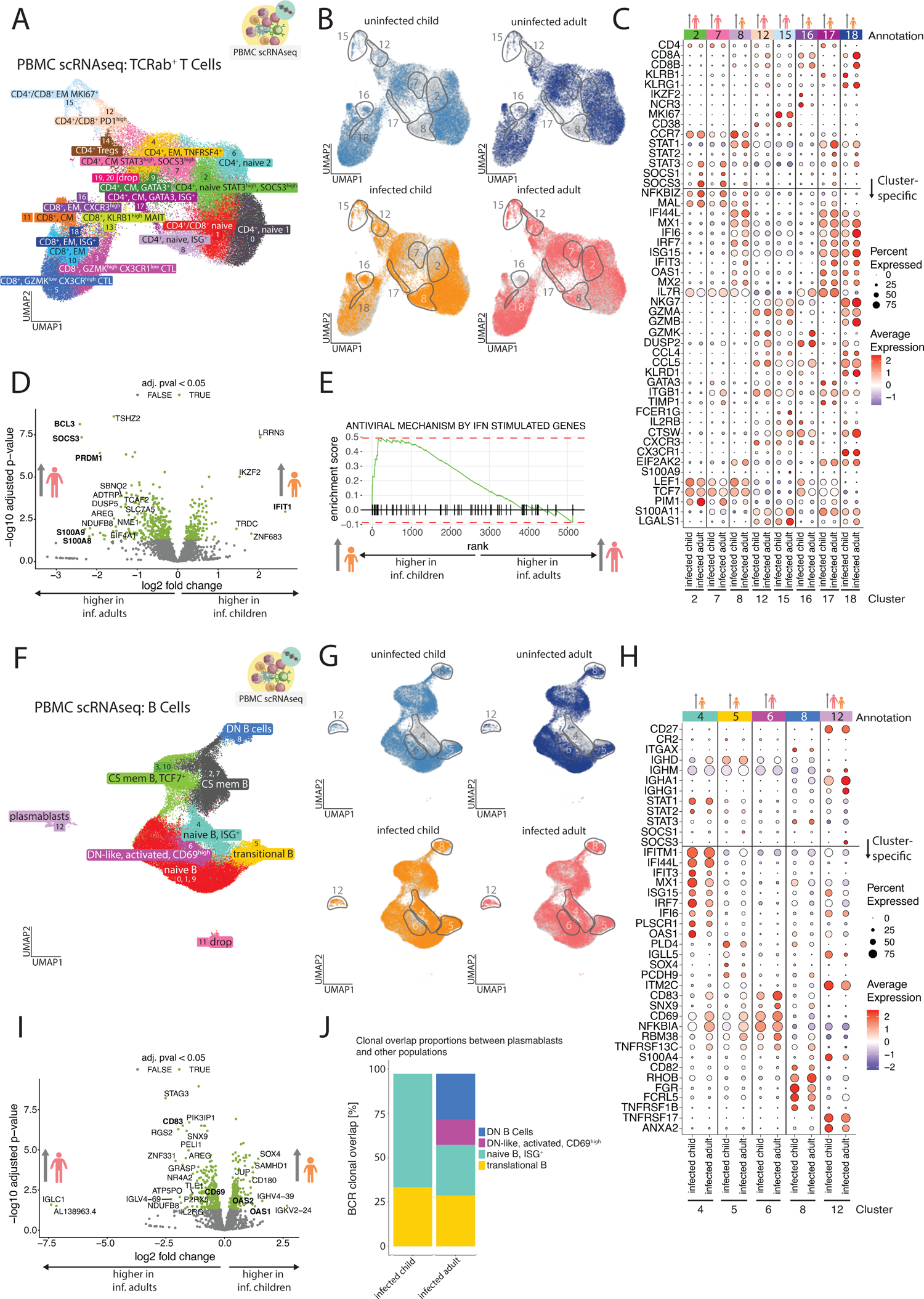
Molecular characterization of infection-associated TCRab^+^ T Cells and B cells indicates increased STAT3 involvement and altered differentiation during ageing. (A) UMAP, showing TCRab^+^ T cells, subset from the PBMC scRNAseq data. 21 clusters have been produced using a graph-based approach as implemented in Seurat package (KNN graph with Louvain community detection). Clusters annotated as “dropped” were of low quality (as concluded from inspecting number of features, n of counts and percentage of mitochondrial genes as well as other population-specific genes). UMAP presents all the patients included in the dataset and used for clustering: uninfected children (RECAST study n = 5; median age = 7), uninfected adults (RECAST study n = 4, EICOV/COVIMMUNIZE study n = 4; median age = 66), infected children (RECAST study n = 13; median age = 9; asymptomatic n = 5, symptomatic n = 8) and infected adults (RECAST study n = 1; PA-COVID study n = 18; median age = 67; mild n = 5, severe n = 7, severe with IFN autoantibodies n = 7). (B) UMAPs, showing the location of cells belonging to the respective group (colored in, whereas grey identifies cells in other groups). Grey outlines indicate cluster regions enriched in infected children or adults. Severe patients with IFN autoantibodies were excluded from the analysis. Patients with DPSO range from 1 to 19 days were included (median of 8.5, inf. children median = 4.5, inf. adult median = 10). (C) Dotplot, showing scaled average expression of genes in TCRab^+^ T cell clusters, increased with infection (2, 7, 8, 12, 15, 16, 17 and 18). Clusters are split between infected children and infected adults. Severe patients with IFN autoantibodies were excluded from the analysis. A horizontal line splits the dotplot in two parts – genes above the line were curated based on the presence of clusters with pronounced ISG signature and include other genes useful for annotation, genes below the line were found to be differentially expressed between the clusters (FindMarkers Seurat function). Patient group color-coded figurines on the top point out cluster accumulation patterns as seen in S5B. (D) Volcano plot, showing the results of differential expression (DE) analysis of TCRab^+^ T cells from clusters expanded with infection (2, 7, 8, 12, 15, 16, 17 and 18) using DESeq2. Significantly enriched genes (baseMean counts over 50 and adjusted p value < 0.05, a total of 464 genes), are colored green and 20 significant genes with highest absolute log2 fold change are labeled. Additional significant genes, tangential to other findings, were labelled ad-hoc. Positive log2 fold change values mean an enrichment in pediatric infected patients and negative – in infected adults. Severe patients with IFN autoantibodies were excluded from the analysis. (E) Gene set enrichment analysis (GSEA) applied to TCRab^+^ T cells from the PBMC scRNAseq experiment and clusters, expanded with infection (2, 7, 8, 12, 15, 16, 17 and 18). GSEA was done using genes from the infected adults – infected children comparison with baseMean counts > 50 (total of 5181 genes). R package fgsea was used with Reactome gene list R-HSA-1169410: Antiviral mechanism by IFN-stimulated genes (listed in Supplementary Table 3). Genes are shown as ticks on a bold black line and ranked by log2 fold change. The green line shows a trend in enrichment score, where positive trend (upwards) means that the gene is enriched in reference group – infected children. Severe patients with IFN autoantibodies were excluded from the analysis. (F) UMAP, showing B cells, subset from the PBMC scRNAseq data. 13 clusters have been produced using a graph-based approach as implemented in Seurat package (KNN graph with Louvain community detection). Some clusters share annotation due to phenotypical similarity and accumulation pattern and are treated as one population for abundance testing in S4E. Cluster annotated as “dropped” was of low quality (as concluded from inspecting number of features, n of counts and percentage of mitochondrial genes as well as other population-specific genes). UMAP presents all the patients included in the dataset and used for clustering: uninfected children (RECAST study n = 5; median age = 7), uninfected adults (RECAST study n = 4, EICOV/COVIMMUNIZE study n = 4; median age = 66), infected children (RECAST study n = 13; median age = 9; asymptomatic n = 5, symptomatic n = 8) and infected adults (RECAST study n = 1; PA-COVID study n = 18; median age = 67; mild n = 5, severe n = 7, severe with IFN autoantibodies n = 7). (G) UMAPs, showing the location of cells belonging to the respective group (colored in, whereas grey identifies cells in other groups). Grey outlines indicate cluster regions enriched in infected children or adults. Severe patients with IFN autoantibodies were excluded from the analysis. Patients with DPSO range from 1 to 19 days were included (median of 8.5, inf. children median = 4.5, inf. adult median = 10). (H) Dotplot, showing scaled average expression of genes in B cell clusters, increased with infection (4, 5, 6, 8 and 12). Clusters are split between infected children and infected adults. Severe patients with IFN autoantibodies were excluded from the analysis. A horizontal line splits the dotplot in two parts – genes above the line were curated based on the presence of clusters with pronounced ISG signature and include other genes useful for annotation, genes below the line were found to be differentially expressed between the clusters (FindMarkers Seurat function). Patient group color-coded figurines on the top point out cluster accumulation patterns as seen in S5E. (I) Volcano plot, showing the results of differential expression (DE) analysis of B cells from clusters expanded with infection (4, 5, 6, 8 and 12) using DESeq2. Significantly enriched genes (baseMean counts over 50 and adjusted p value < 0.05, a total of 482 genes), are colored green and 20 significant genes with highest absolute log2 fold change are labeled. Additional significant genes, tangential to other findings, were labelled ad-hoc. Positive log2 fold change values mean an enrichment in pediatric infected patients and negative – in infected adults. Severe patients with IFN autoantibodies were excluded from the analysis. (J) Stacked bar chart, showing proportions of cluster labels, given to the cells with callable BCR clonal identity that overlaps with at least one of the cells in the plasmablast population (cluster 12), calculated over all expanded B cell clusters. Severe patients with IFN autoantibodies were excluded from the analysis.

Furthermore, ISG^+^ non-classical monocytes were almost exclusively expanded in children with asymptomatic disease progression indicating that robust IFN responsiveness might lead to reduced clinical symptoms (Mendeley Figure 2). Increased proportions of HLA-DR^low^ classical monocyte scRNAseq clusters align very well with our CyTOF phenotypic characterization of monocytes (Figure 1D,E). HLA-DR^low^ monocyte scRNAseq cluster 4 transcribed high levels of inflammatory mediators such as S100A8/9/12 and the chemokines CXCL8 and CXCL2 (Figure 3C,D). Consequently, Gene Ontology (GO) gene set enrichment analysis (GSEA) of monocytes resulted in enrichment of “neutrophil mediated immunity” for adults (Figure S4F), which was accompanied by higher serum CXCL8 concentrations (Figure S4G). Serum samples from infected adults also contained significantly higher levels of IL-1b, IL-6 and TNF compared to those from infected children (Figure S4G). In contrast, the high *STAT1* and *STAT2* transcription within children-associated ISG^+^ monocyte clusters (Figure 3C & Figure S4D) translated into an overall increased expression of both genes in all monocytes of infected children (Figure 3D) indicative of enhanced type I IFN responsiveness. In summary, monocytes in infected children display signatures indicative of a higher IFN responsiveness and less inflammatory state relative to adults.

### Molecular characterization of SARS-CoV-2 infection induced T cells reveal age-dependent differences in STAT involvement and differentiation

To investigate whether altered type I IFN responsiveness and signaling mechanisms are also responsible for the age-dependent phenotypic differences in activated T cells, we performed subclustering of the TCRab^+^ T cell space from the PBMC scRNAseq data (Figure 4A, Figure S5A). Of the identified 21 T cell clusters we detected significantly higher proportions of ISG^+^ CD4^+^ and CD8^+^ T cell clusters 8, 17 and 18 in samples from infected children (Figure 4B,C, Figure S5B). In contrast, adults responded with expansion of *STAT3*^high^*SOCS3*^high^, *PD1*^high^ and proliferating T cell clusters 2, 7, 12 and 15 (Figure 4B,C, Figure S5B). Differential gene expression analysis revealed higher expression of *PRDM1*, *BCL3* and *SOCS3* in T cells from infected adults (Figure 4D). *PRDM1* encodes for the transcription factor BLIMP-1, which is known to counter-regulate T follicular helper cell differentiation.^29^ Together with increased CCR6 but low CXCR5 expression in CD4^+^ T cells as determined by CyTOF analysis these data hint towards an enhanced accumulation of peripheral T helper cells in adults.^30,31^ The difference in *SOCS3* expression was also intriguing. SOCS3 inhibits JAK activity through its kinase-inhibitory region thereby restricting *STAT1* activation and adequate type I IFN signaling.^32^ *SOCS3* is a down-stream target gene of *STAT3*.^33,34^ Apart from inducing *SOCS3* and thus inhibiting type I IFN signaling, *STAT3* also controls its own transcription in a positive feedback loop.^33,35^ Similarly, canonical type I IFN signaling via the IFN-stimulated gene factor 3 (ISGF3) complex promotes gene expression of *STAT1*, *STAT2* and *IRF9*.^34^ Thus, gene expression levels of *STAT1* and *STAT2* versus *STAT3* can serve as an indicator of canonical ISGF3-STAT1/2 mediated and gamma-activated sequence (GAS)-STAT3 mediated signaling, respectively. Consistent with this notion, only ISG^+^ T cells enriched in samples from infected children (clusters 8, 17 and 18) were characterized by high *STAT1* and *STAT2* transcription (Figure 4C). In T cells from adult-associated clusters 2 and 7 we also detected high *STAT3*, *SOCS1* and *SOCS3* transcription levels, which was even more pronounced for cells from infected adults (Figure 4C). In T cells from adult-associated clusters 2 and 7 we also detected high *STAT3*, *SOCS1* and *SOCS3* transcription levels, which were even more pronounced for cells from infected adults (Figure 4C). Additionally, *BCL3* is a known positive regulator of *STAT3* and is associated with active, phosphorylated has been associated with the presence of its phosphorylated form p-STAT3.^36,37^ Similar to monocytes, before we also observed enrichment of proinflammatory *S100A8* and *S100A9* genes in TCRab^+^ T cells of infected adults, indicative inflammatory signaling. The selective increase in ISG^+^ T cells in samples from infected children was further confirmed by gene set enrichment analysis (Figure 4E). We next compared average scaled expression in all infection-induced TCRab^+^ T cells (Figure S5C). Indeed, T cells displayed higher *STAT3* transcription in adults, when compared to their counterparts in children. Similarly, *SOCS1* and *SOCS3* were elevated in T cells from infected adults, wheres *STAT1* and *STAT2* transcript levels were increased in infection-induced TCRab^+^ T cells in infected children.

In summary, T cells from children and adults infected with SARS-CoV-2 were characterized by dichotomous expression of STAT molecules and their downstream target genes. In children, a prominent ISG profile was associated with high levels of *STAT1* and *STAT2* transcription, whereas elevated levels of *STAT3*, *SOCS1* and *SOCS3* were found in T cells from adults.

### Divergent molecular profiles of infection-induced B cells translate into different plasmablast differentiation programs

To understand whether also the disparate B cell phenotypes could be aligned to differences in type I IFN responsiveness, we analyzed gene transcription in CD19^+^ B cells in our single cell RNAseq data (Figure 4 F-J; S5D,E). We identified 13 B cell clusters, which we merged into 8 clusters, based on transcriptional similarity (Figure 4F-H, Figure S5D,E).

In agreement with the CyTOF data, we detected an increase of highly CD69-expressing B cells in samples of infected adults (cluster 6, *CD69*^high^; Figure 4F,G and Figure S5D,E). B cells belonging to that cluster displayed some characteristics of DN B cells, that are negative for *CD27* and *CR2*, which encodes for CD21, (Figure 4H, S5D). B cells did still transcribe low levels of *IGHD* and *CXCR5*. Given the lower input and fewer clusters, cluster 6 likely resembled a mixture of the CD69-expressing GCB-like cells and CD69-expressing DN-like B cells that we detected by CyTOF (clusters 14 and 1, respectively) (Figure S3A).

Importantly, high *CD69* transcription was observed in infection-induced B cells in adults but not in children (Figure F4I). In turn, B cells increased in infected children transcribed genes characteristic for transitional B cells (cluster 5 *CD24*^+^*MME*^+^*CD38*^+^*CR2*^+^) and IFN-responsive naïve B cells (cluster 4 *IGHD*^+^*IGHM*^+^*CR2*^+^*ISG*^+^). This was in line with the increased proportions of the CD10^high^CD38^high^TIGIT^-^ transitional-like B cells (cluster 3) and IgD^+^CD21^+^ naïve-like B cells (cluster 5) in the CyTOF data analysis. Importantly, B cells belonging to the scRNAseq cluster 4 (GCB-like naïve, ISG^+^) also transcribed high levels of *CXCR5*. Most likely the cluster also contained GCB-like cells, corresponding to IgD^+^CD21^+^CXCR5^high^ GCB cells (cluster 11) in the CyTOF data. One additional cross-omics characteristic of infection-induced B cells in children was the high expression of *ILRA* encoding for CD123, which was not detected in any adult-specific subset. Plasmablast frequencies were similar in infected adults and children.

A more detailed comparison of B cells from infected children and adults accumulating in the clusters described above revealed further important transcriptional differences (Figure 4H). First, the ISG signature of B cell cluster 4 was much more pronounced in samples of infected children, whereas expression of lead genes (*CD83*, *CD69* and *TNFRSF13C*, encoding for the BAFF receptor) within DN-like, activated, *CD69*^high^ B cells (cluster 6) was increased in adults. Expression of *CD83* and *CD69* was higher in adults compared to children when analyzing the total number of B cells contained in all infection-induced clusters (Figure 4I) highlighting the concordance between the CyTOF (Figure S3F) and scRNAseq data. Plasmablasts from infected adults showed little *ISG15* expression but prominent *SOCS3* transcription (Figure 4H). Those transcriptional differences between plasmablasts of infected children and adults could indicate diverging differentiation routes. Indeed, analyzing the clonal overlap of B cell receptors between infection-induced B cell populations and plasmablasts revealed that children exclusively generated plasmablasts from transitional and ISG^+^ B cells, whereas both DN and DN-like, activated, CD69^high^ B cell populations in addition to ISG^+^ B cells contributed to the plasmablast pool in infected adults (Figure 4J). This might explain the enhanced *SOCS3* transcription which is known to be increased in CD21^-^ DN B cells due to *STAT3* overactivation.^38^

In summary, disparate type I IFN responsiveness may contribute to the shift from predominantly GCB responses to activation of both GCB and extrafollicular B cells as a source of plasmablasts in older individuals.

### Consequences for local T cell activation and antibody profiles

To establish a link between the observed systemic age-dependent differences and local responses, we reanalyzed T cells from our previously published nasal swab scRNAseq data from samples of infected patients and controls.^8^ Subclustering of TCRab^+^ T cells resulted in 8 main clusters (Figure S6A). From those, T cell cluster 6 was only enriched in infected children, whereas clusters 0, 4, 5 and 7 were enriched in both infected children and infected adults (Figure 5A, Figure S6B). Clusters 0 and 7 were ISG^+^ CD8^+^ and CD4^+^ T cell clusters, respectively (Figure 5A, S6A). Transcription of IFN response genes and especially of *CD38* was higher in cluster 7 T cells from infected children in comparison to infected adults (Figure 5A). In contrast, cluster 0 and 4 T cells from infected adults showed increased *TNF* gene expression. Consequently, plotting mean scaled expression of *CD38* and *TNF* in infection-induced expanded clusters against age revealed significant correlations. Similar to the systemic response shown before (Figure 2D) we detected a negative correlation of *CD38* expression in mucosal T cells with age, whereas pro-inflammatory *TNF* transcription gradually increased with age (Figure 5B).

**Figure 5.**
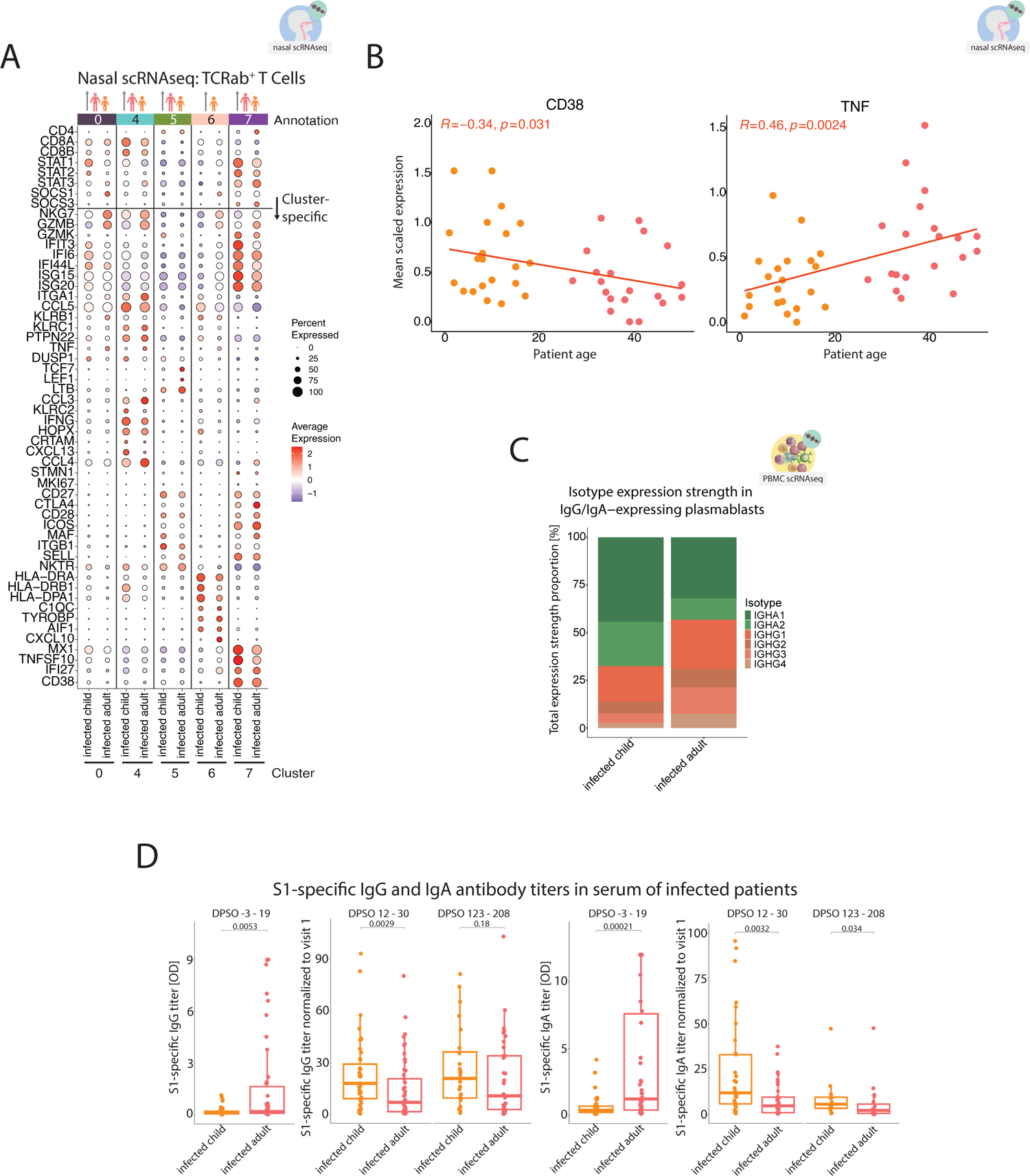
Consequences for local T cell responses and generated antibody profiles. (A) Dotplot, showing scaled average expression of genes in TCRab^+^ T cells, subset from the nasal swab scRNAseq data.^8^ Clusters, increased with infection (0, 4, 5, 6 and 7) are shown. A total of 8 clusters has been produced using a graph-based approach as implemented in Seurat package (KNN graph with Louvain community detection). Clusters are split between infected children (RECAST study n = 24; median age = 9; asymptomatic n = 2, symptomatic n = 22) and infected adults (RECAST study n = 21; median age = 39; mild n = 21). Patients with DPSO range from 0 to 12 days were included (inf. children mean = 4.5, inf. adult mean = 5). A horizontal line splits the dotplot in two parts – genes above the line were curated based on the presence of clusters with pronounced ISG signature and include other genes useful for annotation, genes below the line were found to be differentially expressed between the clusters (FindMarkers Seurat function). Patient group color-coded figurines on the top point out cluster accumulation patterns as seen in S6B. (B) Scatter plots showing CD38 and TNF genes transcription (average scaled expression in clusters 0, 4, 5, 6 and 7, expanded with infection) for each donor, plotted against donor’s age, using TCRab^+^ T cells, subset from nasal swab scRNAseq data.^8^ Only infected children (RECAST study n = 24; median age = 9; asymptomatic n = 2, symptomatic n = 22) and infected adults (RECAST study n = 21; median age = 39; mild n = 21) are shown. Patients with DPSO range from 0 to 12 days were included (inf. children mean = 4.5, inf. adult mean = 5). Linear models were fitted to the data points and Spearman’s rank correlation coefficients are shown. (C) Stacked bar chart showing relative expression strength of heavy chain genes encoding for the different IgG and IgA isotypes in plasmablasts (B cell cluster 12, PBMC scRNAseq experiment). Plasmablasts, expressing either of the heavy chain genes, were pre-selected. Expression values for each gene were calculated and normalized to the total expression of all heavy chain genes. Infected children (RECAST study n = 13; median age = 9; asymptomatic n = 5, symptomatic n = 8) and infected adults (RECAST study n = 1; PA-COVID study n = 11; median age = 63.5; mild n = 5, severe n = 7). (D) Box plots of S1-specific IgG (left) and IgA (right) antibody titers for the acute infection phase measurement (median DPSO = 6, inf. children median DPSO = 4, inf. adults median DPSO = 8) as well as follow-up measurements done approximately two weeks later (median DPSO = 21, inf. children median DPSO = 18, inf. adults median DPSO = 22) and six months later (median DPSO = 170, inf. children median DPSO = 146, inf. adults median DPSO = 170). Titers for second and third timepoint are normalized to the first timepoint for each patient (fold change, ratio). Infected children (RECAST study n = 52; median age = 8; asymptomatic n = 9; symptomatic n = 43) and infected adults (RECAST study n = 40, PA-COVID study n = 35; median age = 42; mild n = 55, severe n = 20) are shown. Wilcoxon; *, p < 0.05; **, p < 0.01; ***, p < 0.001; ****, p < 0.0001.

Next, we studied whether the diverse plasmablast differentiation programs were linked with age-dependent differences in antibody responses. Transcriptional analysis of Ig heavy chain genes in plasmablast revealed an increased relative expression of IgA isotypes in samples of children in comparison to those from adults, which had higher expression of IgG and especially complement-fixing IgG1 and IgG3 isotypes (Figure 5C).

Studying antibody responses in serum of infected patients, SARS-CoV-2 infected adults showed a swift anti-S1 IgA and IgG antibody response during the first two weeks after symptom onset (visit 1, DPSO −3–19; Figure 5D), which was significantly enhanced, compared to infected children. Although children responded with a delay, we detected a strong increase of IgA production at later time points (visits approximately two weeks after the first, DPSO 12-30 and approximately six months after the first, DPSO 123-208; Figure 5D).

### Age-dependent immune cell activation profiles are independent of comorbidities and conserved in a hamster infection model

Different risk factors have been established for COVID-19 severity.^39^ In order to exclude comorbidities as driving factors of the observed age-dependent phenotypes, we tested for confounding effects using linear regression modelling.

Using data from the CyTOF measurements, we displayed results from the CD4^+^ and B cell profiling of SARS-CoV-2-infected patients in a scatterplot (Figure S6C). Patient age was plotted against mean scaled marker expression values. We used Charlson Comorbidity Index (CCI, turquois fill for CCI >0) as a proven measure of patient comorbidity status. Obesity was defined as BMI > 30 (binary variable, triangle for BMI>30). We calculated linear regression models with the formulas “Expression ∼ Age” (black line) and “Expression ∼ Age + CCI score + obesity (0 or 1)” (turquoise line). Distance between the two lines thus represents the influence of the CCI score and obesity on the intercept (mean value of the response variable when all the predictor variables are zero). The values above each panel show the coefficient and the p value for the respective variable from the model output. CCI score was calculated excluding age, as age is already present in the model as a separate variable.

Importantly, the presence of comorbidities and/or obesity did not significantly alter the linear correlation between age and CD38 and CCR6 expression by CD4^+^ T cells or CXCR5 and CD69 expression by B cells, indicating that the differences in the immune response to infection that we identified in our study are indeed driven by age, and not a function of confounding comorbidities.

Importantly, patient stratification according to disease severity showed that even when comparing symptomatic children to mildly affected adults, the age-dependent patterns of infection-induced monocyte, CD4^+^ and B cell subsets remained (see Mendeley Figure 2). However, the samples from the severely affected patients showed a more pronounced phenotype.

To further substantiate our findings, we made use of existing samples from a SARS-CoV-2– infection model in hamsters. Similar to humans, hamsters show age-dependent severity of SARS-CoV-2 infection^40^. We had access to samples from infected young and old animals.^40^ We performed bulk RNA sequencing of whole blood samples from young (6 weeks) and old (32-34 weeks) infected hamsters as well as uninfected controls. Specifically, we investigated whether the gene sets identified in the human system can distinguish between samples from young and old infected hamsters. Indeed, young and old infected hamsters clustered separately. (Figure S6D) Whereas whole blood samples from young hamsters showed increased expression of ISGs *Isg15*, *Oas3*, *Ifit1* and *Irf7*, samples from old hamsters were characterized by higher transcription of *Il1b* and *S100* genes. This closely resembles the transcription profile of human monocytes and T cells (Figure 3C,D and Figure 4C,D). None of the transcripts were detected in samples of uninfected hamsters. These results confirm the age-dependent immune responses to SARS-CoV-2 infection in a well-established experimental system without potential confounding effects of comorbidities.

### Disturbed type I IFN responsiveness reduces formation of CD38^+^ CD4^+^ T cells and enhances cytokine production in older individuals

To confirm that the opposing T cell phenotypes in children and adults were linked to differential responsiveness to type I IFN stimulation, we set up *in vitro* cultures using PBMCs from uninfected children and adults (Figure 6A). PBMCs were stimulated with a superantigen (Staphylococcus enterotoxin B; SEB) as a TCR stimulus in the presence or absence of recombinant IFN alpha (IFNa), or CpG DNA-oligonucleotides that activate TLR-9 to induce strong IFNa production, mimicking virus-induced immune responses (CpG2216, Figure 6A).^41,42^ Addition of CpG to PBMC cultures increased CD38 expression on CD4^+^ T cells (Figure 6B). This was however only significant for children. Importantly, CpG-induced enhancement of CD38 expression on CD4^+^ T cells could be mimicked by recombinant IFNa and reversed via blocking of type-I IFN signaling by addition of type-I IFN decoy receptor (B18R; open reading frame in the Western Reserve strain of vaccinia virus) (Figure 6C). In contrast, T cells from children showed a lower CCR6 expression by CD4^+^ T cells compared to cells from adults upon TCR stimulation (Figure 6D). The addition of CpG further lowered CCR6 expression. The effect was stronger when adding IFNa. However, T cells of adults were resistant to IFNa-mediated downregulation of CCR6. Notably, addition of CpG elicited comparable IFNa production in PBMCs from children and adults, suggesting that different responsiveness rather than IFN production caused the altered phenotype (Figure 6E). This altered responsiveness to CpG in samples from children and adults was not due to differences in naïve and memory T cell composition, as both compartments from children but not adults were able to significantly enhance CD38 expression (Figure 6F).

**Figure 6.**
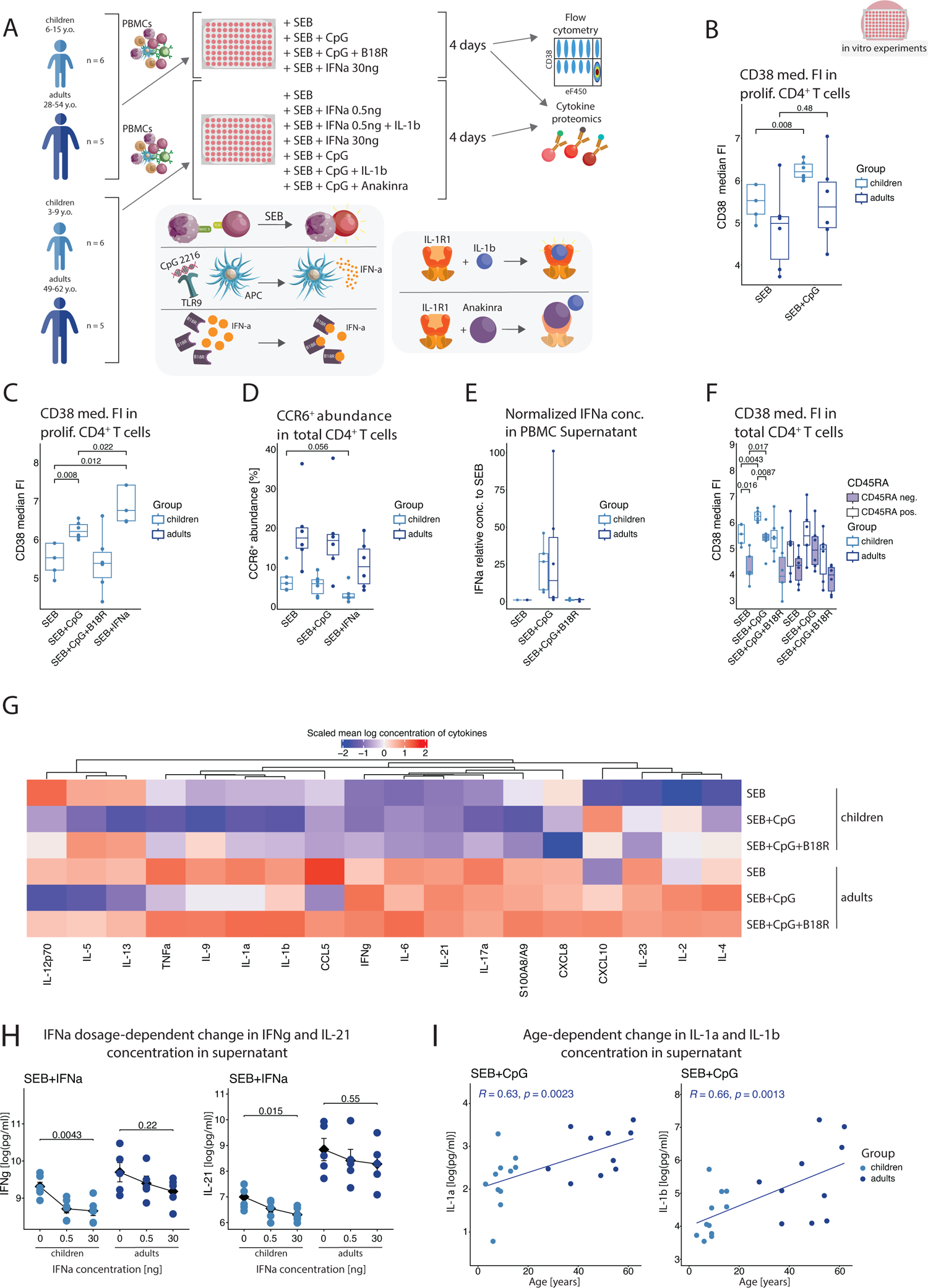
Mechanistic *in vitro* studies link age-dependent rewiring of type I IFN responsiveness with in vivo-detected opposite activation profiles (A) Overview of the workflow used to study the responsiveness to type I IFN and IL-1b. PBMCs from uninfected children and adults were stimulated with either SEB, or a combination of SEB, ODN CpG2216, B18R and recombinant IFN-a. Here, uninfected children (RECAST n = 6; median age = 11) and uninfected adults (RECAST n = 6; median age = 41) were included. In a parallel experiment series, different concentrations of IFN-a as well as combinations of SEB, ODN CpG2216, IL-1b and IL-1b inhibitor Anakinra were tested (uninfected children RECAST n = 6; median age = 8; uninfected adults RECAST n = 5; median age = 55). After 4 days of incubation, phenotypic differences in activation marker expression were determined by flow cytometry, while cell culture supernatants were used for cytokine and chemokine quantification. Experiments focused on IL-1b and Anakinra influence were only measured in cytokine proteomics. (B) Box plot of arcsinh-transformed median CD38 fluorescence intensity in proliferating CD4^+^ T cells, showing influence of CpG2216-mediated activation on CD38 expression. Uninfected children (RECAST n = 6; median age = 11) and uninfected adults (RECAST n = 6; median age = 41). Wilcoxon test p values shown for chosen comparisons. Dropout in uninfected children group SEB condition is due to low cell number, not allowing for inclusion of all activation conditions. (C) Box plot of arcsinh-transformed median CD38 fluorescence intensity in proliferating CD4^+^ T cells, showing influence of CpG2216-medited activation and IFNa (30ng/ml) on CD38 expression in children. Uninfected children (RECAST n = 6; median age = 11) and uninfected adults (RECAST n = 6; median age = 41). Wilcoxon test p values shown for chosen comparisons. Dropouts in SEB and SEB+IFNa perturbations are due to low cell number, not allowing for inclusion of all activation conditions. (D) Box plot of the relative abundance of gated CCR6^+^ population in proliferating CD4^+^ T cells, showing influence of IFN (30ng/ml) on CD38 expression in children. Uninfected children (RECAST n = 6; median age = 11) and uninfected adults (RECAST n = 6; median age = 41). Wilcoxon test p values shown for chosen comparisons. Dropout in uninfected children, SEB perturbation is due to low cell number, not allowing for inclusion of all activation conditions. (E) Box plot of IFNa concentration measured in cell culture supernatant and normalized to values detected in SEB condition for each patient, showing the effectiveness of CpG2216 in provoking IFNa release as well as of B18R in reducing the concentration of soluble IFNa. Uninfected children (RECAST n = 6; median age = 11) and uninfected adults (RECAST n = 6; median age = 41). Dropout in uninfected children is due to low cell number, not allowing for inclusion of all activation conditions. (F) Box plots of CD38 median signal intensity in proliferating CD4^+^ T cells, separated into CD45RA^-^ (violet filling) memory and CD45RA^+^ naive subpopulations, showing the difference in CD38 upregulation in response to CpG2216-mediated activation and IFNa release between memory and naïve CD4^+^ T cells. Uninfected children (RECAST n = 6; median age = 11) and uninfected adults (RECAST n = 6; median age = 41). Wilcoxon test p values shown for chosen comparisons. Dropout in uninfected children, SEB perturbation is due to low cell number, not allowing for inclusion of all activation conditions. (G) Heatmap, showing scaled average log concentration of the 18 cytokines measured in co-culturing experiments for different perturbations using PBMC cells from uninfected children (RECAST study n = 6; median age = 8) and uninfected adults (RECAST study n = 5; median age = 55). (H) Scatter plots, showing the dependence of IFNg and IL-21 concentrations on the IFNa concentration. Uninfected children (RECAST study n = 6; median age = 8) and uninfected adults (RECAST study n = 5; median age = 55). P values represent the result of testing with non-parametric Wilcoxon test. (I) Scatter plots, illustrating the correlation between donor age and IL-1a and IL-1b concentrations in supernatant when PBMCs are stimulated with SEB and CpG. Correlation statistics are presented in the upper left corner of each plot. uninfected children (RECAST study n = 11; median age = 9) and uninfected adults (RECAST study n = 10; median age = 55).

In summary, we observed an age-dependent resistance of activated T cells to respond to exogenous or endogenous type-I IFN. CD38 acts as an inhibitor of T cell activation by consuming NAD.^43^ CD38 enzymatic function was shown to increase Treg activity, while inhibiting T cell cytotoxicity and Th1/Th17 population.^44–46^ In contrast, CCR6 promotes inflammatory responses by guiding effector T cells into peripheral tissues such as the lung.^30^ Additionally, type I IFN can regulate production of inflammatory cytokines by T cells, particularly IL-17A and IL-21.^47–49^ Thus, the divergent responsiveness to type I IFN between pediatric and adult immune cells could also be reflected in differential control of inflammatory cytokine production. Therefore, we tested whether addition of CpG to SEB stimulation of cells from children and adults modulated inflammatory cytokine production. We performed a multiplex analysis of 18 cytokines, chemokines, and S100A8/A9, followed by unsupervised clustering analysis for all conditions and mediators tested (Figure 6G). Four main clusters were identified.

The first cluster encompassed IL-2, IL-4, IL-23 and CXCL10, which were increased upon addition of CpG to SEB in cultures of cells from both age groups.

The second cluster included IL-12p70, IL-5 and IL-13, which were produced in equal amounts by SEB-stimulated pediatric and adult cells and inhibited by the addition of CpG in both age groups. This inhibitory effect was dependent on type I IFN as cytokine production could be restored by B18R.

The third cluster comprised IL-17A, IL-6, S100A8/A9, IFN gamma (IFNg), IL-21, and CXCL8 was also reduced u by addition of CpG to stimulated pediatric and adult cell cultures. However, the concentration of these mediators was slightly higher in supernatants of cultures from adults compared to children (IL-17A p=0.015, IL-6 p=0.049, S100A8/A9 0.041, IFNg p=0.066, IL-21 p=0.11, CXCL8 p=0.27).

Interestingly, these mediators showed a different reactivity to CpG in cell cultures from adults versus children. While IFNg and IL-21 levels remained unchanged in pediatric cultures, addition of CpG increased the release of both IFNg and IL-21 in PBMC derived from adult donors. Conversely, CpG reduced S100A8/A9 and CXCL8 release by pediatric PBMCs, which was not the case in PBMCs from adults. Similar to the CD38 and CCR6 data (Figure 6B-D,F), this points to a differential type I IFN responsiveness. This was further underscored by a clear dose-response relationship with regards to reduced IFNg and IL-21 in cells from children but not in adult cells (Figure 6H).

The fourth cluster comprised IL-1b, IL-1a and TNF, which showed higher concentrations in cultures of adult cells (IL-1b p=0.006, IL-1a p=0.012, IL-9 p=0.049, TNF p=0.066). Consequently, we detected a positive correlation between age and IL-1a or IL-1b (Figure 6I) as well as TNF (R=0.47, p=0.032) and IL-9 (R=0.45, p=0.038). Of note, IL-1 and type I IFN are known to antagonize one another.^50,51^

To test the contribution of IL-1 in interfering with type I IFN-mediated inhibition of cytokine production by T cells, we added IL-1 along with type I IFN to the culture system (Figure 6A, lower cohort). Addition of type I IFN decreased the production of IL-6, IL-9 and IL-17A (Figure S7), while adding IL-1b along with type I IFN restored the secretion. Similarly, neutralizing IL-1 with the soluble IL-1 receptor antagonist (Anakinra) in SEB & CpG stimulated cultures reduced IL-17A production, whereas adding IL-1b to SEB & CpG cultures further enhanced IL-6, IL-9 and IL-17A secretion. Similar counterregulatory function of IL-1 was not observed for other cytokines such as IL-5.

Importantly, the higher *in vitro* production of inflammatory mediators by adult immune cells is in line with our *in vivo* findings of increased IL-1b, IL-6, IL-8 (CXCL8), TNF concentrations in serum samples of infected adults (Figure S4G) as well as increased S100A8/A9 transcription in adult monocytes and CD4^+^ T cells (Figure 3D & Figure 4D) and TNF transcription in nasal swab T cells (Figure 5B). Thus, increased production of inflammatory mediators may be due to an altered reactivity to type I IFN (e.g. IFNg, IL-21) and counter regulation by IL-1 (IL-6, IL-9, IL-17A).

### Enhanced STAT3 activation characterizes IFNa signaling in T and B cells during aging

Our transcriptional analyses indicated a shift from a predominant STAT1 to STAT3 activity in T and B cells from SARS-CoV-2 infected adults. To confirm the altered balance, we determined the amount of phosphorylated STAT1 (pSTAT1) and STAT3 (pSTAT3) by flow cytometry in the *in vitro* culture system (Figure 7A, B). SEB stimulation alone did not lead to an altered balance of STAT1 and STAT3 activation in CD4^+^ T and B cells from children and adults (Figure 7B). Addition of IFNa, lead to an increase in the pSTAT1 to pSTAT3 ratio in PBMC cultures from children. In contrast, adult cells responded to IFNa stimulation with enhanced STAT3 activation maintaining a lower pSTAT1/pSTAT3 ratio.

**Figure 7.**
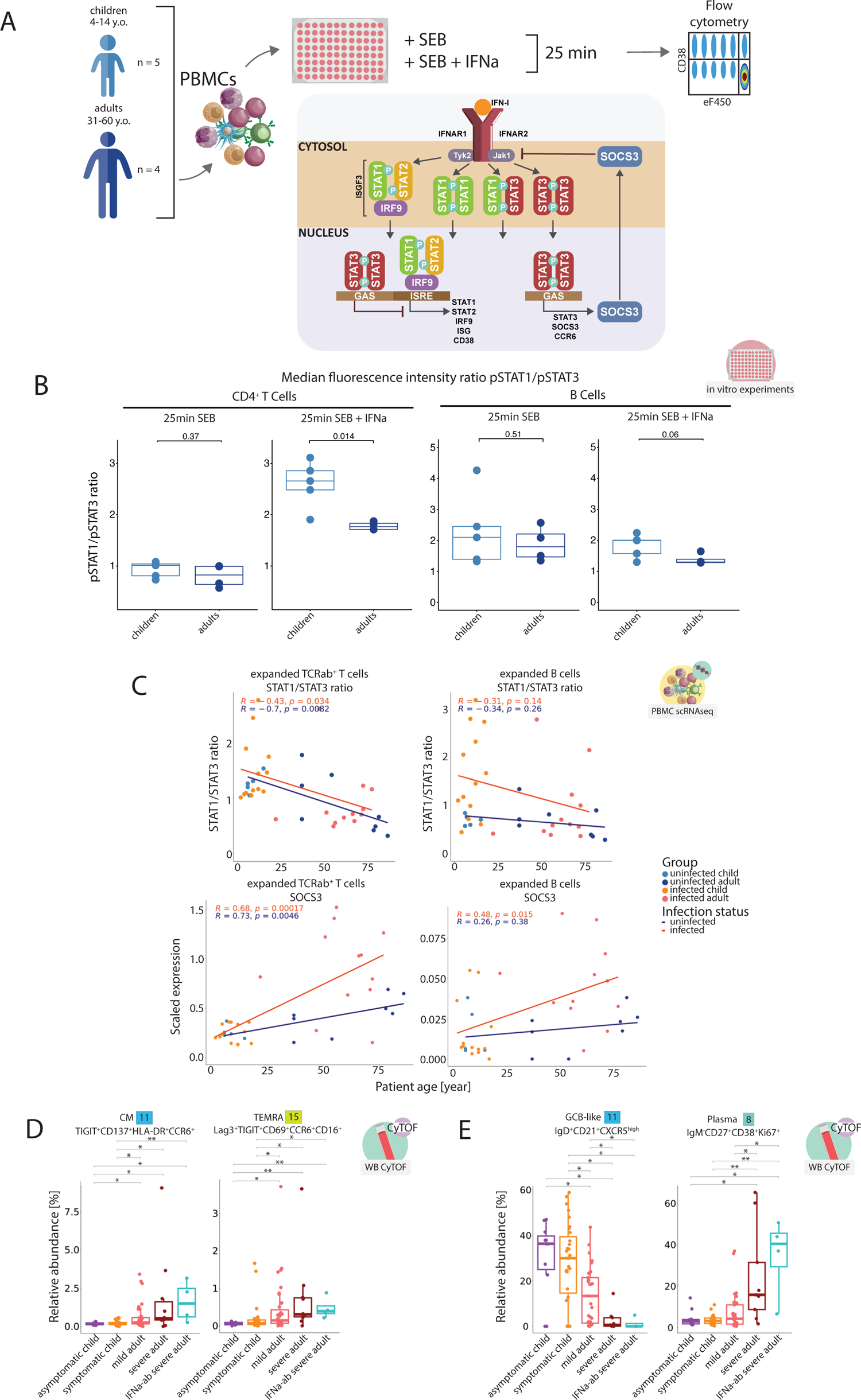
Altered type I IFN signaling and gradual involvement of STAT3 activation in stimulated T and B cells of older individuals. (A) Overview of the workflow studying the phosphorylation dynamics of STAT1 and STAT3 in T cells and B cells from children and adults. PBMCs from control children and adults were stimulated with either SEB, or a combination of SEB and recombinant IFN-a. After 25 minutes of incubation levels of phosphorylated STAT1 and STAT3 were determined by flow cytometry for pre-gated populations. Uninfected children, RECAST study n = 5; median age = 9; uninfected adult, RECAST study n = 4; median age = 53.5. (B) Box plots, showing the ratio of pSTAT1 to pSTAT3 (based on measured median fluorescence intensity) for CD4^+^ T cells and B cells of control children and adults following 25-minute incubation with SEB or SEB and IFNa. Uninfected children, RECAST study n = 5; median age = 9; uninfected adult, RECAST study n = 4; median age = 53.5. Wilcoxon p value shown. (C) Scatter plots of STAT1/STAT3 transcription ratio and SOCS3 transcription (average scaled expression in clusters expanded with infection) for each donor, plotted against donoŕs age, using PBMC scRNAseq experiment data. CD4^+^ T cells were a subset of the total TCRab^+^ pool using CD8A and CD8B genes (assigned CD4 flag, if cell is negative for both). Uninfected children (RECAST study n = 5; median age = 7), uninfected adults (RECAST study n = 4, EICOV/COVIMMUNIZE study n = 4; median age = 66), infected children (RECAST study n = 13; median age = 9; asymptomatic n = 5, symptomatic n = 8) and infected adults (RECAST study n = 1; PA-COVID study n = 11; median age = 67; mild n = 5, severe n = 7). Severe patients with IFNa autoantibodies were not used in this analysis. Linear models were fitted to the data points and Spearman’s rank correlation coefficients are shown. (D) Box plots showing relative abundance of infection-induced clusters 11 and 15 from the FlowSOM algorithm, calculated per sample within all CD4^+^ T cells from the CyTOF data. Only samples measured during the acute phase of infection (defined as first 14 days after symptom onset) are shown. Patients with anti-IFN autoantibodies are included as a separate group. Group composition as follows: uninfected children (RECAST n = 13; median age = 7), uninfected adults (RECAST n = 25; median age = 48), infected children (RECAST study n = 48; median age = 8; asymptomatic n = 11, symptomatic n = 37) and adults (RECAST n = 21, PA-COVID study n = 30; median age = 47; mild n = 37, severe n = 10, severe with IFN autoantibodies n = 4). Kruskal-Wallis + Wilcoxon; *, p < 0.05; **, p < 0.01; ***, p < 0.001; ****, p < 0.0001. (E) Box plots, showing relative abundance of infection-induced clusters 8 and 11 resulting from the FlowSOM algorithm, calculated per sample within all B cells from the CyTOF data. Only samples measured during the acute phase of infection (defined as first 14 days after symptom onset) are shown. Severe patients with IFN autoantibodies are included as a separate group. Group composition is thus as follows: uninfected children (RECAST n = 13; median age = 7), uninfected adults (RECAST n = 25; median age = 48), infected children (RECAST study n = 48; median age = 8; asymptomatic n = 11, symptomatic n = 37) and adults (RECAST n = 21, PA-COVID study n = 30; median age = 47; mild n = 37, severe n = 10, severe with IFN autoantibodies n = 4). Kruskal-Wallis + Wilcoxon; *, p < 0.05; **, p < 0.01; ***, p < 0.001; ****, p < 0.0001.

Interestingly, the differences in the balance between STAT1 and STAT3 involvement were also apparent *ex vivo* as we observed an age-dependent gradual decline in the ratio of *STAT1*/*STAT3* transcription for infection-induced (expanded) CD4^+^ T cells as well as B cell populations (Figure 7C). Consequently, *SOCS3* as a STAT3 target gene, followed an opposite pattern.

To functionally validate the importance of type I IFN signaling for the disparate phenotypes of CD4^+^ T and B cells during SARS-CoV-2 infection, we investigated samples from individuals with neutralizing antibodies to type I IFN, which we had previously identified (Figure 7D,E). Indeed, both CD4^+^ T cells and B cells from patients with autoantibodies to IFN showed the an “aged” phenotype, characterized by the highest median abundance of both adult-specific CD4^+^ T cell populations (CyTOF cluster 11 TIGIT^+^CD137^+^HLA-DR^+^CCR6^+^ CM and cluster 15 Lag3^+^TIGIT^+^CD69^+^CCR6^+^CD16^+^ TEMRA), lowest median abundance of GCB-like cells (CyTOF cluster 11 IgD^+^CD21^+^CXCR5^high^) and highest proportion of plasmablasts in total B cells (CyTOF cluster 8 IgM^-^CD27^+^CD38^+^Ki67^+^) among the infected patient samples.

In summary, the altered T and B cell phenotypes in samples from infected adults could be linked to a gradual loss of canonical type I IFN signaling via STAT1 and a shift towards STAT3 signaling.

## Discussion

Different clinical manifestations of viral respiratory infection in children and adults, and particularly in elderly, are well-known, but so far only poorly understood.^10^ Before the emergence of SARS-CoV-2, comparative immune response analyses to common respiratory viral infections across various age groups were difficult to interpret, due to differential pre-existing immunity caused by previous exposures in adults. We took advantage of our RECAST cohort to systematically compare innate and adaptive immune responses in children and adults to the same primary infection.^1^ Our study demonstrates distinct immune cell profiles and a switch from STAT1/STAT2-mediated antiviral ISG responses to a STAT3-driven inflammatory response across all examined immune cell types.

Together with direct *in vitro* demonstration of the opposing immune response patterns of children and adults, our findings reveal an age-associated rewiring of type I IFN signaling with gradual shift from canonical IFN signaling to proinflammatory STAT3 driven responses. This effect was noticeable after adolescence, but was most prominent beyond 55 years of age, in line with the increased risk of severe disease from SARS-CoV-2 infection.

At a younger age, strong ISG induction was associated with increased frequencies of HLA-DR^high^ monocytes and follicular T helper and B cell responses, favoring differentiation of IgA-expressing plasmablasts. In contrast, during aging we observed a continuous replacement of this network by HLA-DR^low^, CXCL8- and S100A8/A9-expressing monocytes, CCR6^+^CD69^+^ peripheral T helper cells as well as CD69^high^ follicular and atypical B cells, leading to a more rapid antibody response dominated by complement fixing IgG antibodies.

Our study confirms previous observations of an aging-induced reduction of circulating pDCs, the main cells that produce type I IFN in the immune system.^52^ However, a broad range of cells is capable of producing type I IFN including other immune cells, epithelial, and stromal cells. In line with this, our *ex vivo* and *in vitro* data showed that IFNa production is not significantly diminished in older patients, indicating that IFN production may not explain the divergent immune response patterns. We therefore investigated whether differential responsiveness to type I IFN and an altered IFN signaling might account for the age-dependent changes in immune responses to SARS-CoV-2.

Binding of type I IFN to the heterodimeric IFN receptor (IFNAR) complex activates the receptor-associated protein tyrosine kinases Janus kinase 1 (JAK1) and tyrosine kinase 2 (TYK2). In the canonical pathway, activated JAK1 and TYK2 phosphorylate STAT1 and STAT2 leading to their dimerization, nuclear translocation, and binding to IRF9 to form the ISGF3 complex with subsequent ISG transcription.^53^ Type I IFN signaling is not limited to this canonical pathway as activation of other STATs such as STAT3 is also possible.^53^ Importantly, STAT3 is not just activated in addition to STAT1 and STAT2 but rather fine-tunes type I IFN responses by negatively regulating the canonical pathway through e.g. sequestration of STAT1 or induction of negative transcriptional regulators like SOCS3.^32–35^ In addition, enhanced expression of other negative regulators such as SOCS1 (Figure 4C) may contribute to the altered type I IFN responsiveness of adult immune cells.^54,55^ Thus, the balance between STAT1/STAT2 versus STAT3 engagement determines the signaling output of type I IFN signaling. Our results revealed diminished infection-induced STAT1 activation in immune cells from older individuals. Indeed, T cells from older adults fail to exclude SHP-1 from the type I IFN-induced JAK/STAT signaling complex to ensure sustained STAT1 activation.^56^ Conversely, aging-associated enhanced STAT3 activation in CD8^+^ T cells interferes with a protective antiviral response and causes severe SIV infections.^57^ Interestingly, early during the pandemic Matsuyama and colleagues postulated that aberrant STAT signaling is central to COVID-19 pathology.^58^ Here, we provide evidence that this is not limited to infected cells but that IFN signaling deviation via STAT3 expands to cells of the innate and adaptive immune system in an age-dependent manner.

Apart from intracellular modulators, extracellular signals can interfere with type I IFN signaling. Interleukin 1 and type I IFN counter-regulate each other.^50^ We show that SARS-CoV-2 infection elicits higher IL-1b serum concentrations in adults (Figure S4G), which was positively correlated with donor age in PBMC cultures, (Figure 6I). We also demonstrated a synergistic activity of IFNa and IL-1b resulting in enhanced secretion of proinflammatory cytokines IL-6, IL-9 and IL-17A (Figure S7).

Along with enhanced STAT3 activation, we demonstrated significant transcriptional alterations in monocytes, CD4^+^ T cells and B cells during aging. SARS-CoV-2-induced monocytes populations in adults showed low HLA-DR expression (Figure 1D,E, Figure S1A,B). STAT3 negatively regulates MHCII expression^34^, whereas adequate type I IFN signaling promotes up-regulation of MHCII and costimulatory molecules on DCs and monocytes, characteristic for SARS-CoV-2 induced populations in children.^59,60^ This is in line with our previous findings which showed that severe COVID-19 is associated with increased numbers of HLA-DR^low^ proinflammatory monocytes.^17^ The present study positions those results into an age- and type I IFN signaling-dependent context. Furthermore, other features of adult-specific monocytes such as transcription of *CXCL8*, *CXCL2*, *S100A8/A9/12* are known downstream targets of STAT3, as per ENCODE database and literature sources.^61–64^

Our additional transcriptional analyses of blood samples from a SARS-CoV-2 infection hamster model confirmed the age-dependent shift from enhanced ISG transcription to *Ili1b* and *S100* gene induction during ageing (Figure S6D). Thus, our data do not confirm the age-dependent increase in the expression of members of the S100 protein family but relate this to qualitative changes in the immune response during an acute viral infection.^65^

The variable engagement of STAT1/STAT2 and STAT3 in CD4^+^ T cells was associated with divergent CD38 and CCR6 expression (Figure 2D, Figure 5B). CD38 acts as an inhibitor of T cell activation by consuming NAD.^43^ CD38 enzymatic function was shown to increase Treg activity, while inhibiting T cell inflammatory potential and Th1/Th17 population.^44–46^ This might explain our findings of low CCR6, *S100,* and *TNF* gene expression by children-specific SARS-CoV-2-induced T cells (Figure 2D, Figure 4D, Figure 5B). CD38 upregulation and activity in response to type I IFN may be one mechanism of immune response modulation to prevent hyperactivation and severe infections. One additional consequence of less inflammatory immune cell phenotypes in children was faster contraction of the response (Figure 2I,J).

STAT3 overactivation favors differentiation of CD21^low^CCR6^+^ DN/atypical extrafollicular B cells, which we identified as a central feature of SARS-CoV-2 infection of adults (Figure 2G, Figure S5E).^38^ Extrafollicular responses have been previously linked to severe COVID-19, but they have not been previously linked to age-induced deviation from canonical IFN signaling to STAT3-driven responses.^23^ Extrafollicular B cell responses are promoted by peripheral T helper cells which were also increased in infected adults.^23,30,66,67^

An effect of the divergent plasmablast maturation paths was the preference for IgA antibody isotypes in infected children and lower levels of genes encoding for complement-fixing IgG isotypes in comparison to plasmablasts from adults (Figure 5C). This would result in reduced formation of immune complexes and activation of the classical complement cascade. Hyperactivation of the complement system has long been implicated in COVID-19 severity and mortality and reduced production of complement-fixing IgG isotypes in the early phase seems to be one of the resilience strategies of infected children.^68,69^ This also ties in with our previous findings, where we could show a direct link between complement activation and T cell-driven immunopathology in adult COVID-19 patients.^18^

Taken together, we show that age-induced rewiring of type I IFN signaling causes qualitative alterations in immune cell activation. In children the STAT1/STAT2-dominated cell signaling elicited a protective immune response without the pathological features observed in adults. Increasing involvement of STAT3 activation during aging is linked to differentiation of hyperinflammatory innate and adaptive immune cells. Our data therefore provide molecular explanations for inflammation-prone responses to infections in older individuals. As the signaling conversion was also observed upon *in vitro* stimulation of cells from controls, as well as in a SARS-CoV-2 hamster infection model, our findings constitute a general phenomenon. It will be interesting to study whether the conversion of type I IFN signaling is linked to variable vaccination outcomes and should be considered when designing age-adapted regimens.

### Limitations of the study

We performed an observational cohort study to compare immune responses of children and adults to acute SARS-CoV-2 infection. The nature of observational studies harbors several limitations, particularly with regards to potential confounders and investigation of causality. In contrast to animal experiments or interventional, randomized trials, observational studies carry the risk of confounding as observed immunological mechanisms and clinical outcomes cannot be directly linked. However, knowing the regular limitations of cohort studies, we went at great lengths to minimize confounding factors and to achieve best-as-possible comparability between cohorts. A subset of children and adults were recruited from the same households, which should ensure largely similar exposure and environmental factors. We also excluded the possibility of comorbidity and obesity status-driven confounders by specifically modeling their influence on the core findings.

In addition to technical limitations due to the study design, there are limitations to the generalizability of our results, since other viral infections occur largely in immune experienced adults, which will influence the immune responses and profiles, compared to the infections in naïve hosts, investigated here.

## Methods

### Resource availability

#### Lead contact

Further information and requests for resources and reagents should be directed to the lead contact, Birgit Sawitzki.

#### Materials availability

This study did not generate new unique reagents.

#### Cohort 1 / Berlin Pa-COVID-19 cohort

Pa-COVID-19 is a prospective observational cohort study assessing pathophysiology and clinical characteristics of patients with COVID-19 at Charité Universitätsmedizin Berlin.^19^ It is being carried out with the approval of the Institutional Review board of Charité (EA2/066/20). Written informed consent was provided by all patients or legal representatives for participation in the study. This cohort includes 50 (+8 patients with neutralizing IFN-I autoantibodies) COVID-19 infected adult patients. All COVID-19 patients were tested positive for SARS-CoV-2 RNA in nasopharyngeal swabs and allocated into mild (WHO 2-4) or severe (5-7) disease groups according to the WHO clinical ordinal scale. Please refer to Table S1 for a list of patient samples included in different experiments, including information about days post symptom onset, age and sex compositions of the cohort. Table S2 provides an overview of the most important cohort statistics.

#### Cohort 2 / Berlin RECAST cohort

RECAST is a subproject of the Pa-COVID-19 observational study, aiming to characterize pathophysiology and SARS-CoV-2 infection progression primarily in patients under 18 years old.^1^ Data was collected from both minors and their family members in a longitudinal manner at three time points - directly after the diagnosis and at follow-up visits after approximately 2 weeks and 6 months. This cohort includes 40 SARS-CoV-2 infected-adults, 58 infected children, 35 control adult donors and 22 non-infected pediatric controls. None of the infected patients were hospitalized and thus fell under the WHO clinical ordinal scale of <3. Patients with either non-variant of concern (VOC) or Alpha (B.1.1.7) variant of SARS-CoV-2 were included in the analysis. Please refer to Table S1 for a list of patient samples included in different experiments, including information about days post symptom onset, age and sex compositions of the cohort. Table S2 provides an overview of the most important cohort statistics.

#### Age-matched elderly control cohort

Whole blood and PBMC samples of control donors over 50 (n = 14), that were included in CyTOF, scRNAseq and serum proteomics were obtained under protocols approved by the ethics committee of Charite – Universitatsmedizin Berlin {EA4/245/20 and EA4/244/20; EICOV and COVIMMUNIZE).^20^ Please refer to Table S1 for a list of patient samples included in different experiments, including information about days post symptom onset, age and sex compositions of the cohort. Table S2 provides an overview of the most important cohort statistics.

#### Control group definition

Control samples were defined as stemming from donors with a negative SARS-CoV-2 PCR test result without clinical signs of an ongoing infection.

## Method details

### Antibodies used for mass cytometry (cohort 1 & 2)

All anti-human antibodies pre-conjugated to metal isotopes were obtained from Fluidigm Corporation (San Francisco, USA). All remaining antibodies were obtained from the indicated companies as purified antibodies and in-house conjugation was done using the MaxPar X8 labeling kit (Fluidigm, USA). Antibodies are listed in the key resources table.

### Sample processing, antigen staining, and data analysis of mass cytometry-based immune cell profiling (cohort 1 & 2)

Sample processing, cell staining and acquisition was done as previously described.^17^ OMIQ.ai cloud-based cytometry analysis software was used for de-barcoding of individual samples, manual gating of singlets and removal of normalization beads and dead cells. Per-channel intensity ranges were aligned between batches using a reference sample - a replicate acquired across all batches, and a proprietary script, based on BatchAdjust function, to compute scaling factors at the event percentiles of choice on per-channel basis.^70^ Populations of interest, such as T cells (defined as CD3^+^CD19^-^, HLA-DR^-/+^CD11c^-/+^, CD14^-^CD15^-^ cells and separated into CD4^+^TCRgd^-^ and CD8^+^TCRgd^-^ populations), B cells (defined as CD3^-^CD19^+^ and CD14^-^CD15^-^ cells) and monocyte-dendritic cell (DC) space (defined as CD3^-^CD19^-^, CD56^-^ and CD14^+^HLA-DR^-/+^ cells), were manually pre-gated (in an approach, similar to our previous project) and subset for further analysis in the R programming environment.^18^

The individual immune populations were then transformed using the inverse hyperbolic sine function (asinh) and z-score normalized per-marker across all samples and all events. Data sets were clustered using FlowSOM algorithm as implemented in cytokfit R package (v. 0.99.0), setting the number of resulting clusters *k* as 30 (for CD4^+^ T cells) and 25 (for Monocyte-DC and B cell data sets).^71^ A pre-selection of markers has been used as basis for clustering for each immune population subset (T cells: CD62L, CD45RO, CD27, CD28, CD226, ICOS, PD1, Lag3, TIGIT, CD96, CD25, CD38, CD56, CD69, CD137, HLADR, Ki67, CXCR3, CXCR5, CCR6, CRTH2, CD161, KLRG1, KLRF1, CD10, CD11c, CD123, CD16, CD95, CD34; B cells: IgD, IgM, CD10, CD21, HLADR, CXCR5, CD27, CD38, CD25, CXCR3, CD69, Ki67, CD95, CD11c, CD137, CCR6, CRTH2, CD62L, CD226, ICOS, PD1, Lag3, TIGIT, CD96, CD123, KLRG1, KLRF1, CD16, CD28, CD45RO, CD56, CD161, CD34; monocytes and DCs: CD14, CD16, HLADR, CD11c, CD123, CD8, CD10, CD69, CD38, CD62L, CD25, Ki67, CD226, CD95, CCR6, CRTH2, CXCR3, CXCR5, ICOS, PD1, Lag3, TIGIT, CD96, CD56, KLRG1, KLRF1, CD27, CD28, CD45RO, CD137, CD161, CD34).

The resulting clusters were then manually merged in a pairwise manner, based on their similarity in z-normalized marker expressions, to correct overclustering. For CD4^+^ T cells, measurement timepoints beyond acute infection were included (approximately two weeks and six months after the first visit). As these samples were added post-hoc, KNN classification algorithm was used to assign new cells to the existing cluster classes, using clustering of acute samples as training set (R package “class” v.7.3-22, function knn, default parameters). UMAPs were calculated on the same pre-selected markers, using the R package “uwot” (version 0.1.14, n_neighbors = 20, min_dist = 0.1, Euclidean distance).^72^ The frequency of each cluster was calculated as the percentage of cells in each cluster for each patient and for each immune cell compartment. Statistical testing for the difference in the frequency of each cluster across severity groups was calculated with the adjusted Wilcox test (Benjamini-Hochberg) for clusters with significant Kruskal-Wallis test (adjusted p-value (Benjamini-Hochberg) < 0.05, adjusted across all clusters in each immune cell compartment). For cluster abundance-age and average signal-age scatterplots, a linear model was fitted using “geom_smooth” function (ggplot2 package, version 3.4.0). For some analyses, activated T cell clusters were pre-selected, defined as clusters having above average z-scored expression of activation markers (CD25, HLA-DR, CD38, CD137, CD69, and Ki67).

### Isolation of blood cells for scRNA-seq (cohort 1 & 2)

Human peripheral blood mononuclear cells (PBMCs) were isolated from heparinized whole blood by density gradient centrifugation over Pancoll (density: 1,077g /ml, PAN-Biotech, Germany). Subsequently, the cells were counted, frozen and stored in liquid nitrogen. On the day of the experiment, the frozen PBMCs were thawed in pre-warmed thawing medium (RPMI 1640, Gibco; 2% FCS, Sigma; 0,01% Pierce Universal Nuclease, Thermo Fisher, USA).

### 10x Genomics Chromium single-cell RNA-seq (cohort 1 & 2)

PBMCs were resuspended in staining buffer (DPBS, Gibco; 0,5% BSA, Miltenyi Biotec, Germany; 2 mM EDTA, Gibco, Thermo Fisher Scientific, USA} and hashtagged with 0.5 µg Total-Seq-C™ Hashtag antibodies for 30 min at 4°C. After the incubation, the PBMCs were washed three times, resuspended in DPBS, filtered through a 40 µm mesh (Flowmi™ Cell Strainer, Merck, Germany) and counted using the C-Chip hemocytometer (NanoEntek, South Korea). Subsequently, up to seven different samples were pooled equally. The cell suspension was super-loaded with 40000 - 50,000 cells per lane, in the Chromium™ Controller for partitioning single-cells into nanoliter-scale Gel Bead-In-Emulsions (GEMs).

In order to achieve a high enough cell number for each population on interest, the process above was repeated twice for each PBMC pool and additionally, B cells were enriched using untouched human B Cell Isolation Kit II (Miltenyi Biotec, Germany) and loaded separately with approximately 20000 cells per lane.

The Chromium Next GEM Single Cell 5’ v.2 Dual Index kit was used for reverse transcription, cDNA amplification and library construction of the gene expression libraries (10x Genomics, USA). For additional VDJ and hashtag libraries the Chromium Single Cell V(D)J Enrichment Kit, Human T Cell and Human B Cell (10x Genomics, USA), as well as the Chromium Single Cell 5’ Feature Barcode Library Kit (10x Genomics, USA) were used. All libraries were prepared following the detailed protocols provided by 10x Genomics, quantified by Qubit Flex Fluorometer (Thermo Fisher, USA) and quality was checked using 4150 TapeStation automated electrophoresis system (Agilent, USA). Sequencing was performed in paired-end mode with a S1 and S2 flow cell using NovaSeq 6000 sequencer (Illumina, USA).

### Human nasal swab scRNA-seq and pre-processed data treatment

Nasal swab data stems from a publication and is freely available.^8^ We, however, asked the authors to share a version of the data set with non-normalized counts, suitable for pseudo bulk-based differential expression analysis. Please refer to the original publication for a detailed description of the experimental process, raw data pre-processing, quality control and integration.^8^

Integrated expression data were normalized by total UMI count per cell (log10(TP10k+1)) and scaled using Seurat (version 4.3.0) R package.^73^ Subsequently, immune cell populations of interest (T and plasmacytoid dendritic cells (pDCs)) were subset using cluster annotation from the original publication and treated separately. Principal component analysis (PCA) was performed on top 3000 variable genes and the first 15 PCs were used to construct a KNN graph and cluster the cells using Eucledian distance and Louvain algorithm (FindVariableFeatures, RunPCA, FindNeighbors and FindClusters functions, in that order) with the resolution of 0.5. This process has resulted in 14 and 8 clusters for T lymphocytes and pDCs, respectively. UMAP dimension reduction was also computed with the first 15 PCs in both cases using the default parameters. Further analysis was done using the same pipeline for both the nasal swab data set and the PBMC data set, please refer to the corresponding sections of the methods.

### Pre-processing and integration of 10x Genomics Chromium PBMC scRNA-seq data (cohort 1 & 2)

Raw sequencing data were processed with CellRanger’s (v5) multi workflow and aligned against the GRCh38 reference, including TotalSeq C hashtag barcodes and VDJ data.

Cells from pooled samples were demultiplexed using a combination of HTODemux implemented in Seurat (v.4.3.0) and vireo (v0.5.6) after scoring common variants from the 1000Genomes project with cellsnp-lite (v1.2.0).^74,75^ Events classified as “Negative” and “Doublet” by the HTODemux algorithm were assigned an ID via vireo classification.

Demultiplexed batches of measurements were loaded into the environment and normalized separately, where gene expression values were normalized by total UMI counts per cell, multiplied by 10,000 (TP10K) and then log transformed by log10(TP10k+1) with NormalizeData Seurat function. Top 2000 variable features per batch were then selected and used to rank features for integration (FindVariableFeatures and SelectIntegrationFeatures functions). After per-batch scaling of gene expression and calculation of 50 PCs, integration was done using FindIntegrationAnchors and IntegrateData functions with reduction = “rpca” and dims = 1:50 parameters.

### ScRNA-seq data analysis of 10x Chromium data set (cohort 1 & 2) Data quality control

Subsequent to integration, cells were filtered by number of features (over 200 and less than 5000), percent mitochondrial genes (<10 % mitochondrial UMIs) and number of counts per cell (<20000) to exclude debris and doublets.

### Definition of the immune population spaces

Now integrated data were scaled and the first 15 of the newly computed PCs were used for clustering (resolution = 0.8) and UMAP calculation. Resulting 25 clusters were annotated based on their feature expression levels into non-granular subsets of main immune populations (T and B lymphocytes, NK cells, Monocytes, DCs) and subset into separate Seurat objects to be analyzed independently. Clusters that were either a mix of different immune cell lineages or did not have a high enough PTPRC gene expression were annotated as “drop” and excluded from further analysis.

### Refining of separate immune populations and subclustering

To refine the data subsets, we filtered out cells, expressing genes exclusive to other populations. We made sure to clean T lymphocytes of gamma-delta T cells (defined as TRGC1>0 OR TRGC2>0); B lymphocytes of T cells, NK cells and monocytes (by removing CD3E>0 OR CD3G>0 OR CD3D>0 OR NKG7>0 OR CD14>0 cells); Monocyte subset was refined through removing NK cells, B lymphocytes and T lymphocytes (NKG7>0 OR CD19>0 OR CD3E>0 cells).

Following this, integrated data slot was used to rescale data and recalculate PCA values. The number of PCs to use for UMAP and clustering computation was selected by analyzing an elbow plot and it varied for different subsets: 15 for T cells, 14 for B cells, 15 for DCs and 17 for monocytes. NK cells were not analyzed, as mass cytometry did not show many clear patterns in this immune space.

Subclustering was calculated with a resolution of 1 for T cells and 0.5 for all other subsets. UMAP was calculated with default parameters for all subsets. Same as previously, scaling, PCA, UMAP and clustering calculation was done using Seurat R package (v.4.3.0).

### Cluster annotation and statistical testing

Subclustering of separate immune populations has netted 21 clusters (of which 2 were dropped) for T cells, 13 clusters (of which 1 was dropped) for B cells, 11 clusters (of which 1 was dropped) for Monocytes and 11 clusters (of which 1 was dropped) for DCs. Annotation of clusters has been done using a combination of manually pre-selected marker genes and automatically detected cluster-specific genes (using FindAllMarkers function, showing top 10 significantly enriched genes, detected in at least 25% of events and having a log-fold-change of at least 0.25). Manual labels were assigned to each cluster. In some cases, if clusters were highly similar, the same label was assigned, merging the cluster for further analysis.

Annotated cluster abundances were compared between the respective groups and the statistical significance was calculated using the adjusted Wilcox test (Benjamini-Hochberg) for clusters with significant Kruskal-Wallis test (adjusted p-value (Benjamini-Hochberg) < 0.05, adjusted across all annotated clusters in each immune cell compartment). Adult patients with anti-IFN-a autoantibodies were removed from infected adults - infected children comparison to keep the comparison more representative of the general population.

Analysis of cluster frequencies has allowed us to pre-select annotated clusters expanded in acute SARS-CoV-2 infection, thus making further analysis more informative through isolation of infection-specific effects. If not specifically stated otherwise, differential gene expression (DE) analysis and analyses based on its output were done using cells from expanded clusters. For T lymphocyte space, CD8^+^ T lymphocytes were defined as CD8A AND CD8B expressing cells, with CD4^+^ identity being assigned to the rest of the cells.

### Differential Expression (DE) and Gene Ontology (GO) Enrichment Analysis

For the identification of differentially expressed genes between disease groups, we used pseudobulk gene expression, defined as the sum of the raw counts from all cells of each patient among selected clusters of interest. The pseudobulk samples were then normalized and modelled according to the DESeq2 pipeline (v. 1.38.2).^76^ Normalization was done using DESeq2’s median-of-ratios method, the DESeq2 function DESeq() was used to estimate size factors, dispersion, and to fit a generalized linear model for each gene. Patient group was included as a sole factor. Differential expression was assessed using the Wald test for each gene. P-values were adjusted for multiple testing using the Benjamini-Hochberg method to control the false discovery rate (FDR).

For further enrichment analysis, we selected differentially expressed genes with high enough counts (“baseMean” > 50) and p-value lower than 0.05. GO enrichment analysis was performed with the R package “enrichR” (v.3.1) and “GO_Biological_Process_2018” database.^77^

### Gene Set Enrichment Analysis (GSEA)

The log2-fold change of differentially expressed genes from DESeq2 was used to define the ranked gene list used for GSEA. We tested different annotated gene lists for different immune populations (Supplemental Table 3). GSEA was performed with the R package “fgsea” (v. 1.24.0) with 1000 permutations for statistical testing.^78^

### Clonal composition analysis

Our experimental design included VDJ region sequencing to enable clonal composition analysis. We used an R package called scRepertoire (v. 1.8.0) to do that.^79^ scRepertouire interacts with the Seurat object to combine gene expression data and clonotype information that can be called from the VDJC gene or the sequence of the CD3R region. In our case we mostly use the “strict” option for calling the clonotype, which uses both the gene and the nucleotide. Clonal overlap proportions between plasmablasts and other expanded clusters have been calculated via “clonalNetwork” function and used as the basis for the stacked barchart plot, with the clone calling method being set to “gene” in order to increase sensitivity.

### Data visualization

All the graphical visualization of the data was performed in R with the ggplot2 package apart from the heatmaps, which were displayed using the ComplexHeatmap (v. 2.14.0) and pheatmap (v. 1.0.12) packages.^80^

#### Box plots

Box plots are calculated in the style of Tukey, shortly, the center of the box represents the median of the values, the hinges the 25th and 75th percentile and the whiskers are extended no further than the 1.5 * IQR (interquartile range).

#### Heatmap

The heatmap shows the mean value of the z-scaled expression of each marker (or abundance, cytokine concentration) in each cluster or patient group.

#### Dot plot

The dot plot of the signature genes shown was calculated according to the Dotplot Seurat function scaling the expression values by gene.

### Detection of SARS-CoV-2-specific IgG and IgA antibodies

For the detection of IgG and IgA to the S1 domain of the SARS-COV-2 spike (S) protein, anti-SARS-CoV-2 assay was used according to the manufactureŕs instructions (Euroimmun, Lübeck, Germany). Serum samples were tested at a 1:101 dilution using the fully EUROIMMUN Analyzer. Optical density (OD) ratios were calculated by dividing the OD at 450 nm by the OD of the calibrator included in the kit. The calculated OD ratios can be used as a relative measure for the concentration of antibodies in the serum.^81^

### *Ex vivo* functional analyses of T cells Cell purification

Frozen PBMCs of control donors were thawed and washed with a Benzonase-containing medium (RPMI, 2% FCS, Pierce Universal Nuclease, 250U/mL), transferred to MACS buffer (PBS, 0.5% BSA, 2mM EDTA) and counted. Cells were split into CD45RA^+^ and CD45RA^-^ fractions using magnetic separation with CD45RA MicroBeads (Miltenyi Biotec) according to the manufacturer’s protocol. CD45RA-fraction was labeled with eF450 proliferation dye in PBS-for 10 minutes at room temperature (10uM final eF450 concentration). Labeling was stopped by adding 4 volumes of cold complete medium (containing 10% FCS) and cells were incubated on ice for 5 minutes followed by two washing steps with RPMI/5%FCS. CD45RA^+^ fraction was labeled with CFSE proliferation dye in RPMI for 7 minutes at room temperature (5uM final CFSE concentration) and washed thrice with RPMI/5%FCS, after which cells rested on ice for 30 minutes. Labeled cells were then pooled for each donor and volume was adjusted to 2000 cells/ul in complete medium (filtrated RPMI +glutamine, 10% (V/V) heat inactivated FCS, 1% HEPES, 1% NEAA, 1% GlutaMAX, 1% sodium pyruvate, 1% Pen. Strep.).

### Stimulation approach

A 96-well, rounded bottom plate was used for the culture, to accommodate all the challenge conditions and donor groups per batch. If the cell number was high enough, two replicates of each condition per donor were included. Reagents were diluted in 100ul of complete medium to a final concentration of 0.324uM ODN CpG2216, 100ng/ml SEB, 0.5ng/ml or 30ng/ml recombinant IFNa, 1ug/ml B18R, 20mg/ml recombinant IL-1b and 10 ug/ml Anakinra (“Kineret”) in following combinations: SEB, SEB+CpG2216, SEB+CpG2216+B18R, SEB+CpG2216+IL-1b, SEB+CpG2216+Anakinra, SEB+rec. IFNa and SEB+rec. IFNa+IL-1b and placed into respective wells. Following this, 100ul of cell suspension per well have been mixed into the reagent containing medium, thus achieving the equal amount of 200000 cells per well. Cells were then cultured for 96 hours at 37°C and 5% CO2.

### Full-spectrum flow cytometry measurement and analysis

After 96 hours, cell culture supernatant was harvested and frozen at −80°C. Cells were washed with MACS buffer and resuspended in either 50ul (for pooled duplicates) or 30ul (for single wells) of freshly prepared surface staining antibody mix, including a live-dead marker (see Key Resources Table) and incubated for 30 minutes at 4°C in the dark. Marker panel has been designed, based on the results detected in CyTOF and scRNAseq experiments. Subsequently, cells were washed twice with cold MACS buffer and fixed in either 100ul Cytofix/Cytoperm buffer (BD Biosciences, USA, for pooled duplicates) or 50ul Cytofix/Cytoperm buffer (for single wells). After a 20 min incubation at 4°C in the dark, cells were washed twice with Cytoperm wash buffer and resuspended in freshly prepared intracellular staining antibody mix (antibodies were diluted in Cytoperm wash buffer to keep cells permeabilized). Cells were incubated for 30 min at 4°C in the dark, washed once with Cytoperm wash buffer, once with MACS buffer, transferred to plastic FACS tubes, resuspended in 150ul of MACS buffer and immediately measured. Cytek Aurora (Cytek Biosciences, USA) full-spectrum flow cytometer was used to acquire all the samples. SpectroFlo (Cytek Biosciences, USA) software was used for spectral unmixing (compensation) of fluorophore signals using SEB+CpG2216-activated unstained reference samples of adult donors, measured with every batch. Single-stained references were prepared either with compensation beads (CompBeads, BD Biosciences, USA) or SEB+CpG2216-activated adult donors’ cells, depending on the marker expression level and fluorophore intensity. Compensated data was uploaded to OMIQ.ai cloud-based cytometry analysis software for gating and export of population abundances and median signal intensities. Exported data was imported into R programming environment, where it was aggregated and analyzed.

### Quantification of cytokines, chemokines

Plates with frozen cell culture supernatants were thawed on ice and centrifuged at 1000xg at 4°C for 10 minutes to remove particulates. Cytokine/Chemokine/Growth Factor 45-Plex Human ProcartaPlex™ Panel 1 kit (ThermoFischer Scientific, USA) and a custom 18-Plex Human ProcartaPlex™ Mix&Match panel (ThermoFischer Scientific, USA) were used in combination with Luminex xMAP technology-based Bio-Plex 200 system (Bio-Rad Laboratories, Inc., USA) according to manufacturer’s protocols. Resulting data set was imported into R programming environment for analysis. Custom panel specifics can be found in the resources table.

### Blood serum cytokine and chemokine quantification

Blood serum samples of patients and control donors have been measured using CorPlex Human Cytokine 10-plex panel 1 assay (Quanterix Corp., USA) on Simoa SP-X (Quanterix Corp., USA) system and IFN-a assay (Quanterix Corp., USA) on Simoa HD-X (Quanterix Corp., USA) system, respectively. Serum samples were thawed on ice and centrifuged at 10000xg for 5 minutes at 4°C to remove particulates and clarify the sample, after which the assays were performed following the detailed protocols provided by the manufacturer. Resulting data has been imported into R programming environment, where it was aggregated and analyzed.

### *Ex vivo* STAT1 and STAT3 phosphorylation assay Cell purification

Frozen PBMCs of control donors were thawed and washed with a Benzonase-containing medium (RPMI, 2% FCS, Pierce Universal Nuclease, 250U/mL). Volume was adjusted for each donor sample to achieve the concentration of 2000 cells/ul in complete medium (filtrated RPMI +glutamine, 10% (V/V) heat inactivated FCS, 1% Pen. Strep.).

### Stimulation approach

A 96-well, rounded bottom plate was used for the culture, to accommodate all the challenge conditions and donor groups per batch. If the cell number was high enough, two replicates of each condition per donor were included. Reagents were diluted in 100ul of complete medium to a final concentration of 100ng/ml SEB and 30ng/ml recombinant IFNa in following combinations: SEB, SEB+rec. IFNa and placed into respective wells. Following this, 100ul of cell suspension per well have been mixed into the reagent containing medium, thus achieving the equal amount of 200000 cells per well. Cells were then incubated for 25 minutes at 37°C and 5% CO2. 10 minutes before the end of the incubation period, a live-dead staining agent was added (eF506, at 1:100 dilution).

### Flow cytometry measurement and analysis

After 25 minutes of incubation, duplicate wells were pooled in total volume of 200ul, washed in FACS buffer (two rounds, 300xg, 5 min, at room temperature) and immediately fixated using pre-warmed (37°C) commercially available fixation buffer, according to manufacturer’s protocol (BioLegend Cat 420801). After 15 minutes of fixation at 37°C, cells were washed in FACS buffer (2 times, 500xg, 5 min, at room temperature) and subjected to Beriglobin blocking (in 25ul for duplicate wells, 20 minutes at 4°C, 160mg/ml stock, used in 1:50). After a single wash step (500xg, 5 min, RT), cells were resuspended in 50ul (for pooled wells) of freshly prepared surface staining solution and incubated for 30 minutes at 4°C in the dark n (see Resources Table for details on cytometry panels). Two washing steps (FACS buffer, 500xg, 5 min, RT) were followed by permeabilization using 200ul True-Phos perm buffer per well, pre-chilled to −20°C according to the manufacturer’s protocol (BioLegend Cat 425401). After the required 1 hour incubation at −20°C in the dark, permeabilized cells were washed twice in FACS buffer (1000xg, 5 min, RT), resuspended in 100ul of freshly prepared intracellular staining solution (see Resources Table for details on cytometry panels) and incubated for 30 minutes in the dark at room temperature. Cells were subsequently washed twice (FACS buffer, 1000xg, 5 min, RT), resuspended in 100ul FACS buffer per well and stored at 4°C in the dark until measurement next day.

BD FACSymphony A5 Cell Analyzer was (BD Biosciences, USA) used to acquire all the samples. FACSDiva software was used for acquisition and compensation of samples. Single-stained references were prepared either with compensation beads (CompBeads, BD Biosciences, USA) or SEB+IFNa-activated adult donors’ cells, depending on the marker expression level and fluorophore intensity.

Compensated data was uploaded to OMIQ.ai cloud-based cytometry analysis software for gating and export of population abundances and median signal intensities. Exported data was imported into R programming environment, where it was aggregated and analyzed.

### Bulk RNAseq of whole blood of SARS-CoV-2-infected hamsters

Details about the animal experiment itself can be found in the original publication.^40^ To extract RNA from whole hamster blood, previously frozen samples were diluted 1:1 with PBS, added to TrizolLS (Invitrogen) reagent at the manufacturer recommended proportion of 1:3 (sample:TrizolLS) and vortexed. After 20min of inactivation, samples were transferred from the BSL3 to a BSL2 facility. RNA extraction was continued according to the manufacturers protocol. RNA was eluted in RNAse free water and stored at −80°C until sequencing.

For RNA extraction from hamster tissue, 50-100 mg of lung tissue was homogenized using a bead mill (Analytic Jena) for 30sec, then 1ml of Trizol reagent (Invitrogen) was added to the tubes. After an additional homogenisation step (30sec) samples were transferred to fresh tubes and incubated for 20min to ensure inactivation before moving them to the BSL2 facility. RNA was then extracted according to the manufacturers instructions, eluted in RNAse free water and stored at −80°C.

Sequencing libraries were generated using the the QuantSeq 3’ mRNA-Seq V2 Library Prep Kit FWD (Lexogen, cat# 193.384) according the manufacturer’s instruction, and sequenced on a Novaseq X device (Illumina) on a 25B flowcell to a depth of about 20-30 million reads per sample. Read 1 of length 151 nucleotides was aligned to the MesAur 2.0 genome assembly (https://www.ncbi.nlm.nih.gov/genome/11998?genome_assembly_id=1585474). The GTF file used for gene counting was described previously.^82^ For data analysis, the R packages quasR, dplyr and and pheatmap (Kolde, R. (2019). pheatmap: Pretty Heatmaps) were used.^83^ All work involving live SARS-CoV-2 virus was conducted under appropriate biosafety conditions in the BSL-3 facility at Institut für Virologie, Freie Universität Berlin, Germany. All animal experimentation was approved by the competent state authority (Landesamt für Gesundheit und Soziales in Berlin, Germany, permit number 0086/20) and performed in compliance with all relevant national and international guidelines for care and humane use of animals.

## DATA AND CODE AVAILABILITY

Debarcoded, batch-corrected and pre-gated fcs files as well as debarcoded, non-normalized and non-pre-gated fcs files of the CyTOF experiment are deposited at Figshare (<u>Figshare Link</u>). Pre-processed (demultiplexed, integrated, quality-controlled and merged with metadata) scRNAseq data is deposited in rds format at Figshare (<u>Figshare Link</u>). Raw count data of the scRNAseq experiment is deposited at GEO repository under the accession ID GSE271284 (<u>GEO</u> <u>Link</u>, review token: **avipyqsklfslfyj**). R-scripts to analyze the CyTOF and single-cell data are available in Mendeley under (reserved) DOI 10.17632/kz7cpw3bnt.1 (<u>Mendeley Link</u>).

## Supporting information

Resources Table

## ACKNOWLEDGEMENTS

We thank Desireé Kunkel and Jacqueline Keye from the BIH Flow and Mass cytometry Core facility for the help with cytometry data generation, the BIH/MDC Genomics Platform for sequencing, Benedikt Obermayer-Wasserscheid for pre-processing of the sequencing data and the Clinical Study Center (CSC) at the Berlin Institute of Health (BIH) and the Central Biobank of the BIH (ZeBanC) for ongoing support of the PA-COVID-19 and RECAST studies.

We are grateful to the patients and donors volunteering to participate in this study, making this research possible in the first place. This work was supported by the German Research Foundation (DFG): SA1383/3-1 and CRC 1444 - project 427826188 to B.S., 322900939, 454024652 and 432698239 to P.B.; and CRC 1449 - project 431232613 to M.A.M., SFB-TR84 114933180 to L.E.S.; the German Federal Ministry of Education and Research (BMBF): NATON, No. 01KX2121, to P.B., B.O., B.M.; 82DZL009B1 to M.A.M; 01IK20337 - RECAST to B.S., J.R., M.A.M., V.M.C., L.E.S., M.R.; FKZ 01KX2021 - COVIM to B.S., L.E.S., and V.M.C., 01GM2202C to P.B., VARIPath (01KI2021) to V.M.C.; the European Research Council: 825392 - RESHAPE to B.S., ERC Consolidator Grant AIM.imaging.CKD, No. 101001791 to P.B.; the Jürgen Manchot Foundation to A.H. and a Charité 3R project to B.S..

## AUTHOR CONTRIBUTIONS

Conceptualization: M.R., M.A.M., V.M.C., L.E.S., J.R., B.S.; Methodology: L.P., S.B., P.G., R.A.-G., N.B., C.I., R.D.B., T.M., V.M.C.; Software/Formal Analysis: L.P., P.G., S.V.S., V.M.C., B.S.; Investigation: L.P., S.B., P.G., A.H., K.V., E.W., H.P.D., S.V.S., L.M., T.M.; Resources: S.W., P.G., A.H., B.M., C.L., J.L., R.E., I.L., B.O., J.T., P.B., M.A.M., V.M.C., J.R.; Data Curation: L.P., S.W., S.V.S., J.L.; Writing - Original Draft: L.P., S.V.S., S.Bedoui., C.M., B.S.; Writing - Review & Editing: L.P., P.G., S.V.S., R.D.B., I.L., P.B., S.Bedoui, V.M.C., L.E.S., J.R., B.S.; Visualization: L.P., S.V.S., B.S.; Supervision: C.I., R.D.B., P.B., L.E.S., J.R., B.S.

## RESOURCES: REAGENTS

| REAGENT | SOURCE | IDENTIFIER |
| --- | --- | --- |
| Cytofix/Cytoperm Kit | BD | 554714 |
| Fixation Buffer | Biolegend | 420801 |
| True-Phos™ Perm Buffer | Biolegend | 425401 |
| CpG2216 | Miltenyi | 130-100-243 |
| Staphylococcal enterotoxin B (SEB) | Sigma | S4881-1MG |
| hIFN-α 2a | Shenandoah | 100-54 |
| B18R | R&D Systems, Biotechne | 8185-BR-025 |
| CD38 Inh_78c | Tocris, Biotechne | 6391 |
| Gibco™ HEPES (1 M) | gibco | 15630-056 |
| MEM Nicht essentielle Aminosäuren-Lösung (100x) | SIGMA-ALDRICH | M7145 |
| Gibco™ GlutaMAX™ Supplement | gibco | 35050-038 |
| Natriumpyruvat (100 mM) | gibco | 11360-039 |
| Penicillin-Streptomycin | SIGMA-ALDRICH | P4333 |
| CD45RA microbeads | Miltenyi | 130-045-901 |
| Invitrogen™ eBioscience™ CFSE | invitrogen | 65-0850-84 |
| eBioscience™ Cell Proliferation Dye eFluor™ 450 | Thermo Fisher | 88701 |
| RPMI medium | gibco | 31870-025 |
| Pierce Universal Nuclease | Thermo Fisher | 88702 |
| FCS heat-inactivated | Sigma | S0615-500mL |
| Bovine Serum Albumin (BSA) | Miltenyi | 130 091 376 |
| PBS - | gibco | 14190-094 |
| EDTA | SIGMA-ALDRICH | 03690 |
| Beriglobin | CSL Behring | PZN 4616123 |
| Pierce 16% Formaldehyde (w/v), Methanol-free | Thermo Fisher | Cat# 28908 |
| Maxpar PBS | Fluidigm | Cat# 201058 |
| Maxpar Cell Staining buffer | Fluidigm | Cat# 201068 |
| Maxpar X8 Multimetal Labeling Kit | Fluidigm | Cat# 201300 |
| Nuclease-Free Water | Invitrogen | Cat# AM9937 |
| TotalSeq-C0251 anti-human Hashtag 1 | Biolegend | Cat# 394661; RRID: AB_2801031 |
| TotalSeq-C0252 anti-human Hashtag 2 | Biolegend | Cat# 394663; RRID: AB_2801032 |
| TotalSeq-C0253 anti-human Hashtag 3 | Biolegend | Cat# 394665; RRID: AB_2801033 |
| TotalSeq-C0254 anti-human Hashtag 4 | Biolegend | Cat# 394667; RRID: AB_2801034 |
| TotalSeq-C0255 anti-human Hashtag 5 | Biolegend | Cat# 394669; RRID: AB_2801035 |
| TotalSeq-C0256 anti-human Hashtag 6 | Biolegend | Cat# 394671; RRID: AB_2820042 |
| TotalSeq-C0257 anti-human Hashtag 7 | Biolegend | Cat# 394673; RRID: AB_2820043 |
| TotalSeq-C0258 anti-human Hashtag 8 | Biolegend | Cat# 394675; RRID: AB_2820044 |
| TotalSeq-C0259 anti-human Hashtag 9 | Biolegend | Cat# 394677; RRID: AB_2820045 |
| TotalSeq-C0260 anti-human Hashtag 10 | Biolegend | Cat# 394679; RRID: AB_2820046 |
| Chromium Next GEM Chip G Single Cell Kit | 10x Genomics | Cat# 1000120 |
| Chromium Next GEM Single Cell 5' Library and Gel Bead Kit v1.1 | 10x Genomics | Cat# 1000167 |
| Chromium Single Cell 5' Library Construction Kit | 10x Genomics | Cat# 1000020 |
| Chromium Single Cell V(D)J Enrichment Kit, Human B Cell | 10x Genomics | Cat# 1000016 |
| Chromium Single Cell V(D)J Enrichment Kit, Human T Cell | 10x Genomics | Cat# 1000005 |
| Single Index Kit T Set A | 10x Genomics | Cat# 1000213 |
| Single Index Kit N Set A | 10x Genomics | Cat# 1000212 |
| Chromium Single Cell 5' Feature Barcode Library Kit | 10x Genomics | Cat# 1000080 |
| Qubit 1X dsDNA HS Assay Kit | Thermo Fisher Scientific | Cat# Q33231 |
| High Sensitivity DNA Kit | Agilent | Cat# 5067-4626 |
| High Sensitivity D5000 ScreenTape | Agilent | Cat# 5067-5592 |
| Simoa CORPLEX™ CYTOKINE PANEL (IFN-γ, IL-1β, IL-4, IL-5, IL-6, IL-8, IL-10, IL-12P70, IL-22, TNFα) | Quanterix | Cat# 85-0329 |
| Cytokine/Chemokine/Growth Factor 45-Plex Human ProcartaPlex™ Panel 1 | Invitrogen, Thermo Fisher Scientific | Cat# EPX450-12171-901 |
| 18-Plex Human ProcartaPlex™ Mix&Match panel (custom, IFN-γ, IL-1a, IL-1b, IL-12p70, IL-13, IL-17aA, IL-2, IL-21, IL-23, IL-4, IL-5, IL-6, IL-8, IL-9, IP-10, RANTES, S100A8/A9, TNFα) | Invitrogen, Thermo Fisher Scientific | Cat# EPX450-12171-901 |
| IL-1b | Miltenyi | Cat# 130-095-374 |
| Anakinra „Kineret“ | Sobi |  |

## RESOURCES: CyTOF PANEL

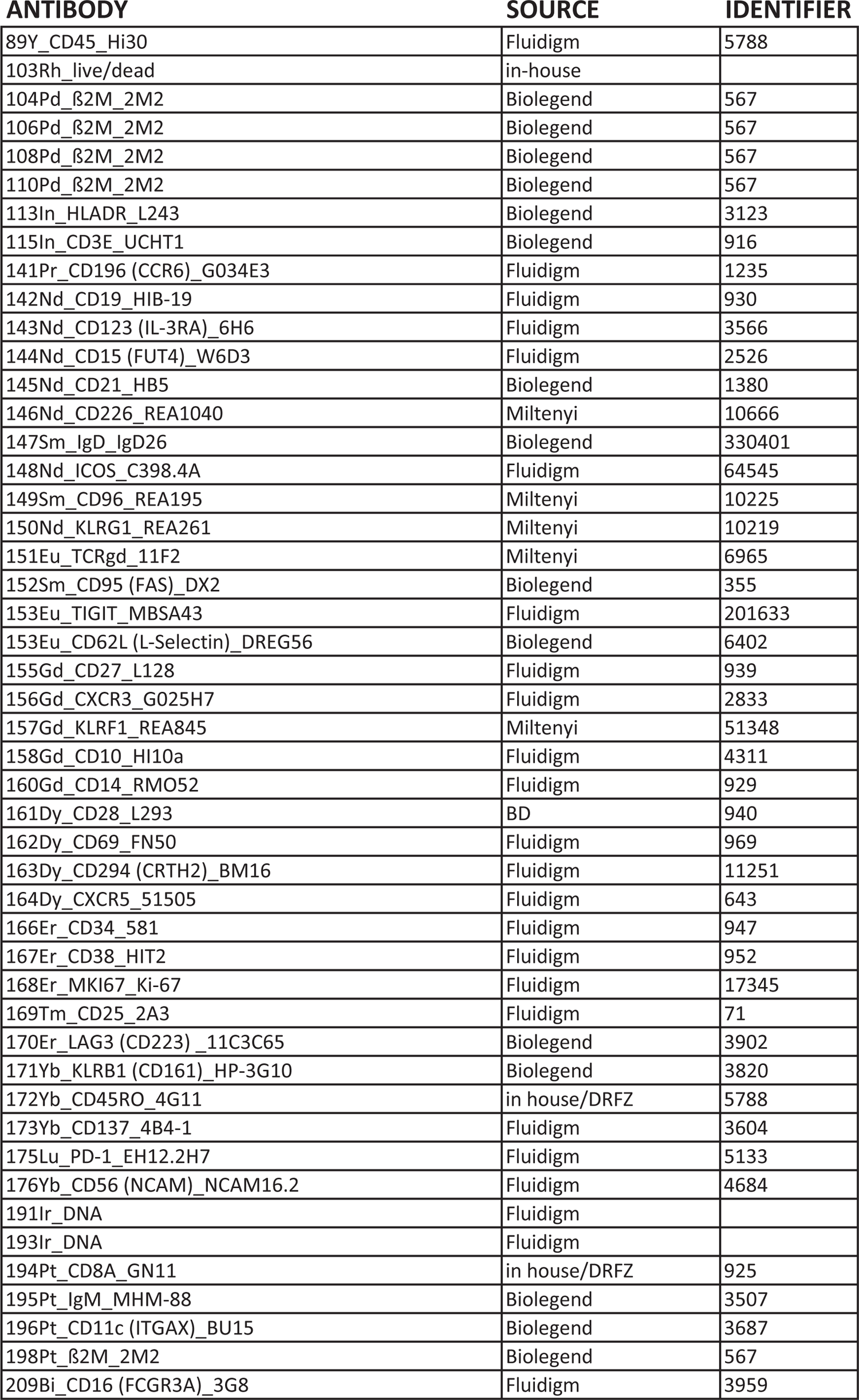

## RESOURCES: FULL SPECTRUM FLOW CYTOMETRY

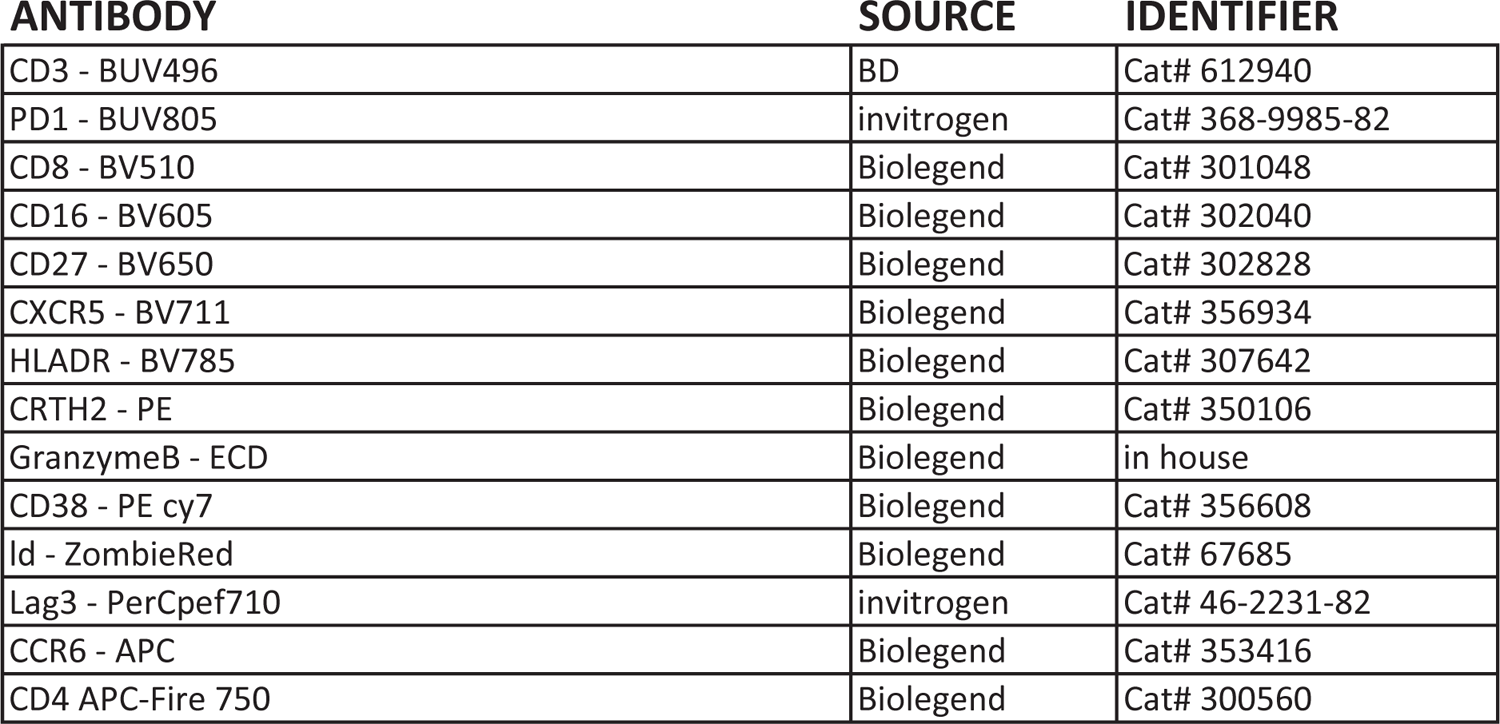

## RESOURCES: FLOW CYTOMETRY

| ANTIBODY | SOURCE | IDENTIFIER |
| --- | --- | --- |
| BD Phosflow™ Alexa Fluor® 488 Mouse Anti-Stat1 (pY701), Clone 4a | BD Biosciences | Cat# 612596, RRID: AB_399879 |
| BD Phosflow™ PE Mouse Anti-Stat3 (pY705), Clone 4/P-STAT3 | BD Biosciences | Cat# 562072, RRID: AB_10893601 |
| APC/Cyanine7 anti-human CD4 Antibody, Clone RPA-T4 | BioLegend | Cat# 300518 |
| eBioscience™ Fixable Viability Dye eFluor™ 506 | Thermo Fisher Scientific | Cat# 65-0866-14 |
| CD19 Pacific Blue | BioLegend | Cat# 302232 |
| BD Horizon™ BUV737 Mouse Anti-Human CD8, Clone SK1 | BD Biosciences | Cat# 612754 |
| BD Horizon™ BUV496 Mouse Anti-Human CD3, Clone UCHT1 | BD Biosciences | Cat# 612940 |

## RESOURCES: SOFTWARE AND ALGORITHMS

| RESOURCE | SOURCE | IDENTIFIER |
| --- | --- | --- |
| R | https://cran.r-project.org/ | 4.2.2 Patched |
| Ubuntu 20.04.5 LTS under WSL2, Windows 11 | Microsoft, Canonical |  |
| Visual Studio Code | Microsoft |  |
| omiq.ai (cloud-based) | Dotmatics |  |
| BatchAdjust (R script) | https://github.com/CUHIMSR/CytofBatchAdjust | used as inspiration for proprietary implementation |
| cytofkit (R package) | https://github.com/JinmiaoChenLab/cytofkit | 0.99.0 |
| uwot (R package) | https://github.com/jlmelville/uwot | 0.1.14 |
| ComplexHeatmap (R package) | http://bioconductor.org/packages/release/bioc/html/ComplexHeatmap.html | 2.14.0 |
| edgeR (R package) | https://bioconductor.org/packages/release/bioc/html/edgeR.html | 3.40.1 |
| ggplot2 (R package) | https://cran.r-project.org/web/packages/ggplot2/index.html | 3.4.2 |
| cowplot | https://cran.r-project.org/web/packages/cowplot/index.html | 1.1.1 |
| ggpubr | https://cran.r-project.org/web/packages/ggpubr/index.html | 0.5.0 |
| ggrepel | https://github.com/slowkow/ggrepel | 0.9.2 |
| ggfortify | https://cran.r-project.org/web/packages/ggfortify/index.html | 0.4.15 |
| flowCore | https://bioconductor.org/packages/release/bioc/html/flowCore.html | 2.10.0 |
| tidyverse | https://cran.r-project.org/web/packages/tidyverse/index.html | 2.0.0 |
| SeuratDisk | https://github.com/mojaveazure/seurat-disk | 0.0.0.9020 |
| Seurat | https://satijalab.org/seurat/ | 4.3.0 |
| stringr | https://stringr.tidyverse.org/ | 1.5.0 |
| DESeq2 | https://bioconductor.org/packages/release/bioc/html/DESeq2.html | 1.38.2 |
| GSVA | https://bioconductor.org/packages/release/bioc/html/GSVA.html | 1.46.0 |
| enrichR | https://cran.r-project.org/web/packages/enrichR/vignettes/enrichR.html | 3.1 |
| fgsea | https://bioconductor.org/packages/release/bioc/html/fgsea.html | 1.24.0 |
| quasR | doi: 10.1093/bioinformatics/btu781 |  |
| pheatmap | Kolde, R. (2019). pheatmap: Pretty Heatmaps |  |
| class | https://cran.r-project.org/web/packages/class/index.html | 7.3-22 |

## RESOURCES: DATA

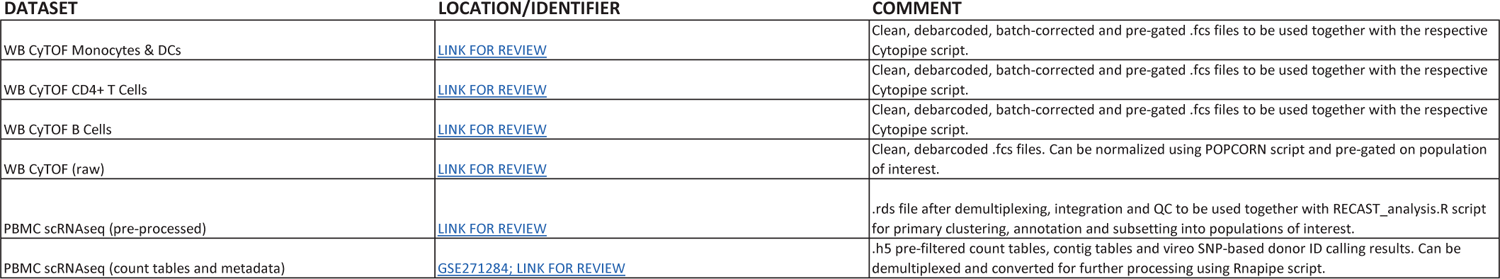

## RESOURCES: CODE

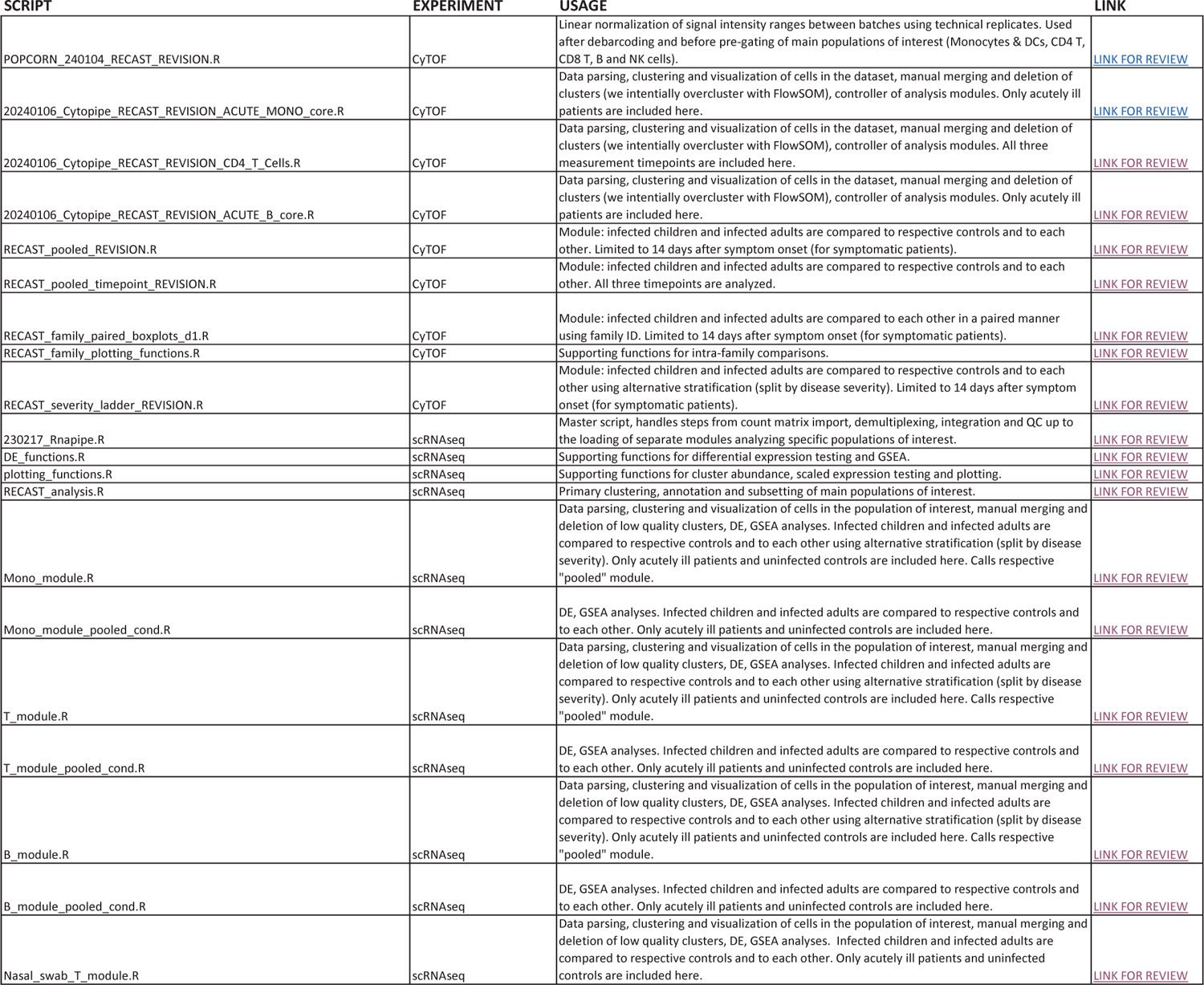

---

**Figure S1.**
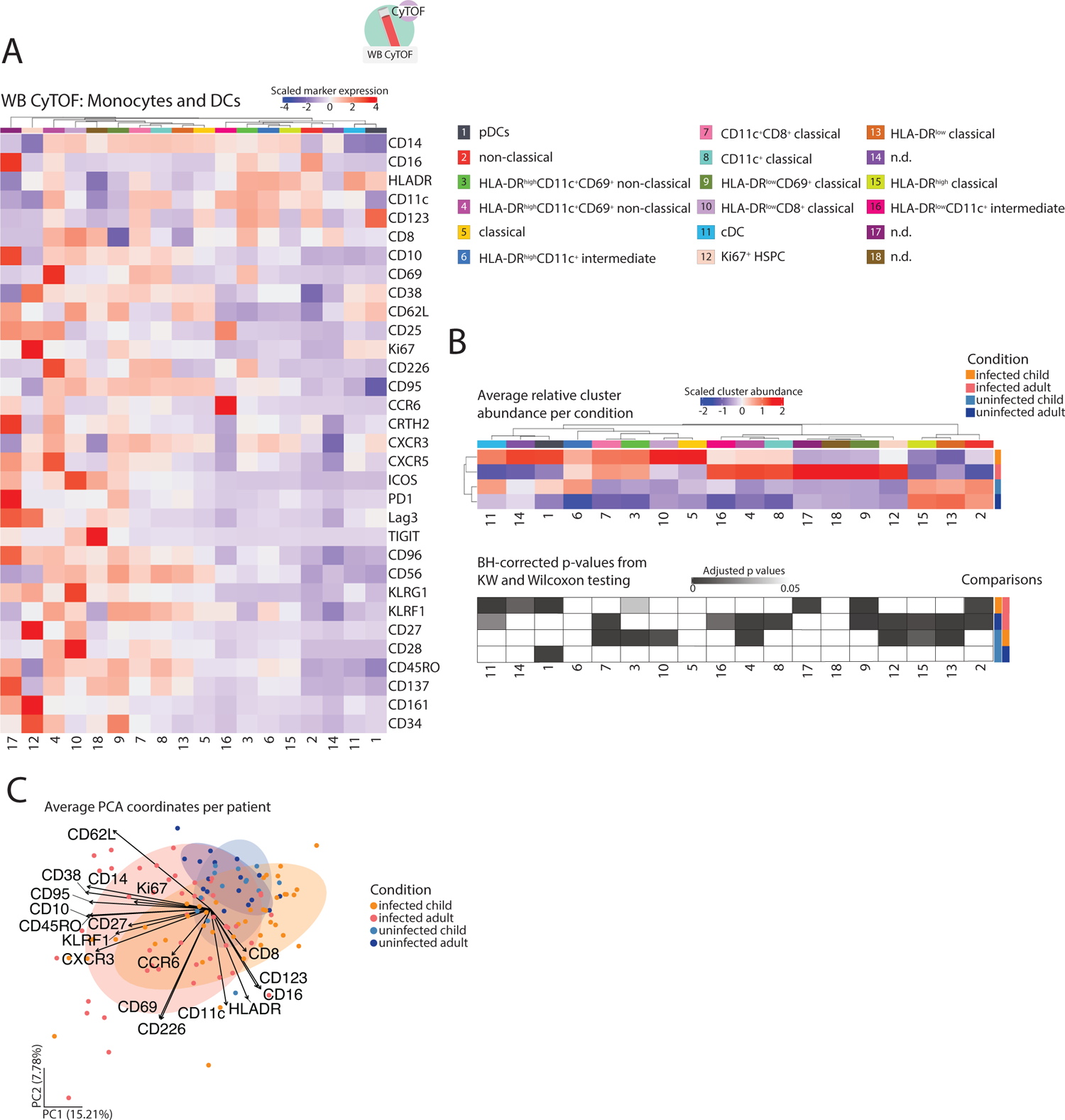
Annotation and statistical analysis of monocyte clusters identified by CyTOF. (A) Heatmap, containing z-score normalized average expression values of each monocyte & dendritic cell cluster resulting from a semi-supervised FlowSOM clustering approach of CyTOF data from whole blood samples (all the patients included in the dataset and used for clustering: uninfected children (RECAST n = 13; median age = 7), uninfected adults (RECAST n = 25; median age = 48), infected children (RECAST study n = 48; median age = 8; asymptomatic n = 11, symptomatic n = 37) and adults (RECAST n = 21, PA-COVID study n = 30; median age = 47; mild n = 37, severe n = 10, severe with IFN autoantibodies n = 4). The dendrogram in the upper part of the panel shows the result of distance-based hierarchical clustering. A color-coded annotation of each cluster is presented on the right. (B) Heatmap, showing scaled mean cluster abundance of monocyte and DC clusters for each patient group (above) and a heatmap presenting the results of statistical testing for all relevant comparisons (below; Benjamini-Hochberg-corrected p values from non-parametric Kruskal-Wallis with post-hoc Wilcoxon). Severe patients with IFN autoantibodies were excluded from the analysis. (C) PCA plot, presenting average principal component 1 and 2 coordinates of all monocyte and DC cells of each patient (colored according to patient group). The plot contains arrows labeled with marker names. These arrows represent the importance of each marker for the calculation of the components. The direction shows which principal components the marker correlates with, and the length of the arrow illustrates the magnitude of the variable’s contribution to the principal components. Each arrow points in the direction where the variable it represents increases in the principal component space and longer arrows indicate that the variable has a strong influence on the principal components. Severe patients with IFN autoantibodies were excluded from the analysis.

**Figure S2.**
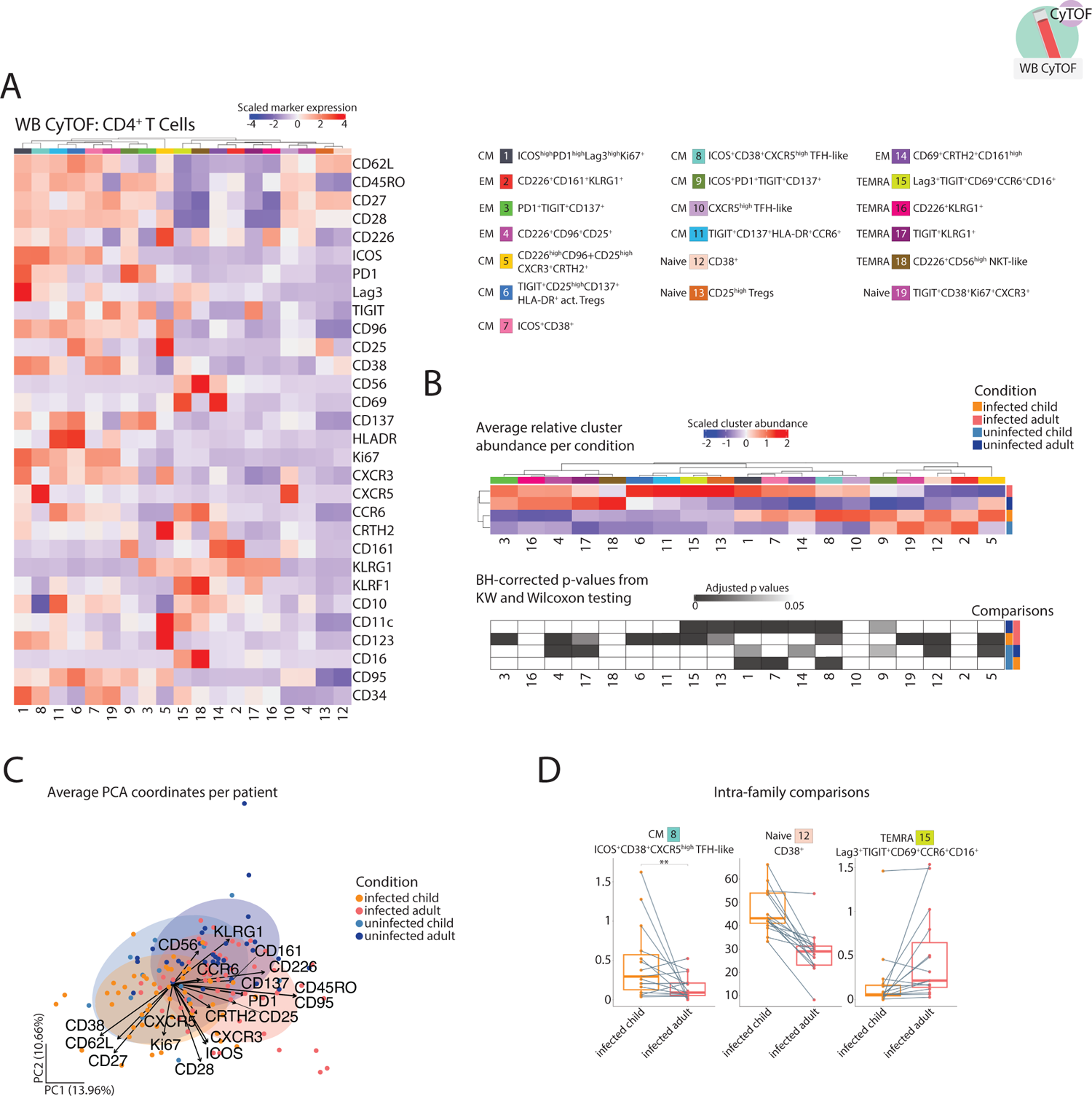
Annotation and statistical analysis of CD4^+^ T cell clusters identified by CyTOF. (A) Heatmap, containing z-score normalized average expression values of each CD4^+^ T cell cluster resulting from a semi-supervised FlowSOM clustering approach of CyTOF data from whole blood samples (all the patients present in the dataset and used for clustering including follow-up measurements of some patients done approximately two weeks and six months after the first, acute infection phase measurement, were included for the heatmap calculation). Patients groups shown are: uninfected children (RECAST n = 13; median age = 7), uninfected adults (RECAST n = 25; median age = 48), infected children (RECAST study n = 53; median age = 8; asymptomatic n = 11, symptomatic n = 42) and adults (RECAST n = 24, PA-COVID study n = 51; median age = 51; mild n = 45, severe n = 24, severe with IFN autoantibodies n = 6). The dendrogram in the upper part of the panel shows the result of distance-based hierarchical clustering. A color-coded annotation of each cluster is presented on the right. (B) Heatmap, showing scaled mean cluster abundance of CD4^+^ T cell clusters for each patient group (above) and a heatmap presenting the results of statistical testing for all relevant comparisons (below; Benjamini-Hochberg-corrected p values from non-parametric Kruskal-Wallis with post-hoc Wilcoxon). Importantly, only samples measured during the acute phase of infection (defined as first 14 days after symptom onset) were used for calculation. Severe patients with IFN autoantibodies were excluded from the analysis. Group composition is thus as follows: uninfected children (RECAST n = 13; median age = 7), uninfected adults (RECAST n = 25; median age = 48), infected children (RECAST study n = 48; median age = 8; asymptomatic n = 11, symptomatic n = 37) and adults (RECAST n = 21, PA-COVID study n = 26; median age = 45; mild n = 37, severe n = 10). (C) PCA plot, presenting average principal component 1 and 2 coordinates of all CD4^+^ T cells of each patient (colored according to patient group). The plot contains arrows labeled with marker names. These arrows represent the importance of each marker for the calculation of the components. The direction shows which principal components the marker correlates with and the length of the arrow illustrates the magnitude of the variable’s contribution to the principal components. Simply put, each arrow points in the direction where the variable it represents increases in the principal component space and longer arrows indicate that the variable has a strong influence on the principal components. Severe patients with IFN autoantibodies were excluded from the analysis. (D) Box plots, showing relative frequencies of annotated clusters, resulting from the FlowSOM algorithm, applied to the CD4^+^ T cell CyTOF data. Cluster abundance is compared between infected children (RECAST study n = 20; median age = 9; asymptomatic n = 3, symptomatic n = 17) and infected adults (RECAST study n = 19; median age = 39; mild n = 19) within the same family. Abundances were averaged, if multiple parents (mother and father) or children were measured in a family (family n = 15). Lines connect parents and children from the same family. Only samples measured during the acute phase of infection (defined as first 14 days after symptom onset) are shown. Statistical testing was done using pairwise Wilcoxon rank sum test. *, p < 0.05; **, p < 0.01; ***, p < 0.001; ****, p < 0.0001.

**Figure S3.**
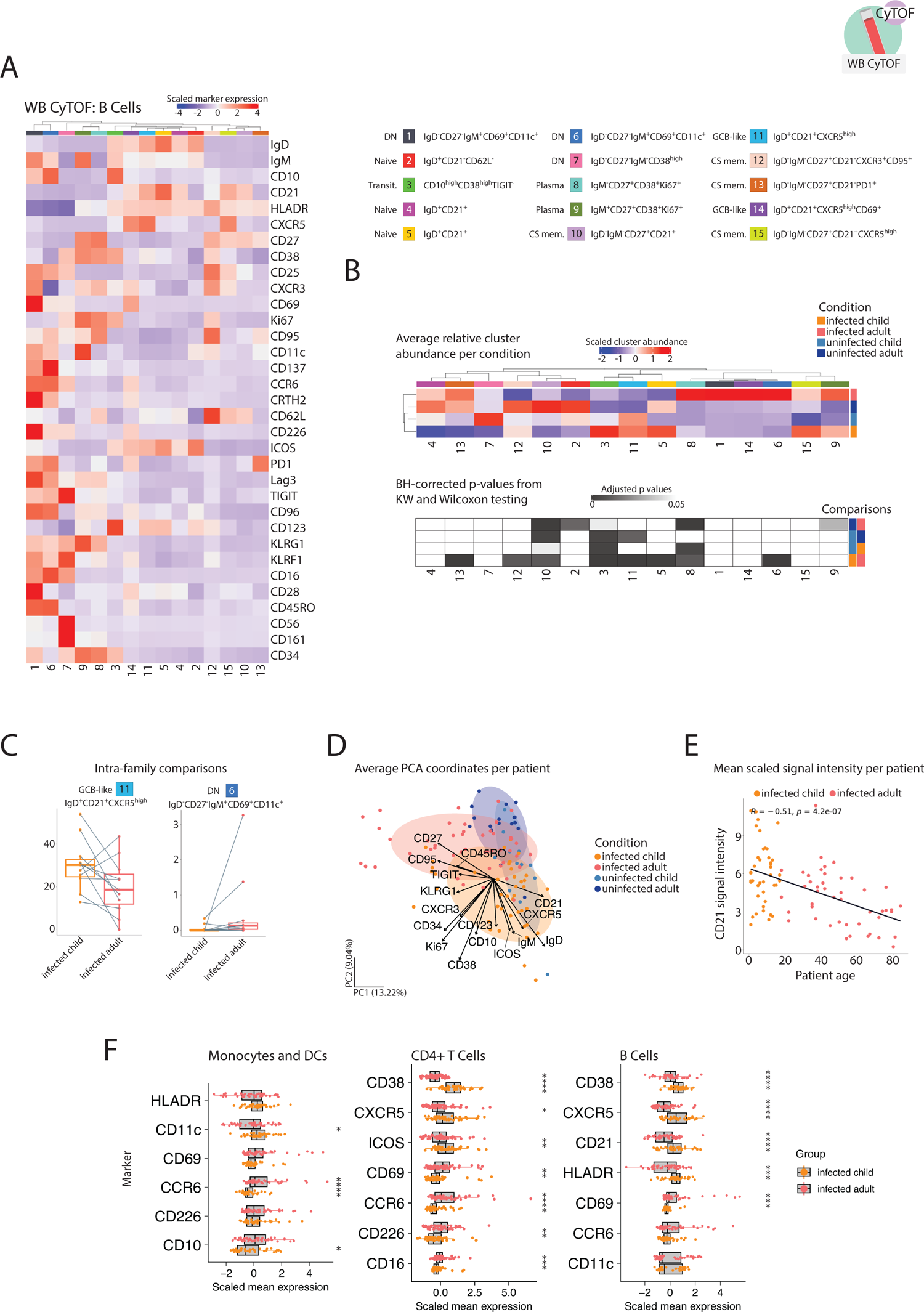
Annotation and statistical analysis of B cell clusters identified by CyTOF, common cross-leukocyte pattern of age-dependency in infection-induced activation marker expression. (A) Heatmap, containing z-score normalized average expression values of each B cell cluster resulting from a semi-supervised FlowSOM clustering approach of CyTOF data from whole blood samples (all the patients included in the dataset and used for clustering: uninfected children (RECAST n = 13; median age = 7), uninfected adults (RECAST n = 25; median age = 48), infected children (RECAST study n = 48; median age = 8; asymptomatic n = 11, symptomatic n = 37) and adults (RECAST n = 21, PA-COVID study n = 30; median age = 47; mild n = 37, severe n = 10, severe with IFN autoantibodies n = 4)). The dendrogram in the upper part of the panel shows the result of distance-based hierarchical clustering. A color-coded annotation of each cluster is presented on the right. (B) Heatmap, showing scaled mean cluster abundance of B cell clusters for each patient group (above) and a heatmap presenting the results of statistical testing for all relevant comparisons (below; Benjamini-Hochberg-corrected p values from non-parametric Kruskal-Wallis with post-hoc Wilcoxon). Severe patients with IFN autoantibodies were excluded from the analysis. (C) Box plots, showing relative frequencies of annotated clusters, resulting from the FlowSOM algorithm, applied to the B cell CyTOF data. Cluster abundance is compared between infected children (RECAST study n = 16; median age = 9; asymptomatic n = 2, symptomatic n = 14) and infected adults (RECAST study n = 14; median age = 38; mild n = 14) within the same family. Abundances were averaged, if multiple parents (mother and father) or children were measured in a family (family n = 11). Lines connect parents and children from the same family. Statistical testing was done using pairwise Wilcoxon rank sum test. *, p < 0.05; **, p < 0.01; ***, p < 0.001; ****, p < 0.0001. (D) PCA plot, presenting average principal component 1 and 2 coordinates of all B cells of each patient (colored according to patient group). The plot contains arrows labeled with marker names. These arrows represent the importance of each marker for the calculation of the components. The direction shows which principal components the marker correlates with and the length of the arrow illustrates the magnitude of the variable’s contribution to the principal components. Simply put, each arrow points in the direction where the variable it represents increases in the principal component space and longer arrows indicate that the variable has a strong influence on the principal components. Severe patients with IFN autoantibodies were excluded from the analysis. (E) Scatter plot, showing mean normalized CD21 signal in relationship to patient age for infected patients (linear model fitted to data and Spearman’s rank correlation coefficient shown in black) within B cell CyTOF data. Severe patients with IFN autoantibodies were excluded from the analysis. (F) Box plots, showing mean normalized signals per patient for a curated list of markers for monocytes and DCs, CD4^+^ T cells and B cells. Signals are compared between infected children (RECAST study n = 48; median age = 8; asymptomatic n = 11, symptomatic n = 37) and adults (RECAST n = 21, PA-COVID study n = 26; median age = 45; mild n = 37, severe n = 10). Wilcoxon; *, p < 0.05; **, p < 0.01; ***, p < 0.001; ****, p < 0.0001.

**Figure S4.**
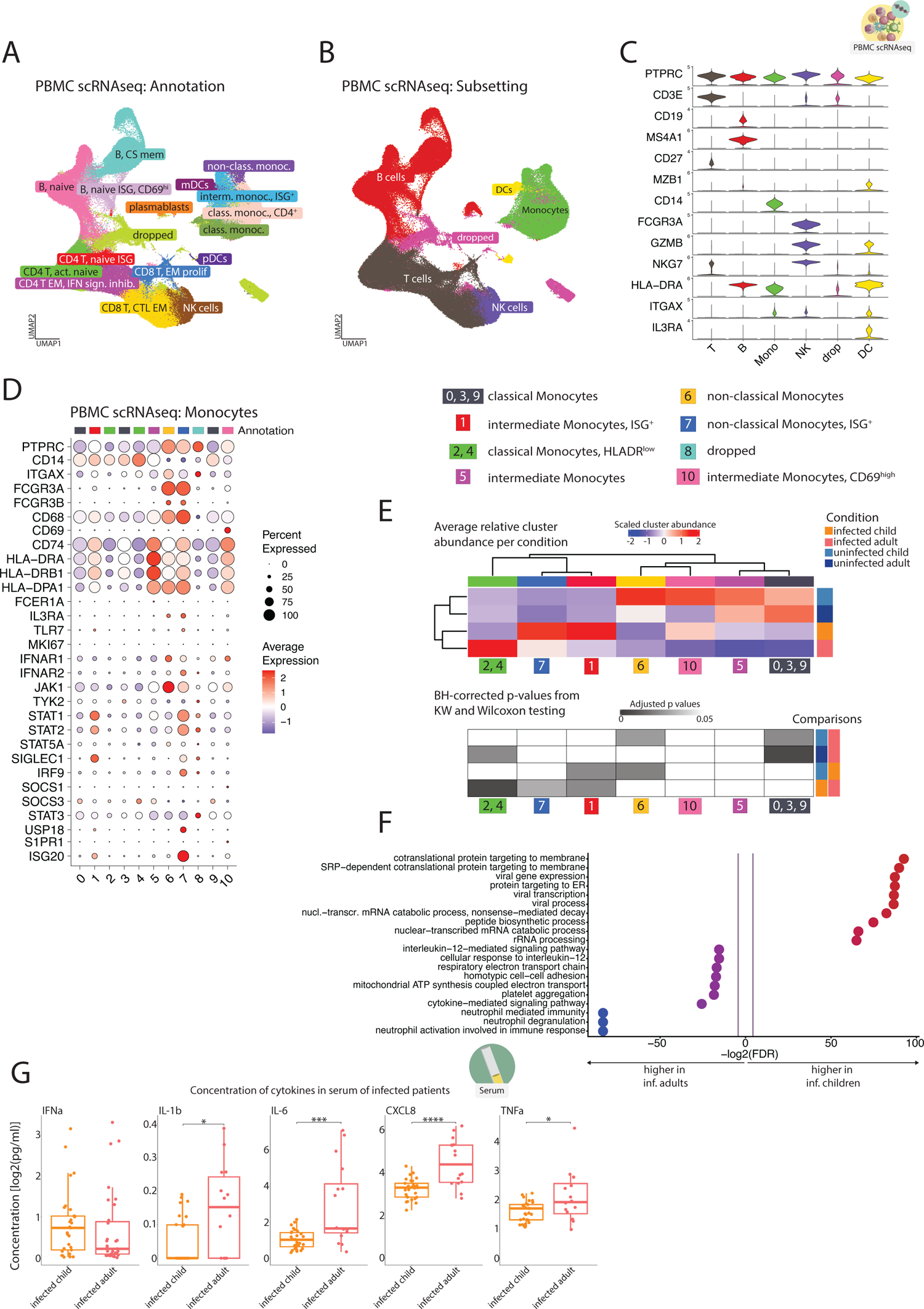
Subsetting of immune populations from PBMC scRNAseq, molecular profiling of monocytes and cytokine concentrations in serum of infected patients point at increased inflammatory state in infected adults. (A) UMAP representation of the total PBMC scRNAseq data. 26 clusters have been produced using a graph-based approach as implemented in Seurat package (KNN graph with Louvain community detection). Clusters were then annotated based on the expression of the most commonly used marker genes of immune metapopulations. UMAP presents all the patients included in the dataset and used for clustering: uninfected children (RECAST study n = 5; median age = 7), uninfected adults (RECAST study n = 4, EICOV/COVIMMUNIZE study n = 4; median age = 66), infected children (RECAST study n = 13; median age = 9; asymptomatic n = 5, symptomatic n = 8) and infected adults (RECAST study n = 1; PA-COVID study n = 18; median age = 67; mild n = 5, severe n = 7, severe with IFN autoantibodies n = 7). (B) UMAP representation of the total PBMC scRNAseq data. 26 annotated clusters from S3A were assigned a metapopulation tag, based on the expression of the most commonly used marker genes. These tags were then used for splitting the dataset into DC, monocyte, NK, T and B cell subsets, which were then separately subclustered to achieve a more granular resolution. Clusters that were either a mix of different immune cell lineages or did not have a high enough PTPRC gene expression were annotated as “dropped” and excluded from further analysis. (C) Violin plots, showing some of the marker genes used for the annotation of the metapopulations. (D) Dotplot, showing average scaled expression of a curated, subset defining list of genes for monocyte cells, subset from the PBMC scRNAseq data. 11 clusters have been produced using a graph-based approach as implemented in Seurat package (KNN graph with Louvain community detection). Some clusters share annotation due to phenotypical similarity and accumulation pattern and are treated as one population for abundance testing in S3E. Annotations of the clusters are shown on the right-hand side from the dotplot. The cluster annotated as “dropped” was of low quality (as concluded from inspecting number of features, n of counts and percentage of mitochondrial genes as well as other population-specific genes). The dotplot presents all the patients included in the dataset and used for clustering: uninfected children (RECAST study n = 5; median age = 7), uninfected adults (RECAST study n = 4, EICOV/COVIMMUNIZE study n = 4; median age = 66), infected children (RECAST study n = 13; median age = 9; asymptomatic n = 5, symptomatic n = 8) and infected adults (RECAST study n = 1; PA-COVID study n = 18; median age = 67; mild n = 5, severe n = 7, severe with IFN autoantibodies n = 7). (E) Heatmap, showing scaled mean cluster abundance of monocyte clusters for each patient group (above) and a heatmap presenting the results of statistical testing for all relevant comparisons (below; Benjamini-Hochberg-corrected p values from non-parametric Kruskal-Wallis with post-hoc Wilcoxon). Severe patients with IFN autoantibodies were excluded from the analysis. (F) Gene set enrichment analysis (GSEA) applied to monocytes from the PBMC scRNAseq experiment and clusters, expanded with infection (1, 4 and 7). GSEA was done using the 189 significantly differentially expressed genes from the infected adults – infected children comparison, shown in volcano plot F3G. R interface EnrichR was used to access Enrichr “GO Biological Process 2018” database for automatic annotation of enriched gene sets. Severe patients with IFN autoantibodies were excluded from the analysis. (G) Box plots of selected pro-inflammatory cytokines’ serum concentrations in infected patients (inf. children RECAST study n = 29; median age = 9; asymptomatic n = 9, symptomatic n = 20; inf. adults RECAST study n = 5, PA-COVID study n = 26; median age = 54; mild n = 12, severe n = 14). Median DPSO in infected children = 5.5, median DPSO in infected adults = 9. Wilcoxon; *, p < 0.05; **, p < 0.01; ***, p < 0.001; ****, p < 0.0001.

**Figure S5.**
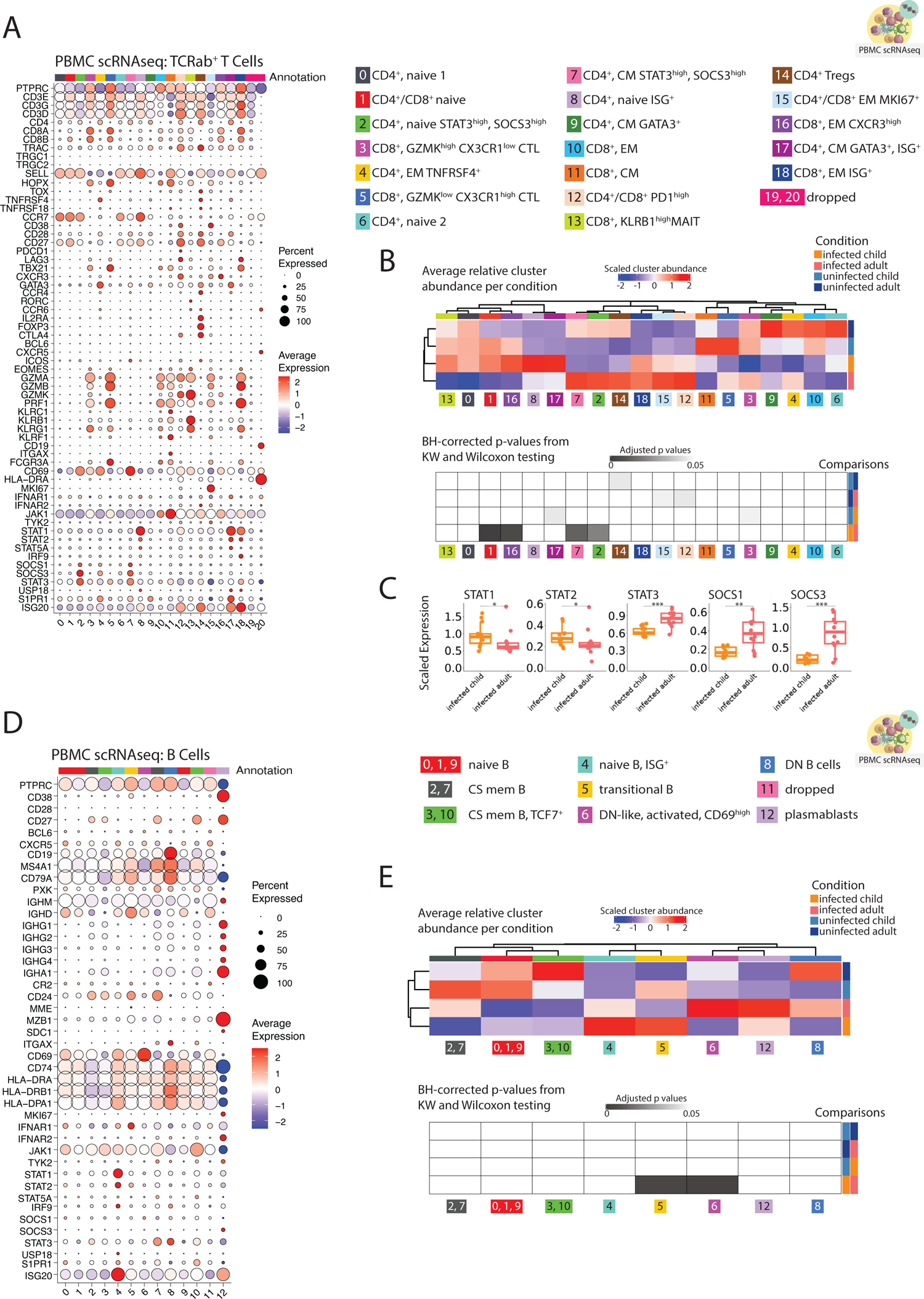
scRNAseq profiling of TCRab^+^ T Cell and B cell space. (A) Dotplot, showing average scaled expression of a curated, subset defining list of genes for TCRab^+^ T cells, subset from the PBMC scRNAseq data. 21 clusters have been produced using a graph-based approach as implemented in Seurat package (KNN graph with Louvain community detection). Annotations of the clusters are shown on the right-hand side from the dotplot. Clusters annotated as “dropped” were of low quality (as concluded from inspecting number of features, n of counts and percentage of mitochondrial genes as well as other population-specific genes). The dotplot presents all the patients included in the dataset and used for clustering: uninfected children (RECAST study n = 5; median age = 7), uninfected adults (RECAST study n = 4, EICOV/COVIMMUNIZE study n = 4; median age = 66), infected children (RECAST study n = 13; median age = 9; asymptomatic n = 5, symptomatic n = 8) and infected adults (RECAST study n = 1; PA-COVID study n = 18; median age = 67; mild n = 5, severe n = 7, severe with IFN autoantibodies n = 7). (B) Heatmap, showing scaled mean cluster abundance of TCRab^+^ T cell clusters for each patient group (above) and a heatmap presenting the results of statistical testing for all relevant comparisons (below; Benjamini-Hochberg-corrected p values from non-parametric Kruskal-Wallis with post-hoc Wilcoxon). Severe patients with IFN autoantibodies were excluded from the analysis. (C) Box plots, showing mean scaled expression of selected genes, involved in IFN signaling, calculated over all TCRab^+^ T cells from clusters expanded with infection (2, 7, 8, 12, 15, 16, 17 and 18) for infected children (RECAST study n = 13; median age = 9; asymptomatic n = 5, symptomatic n = 8) and infected adults (RECAST study n = 1; PA-COVID study n = 11; median age = 63.5; mild n = 5, severe n = 7). Wilcoxon; *, p < 0.05; **, p < 0.01; ***, p < 0.001; ****, p < 0.0001. (D) Dotplot, showing average scaled expression of a curated, subset defining list of genes for B cells, subset from the PBMC scRNAseq data. 13 clusters have been produced using a graph-based approach as implemented in Seurat package (KNN graph with Louvain community detection). Annotations of the clusters are shown on the right-hand side from the dotplot. Cluster annotated as “dropped” was of low quality (as concluded from inspecting number of features, n of counts and percentage of mitochondrial genes as well as other population-specific genes). The dotplot presents all the patients included in the dataset and used for clustering: uninfected children (RECAST study n = 5; median age = 7), uninfected adults (RECAST study n = 4, EICOV/COVIMMUNIZE study n = 4; median age = 66), infected children (RECAST study n = 13; median age = 9; asymptomatic n = 5, symptomatic n = 8) and infected adults (RECAST study n = 1; PA-COVID study n = 18; median age = 67; mild n = 5, severe n = 7, severe with IFN autoantibodies n = 7). (E) Heatmap, showing scaled mean cluster abundance of B cell clusters for each patient group (above) and a heatmap presenting the results of statistical testing for all relevant comparisons (below; Benjamini-Hochberg-corrected p values from non-parametric Kruskal-Wallis with post-hoc Wilcoxon). Severe patients with IFN autoantibodies were excluded from the analysis.

**Figure S6.**
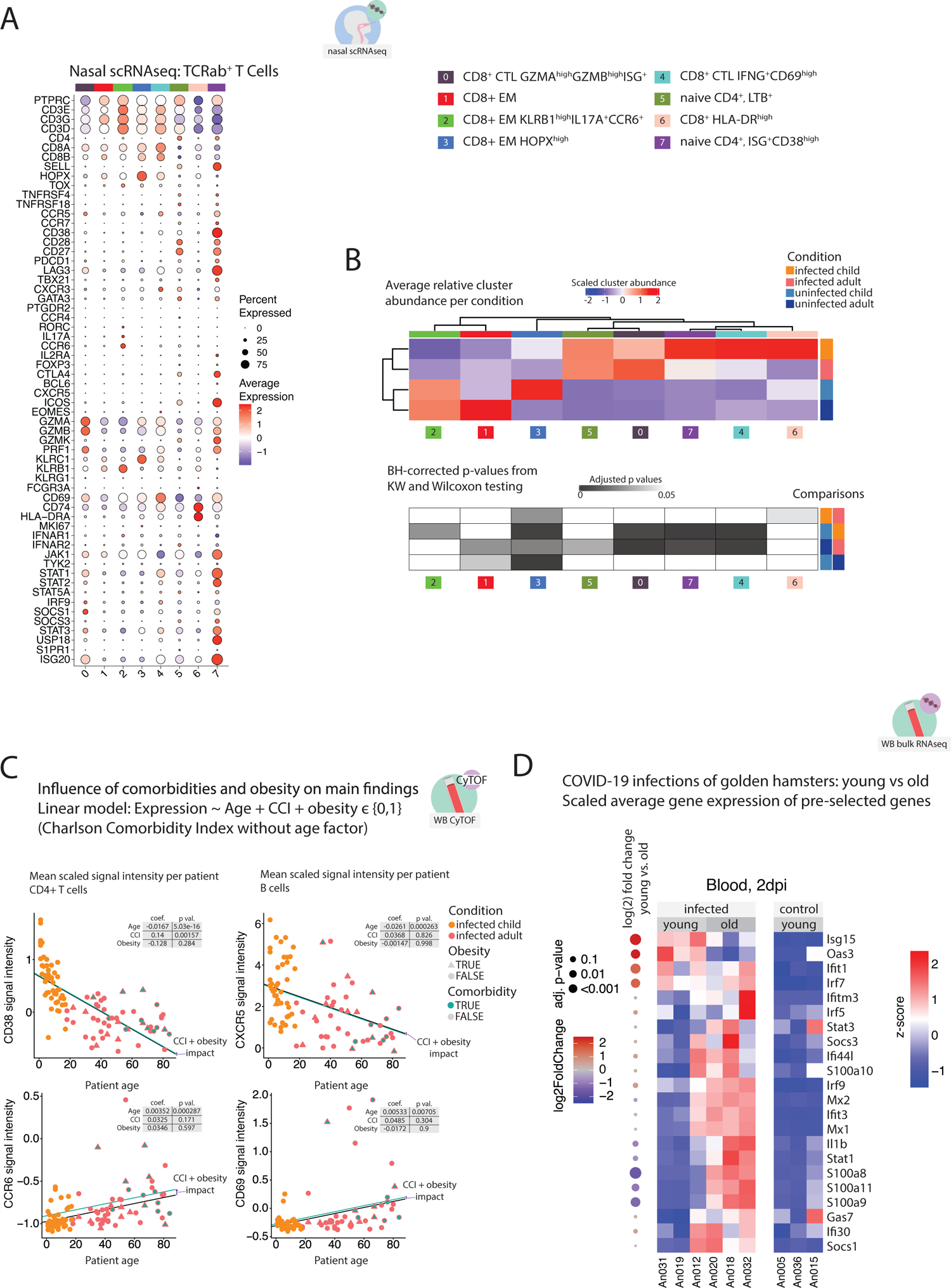
Age-dependent activation profiles are not driven by comorbidities and can be confirmed in a hamster infection model. (A) Dotplot, showing TCRab^+^ cells, subset from the nasal swab scRNAseq data.^8^ 8 clusters have been produced using a graph-based approach as implemented in Seurat package (KNN graph with Louvain community detection). Annotations of the clusters are shown on the right-hand side from the dotplot. The dotplot presents all the patients included in the dataset and used for clustering: uninfected children (RECAST study n = 18; median age = 9), uninfected adults (RECAST study n = 23; median age = 46), infected children (RECAST study n = 24; median age = 9; asymptomatic n = 2, symptomatic n = 22) and infected adults (RECAST study n = 21; median age = 39; mild n = 21). Patients with DPSO range from 0 to 12 days were included (inf. children mean = 4.5, inf. adult mean = 5). (B) Heatmap, showing scaled mean cluster abundance of TCRab^+^ cell clusters for each patient group (above) and a heatmap presenting the results of statistical testing for all relevant comparisons (below; Benjamini-Hochberg-corrected p values from non-parametric Kruskal-Wallis with post-hoc Wilcoxon). (C) Scatterplots, illustrating the results of modelling dependency between age and CD38 and CCR6 expression by CD4^+^ T cells or CXCR5 and CD69 expression by B cells from the CyTOF experiment together with Charlson Comorbidity Index (CCI, w/o age factor) and obesity (as binary variable, 1 if BMI > 30). Light green line shows the impact of CCI and obesity on the intercept value of the model (intercept + CCI coefficient + obesity coefficient). Obesity status is marked by triangular shape of a point, comorbidity by a light green fill. Figure annotation includes the respective coefficients and p values as modelled with linear regression. Infected children (RECAST study n = 48; median age = 8; asymptomatic n = 11, symptomatic n = 37) and adults (RECAST n = 21, PA-COVID study n = 26; median age = 45; mild n = 37, severe n = 10) are shown. (D) Heatmaps, summarizing the differential gene expression analysis results of gene sets identified to be up-regulated in infection-induced leukocyte populations (human PBMC scRNAseq data) determined by bulk RNA sequencing of whole blood samples collected at day 2 after infection from young (n=3, 6 weeks old) and old (n=3, 32-34 weeks old) hamsters (left side) as well as uninfected controls (n=3, right side, 6 weeks old).

**Figure S7.**
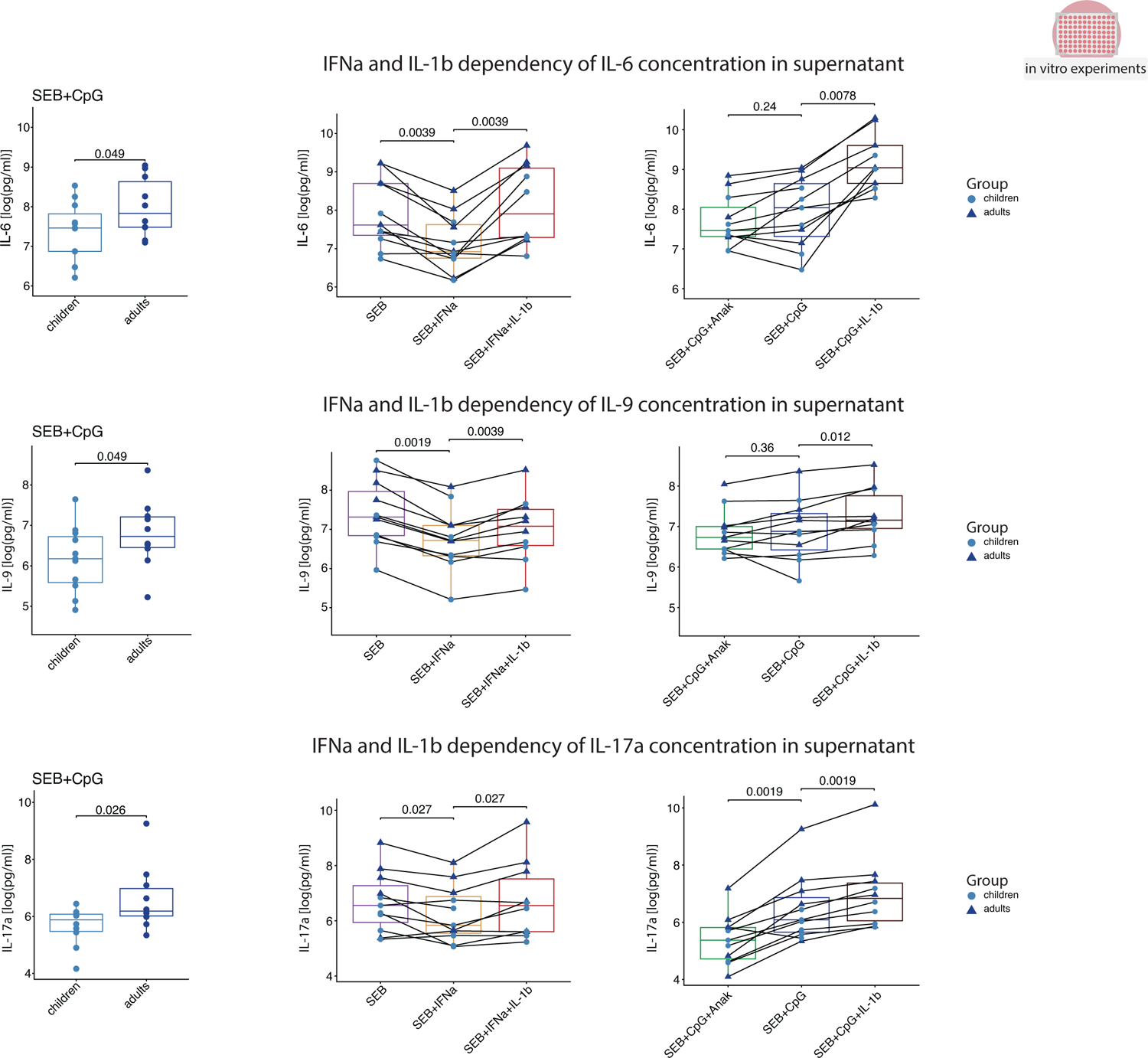
IL-1b-mediated interference with type I IFN signaling explains enhanced cytokine production potential of adult leukocytes. Box plots showing the difference in IL-6, IL-9 and IL-17A release between the two age groups when stimulated by SEB and CpG (first column from the left; uninfected children, RECAST study n = 11; median age = 9; uninfected adult, RECAST study n = 10; median age = 55; one-sided Wilcoxon p value shown), or by different combinations of SEB, CpG, recombinant IFNa (0.5ng/ml), IL-1b and IL-1b inhibitor Anakinra (uninfected children, RECAST study n = 6; median age = 8; uninfected adults, RECAST study n = 5; median age = 55; p values shown for paired Wilcoxon test, adjusted with Hommel procedure).

**TABLE S1:**
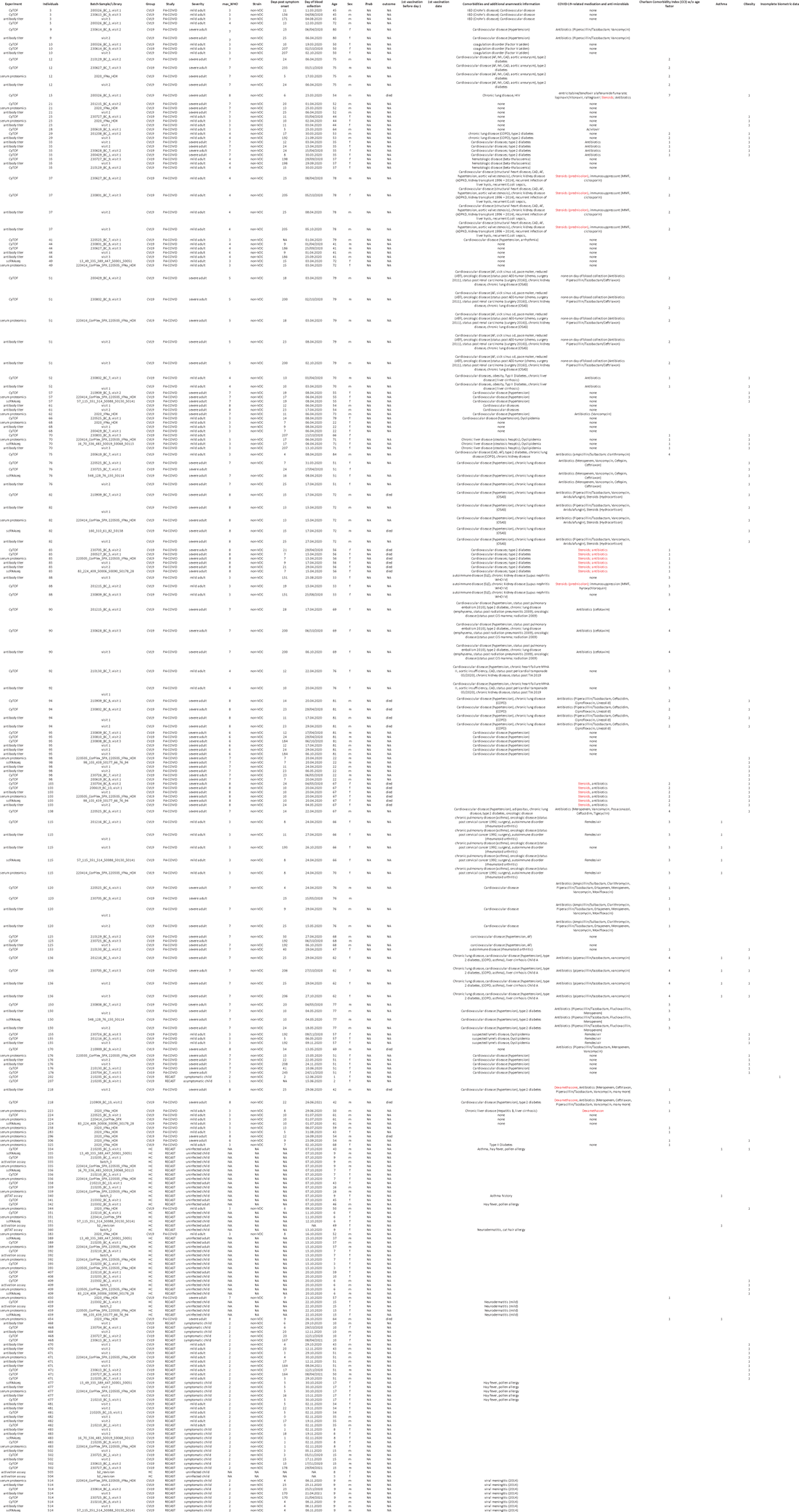
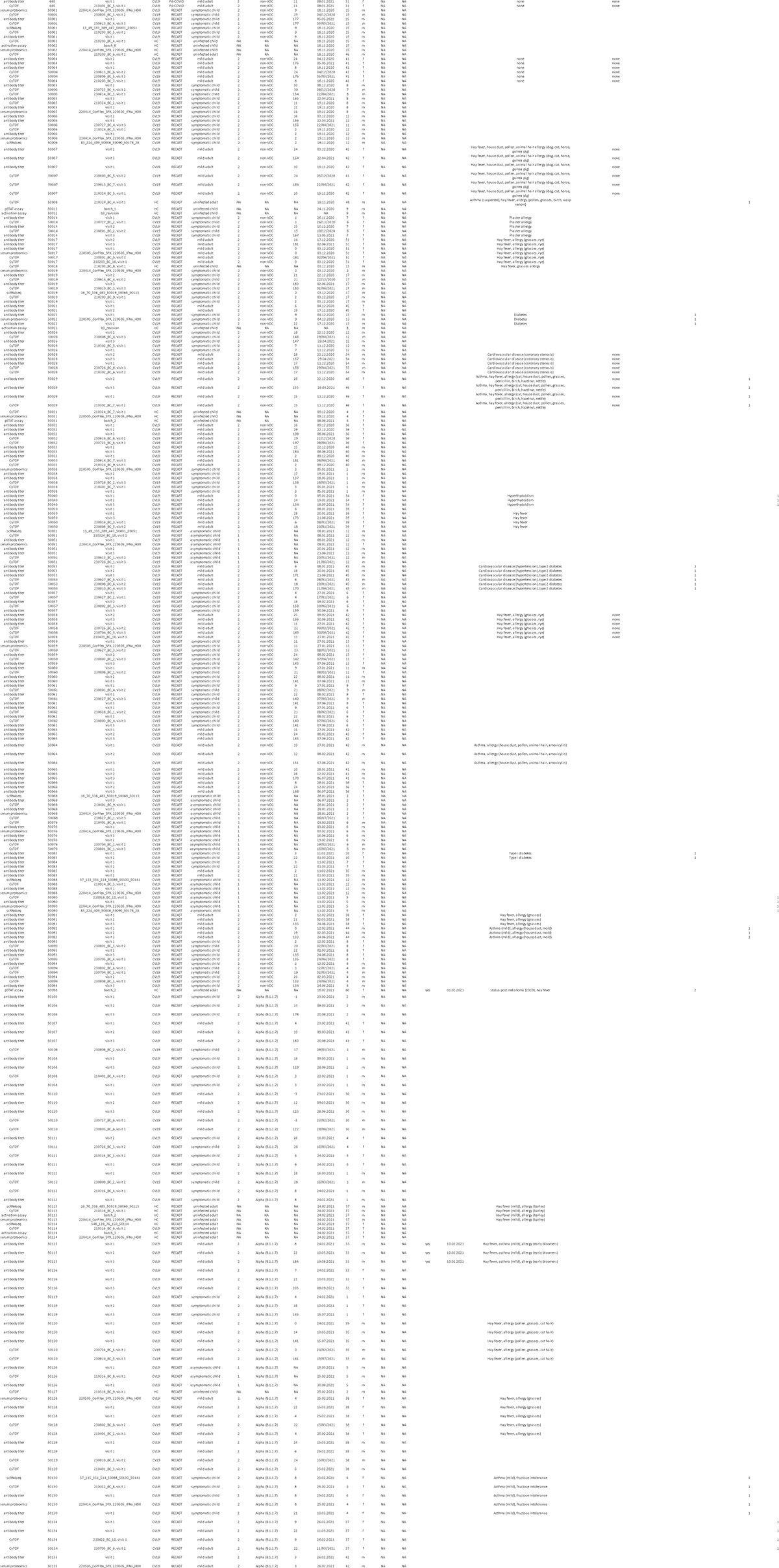
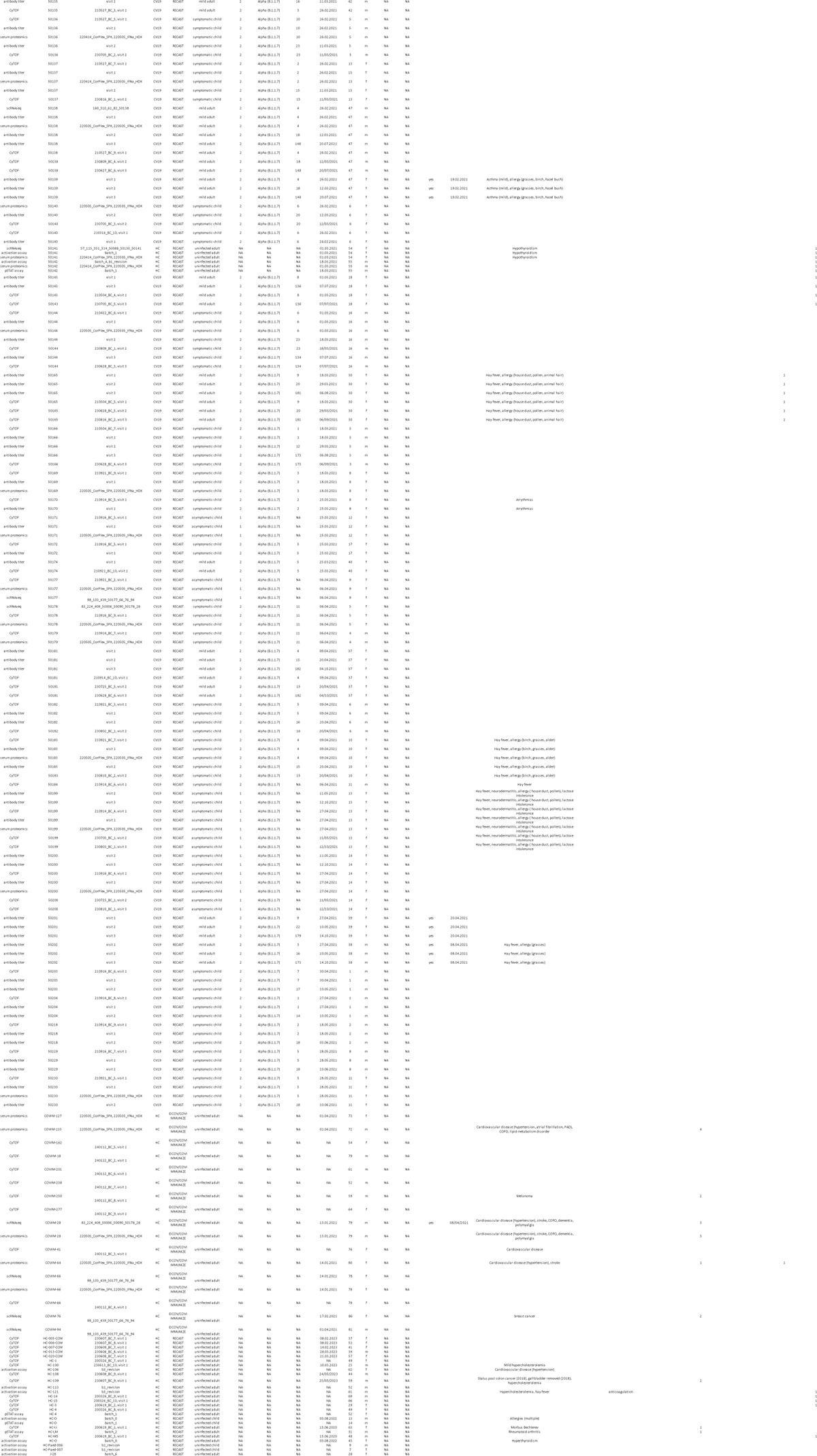
PATIENT SAMPLE OVERVIEW.

**TABLE S1:**
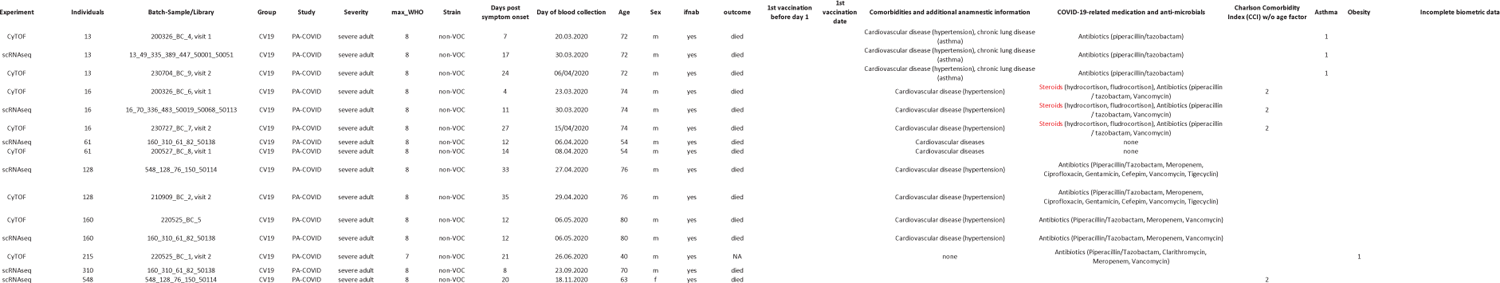
PATIENT SAMPLE OVERVIEW(IFNAAB PATIENTS)

**TABLE S2:**
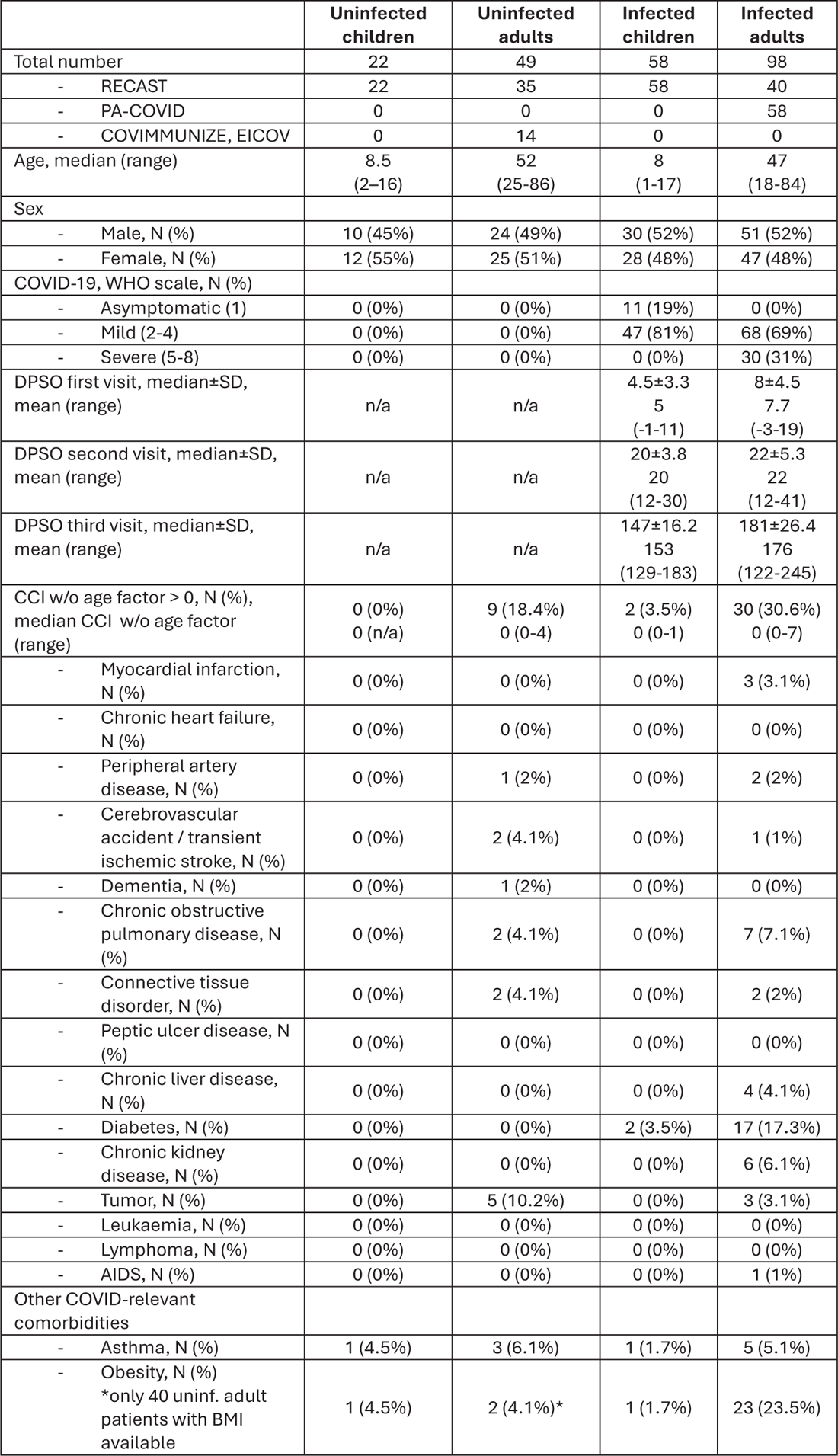
COHORT SUMMARY.

**TABLE S3:**
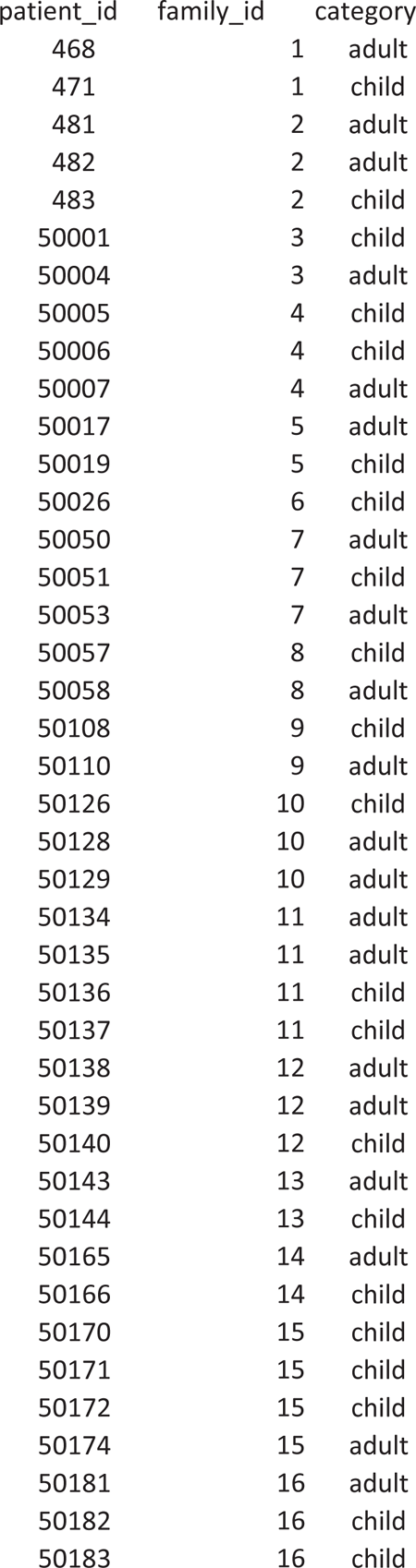
PATIENT FAMILY ID ASSIGNMENT WITHIN RECAST STUDY.

| patient_id | family_id | category |
| --- | --- | --- |
| 468 | 1 | adult |
| 471 | 1 | child |
| 481 | 2 | adult |
| 482 | 2 | adult |
| 483 | 2 | child |
| 50001 | 3 | child |
| 50004 | 3 | adult |
| 50005 | 4 | child |
| 50006 | 4 | child |
| 50007 | 4 | adult |
| 50017 | 5 | adult |
| 50019 | 5 | child |
| 50026 | 6 | child |
| 50050 | 7 | adult |
| 50051 | 7 | child |
| 50053 | 7 | adult |
| 50057 | 8 | child |
| 50058 | 8 | adult |
| 50108 | 9 | child |
| 50110 | 9 | adult |
| 50126 | 10 | child |
| 50128 | 10 | adult |
| 50129 | 10 | adult |
| 50134 | 11 | adult |
| 50135 | 11 | adult |
| 50136 | 11 | child |
| 50137 | 11 | child |
| 50138 | 12 | adult |
| 50139 | 12 | adult |
| 50140 | 12 | child |
| 50143 | 13 | adult |
| 50144 | 13 | child |
| 50165 | 14 | adult |
| 50166 | 14 | child |
| 50170 | 15 | child |
| 50171 | 15 | child |
| 50172 | 15 | child |
| 50174 | 15 | adult |
| 50181 | 16 | adult |
| 50182 | 16 | child |
| 50183 | 16 | child |

**TABLE S4:** MANUAL ANNOTATION MARKERS.

| gene | metalineage |
| --- | --- |
| PTPRC | B, DC, Mono, T |
| CD3E | T |
| CD3G | T |
| CD3D | T |
| CD4 | T, DC |
| CD8A | T |
| CD8B | T |
| TRAC | T |
| TRGC1 | T |
| TRGC2 | T |
| SELL | T |
| CD58 | T |
| CD248 | T |
| RUNX2 | T |
| HOPX | T |
| TOX | T |
| TNFRSF4 | T |
| TNFRSF18 | T |
| CCR5 | T |
| CCR7 | T |
| CD38 | B, DC, Mono, T |
| CD28 | T, B |
| CD27 | T, B |
| PDCD1 | T |
| LAG3 | T |
| TBX21 | T |
| CXCR3 | T, Mono |
| GATA3 | T |
| PTGDR2 | T |
| CCR4 | T |
| RORC | T |
| IL17A | T |
| CCR6 | T |
| IL2RA | T |
| FOXP3 | T |
| CTLA4 | T |
| BCL6 | T, B |
| CXCR5 | T, B |
| ICOS | T |
| EOMES | T |
| GZMA | T |
| GZMB | DC, T |
| GZMK | T |
| PRF1 | T |
| KLRC1 | T |
| KLRB1 | T |
| KLRG1 | T |
| KLRF1 | T |
| NKG7 | B, DC, Mono |
| CD19 | T, B |
| MS4A1 | T, B |
| CD79A | B |
| PXK | B |
| IGHM | B |
| IGHD | B |
| IGHG1 | B |
| IGHG2 | B |
| IGHG3 | B |
| IGHG4 | B |
| IGHA1 | B |
| CR2 | B |
| CD23 | B |
| CD24 | B |
| MME | B |
| MZB1 | B |
| SDC1 | B |
| CD14 | DC, Mono |
| ITGAX | T, B, DC, Mono |
| FCGR3A | DC, Mono, T |
| FCGR3B | DC, Mono, T |
| CD68 | Mono |
| APOE | Mono |
| CD69 | B, DC, Mono, T |
| CD74 | B, DC, Mono, T |
| HLA-DRA | B, DC, Mono, T |
| HLA-DRB1 | B, DC, Mono, T |
| HLA-DPA1 | B, DC, Mono, T |
| FCER1A | DC, Mono |
| IL3RA | DC, Mono |
| TLR7 | DC, Mono |
| CLEC4C | DC |
| MKI67 | B, DC, Mono, T |
| IFNAR1 | B, T |
| IFNAR2 | B, T |
| JAK1 | B, T |
| TYK2 | B, T |
| STAT1 | B, T |
| STAT2 | B, T |
| STAT5A | B, T |
| IRF9 | B, T |
| SOCS1 | B, T |
| SOCS3 | B, T |
| STAT3 | B, T |
| USP18 | B, T |
| SMAD2 | B, T |
| SMAD3 | B, T |
| S1PR1 | B, T |
| ISG20 | B, T |
| ZC3HAV1 | B, T |

**TABLE S4:**
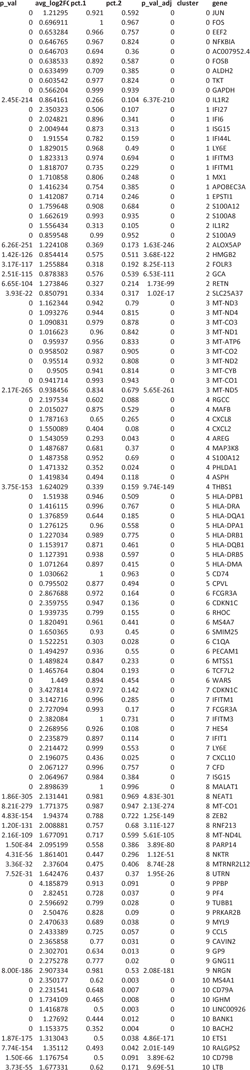
PBMC MONOCYTES CLUSTER MARKERS DE.

**TABLE S4:**
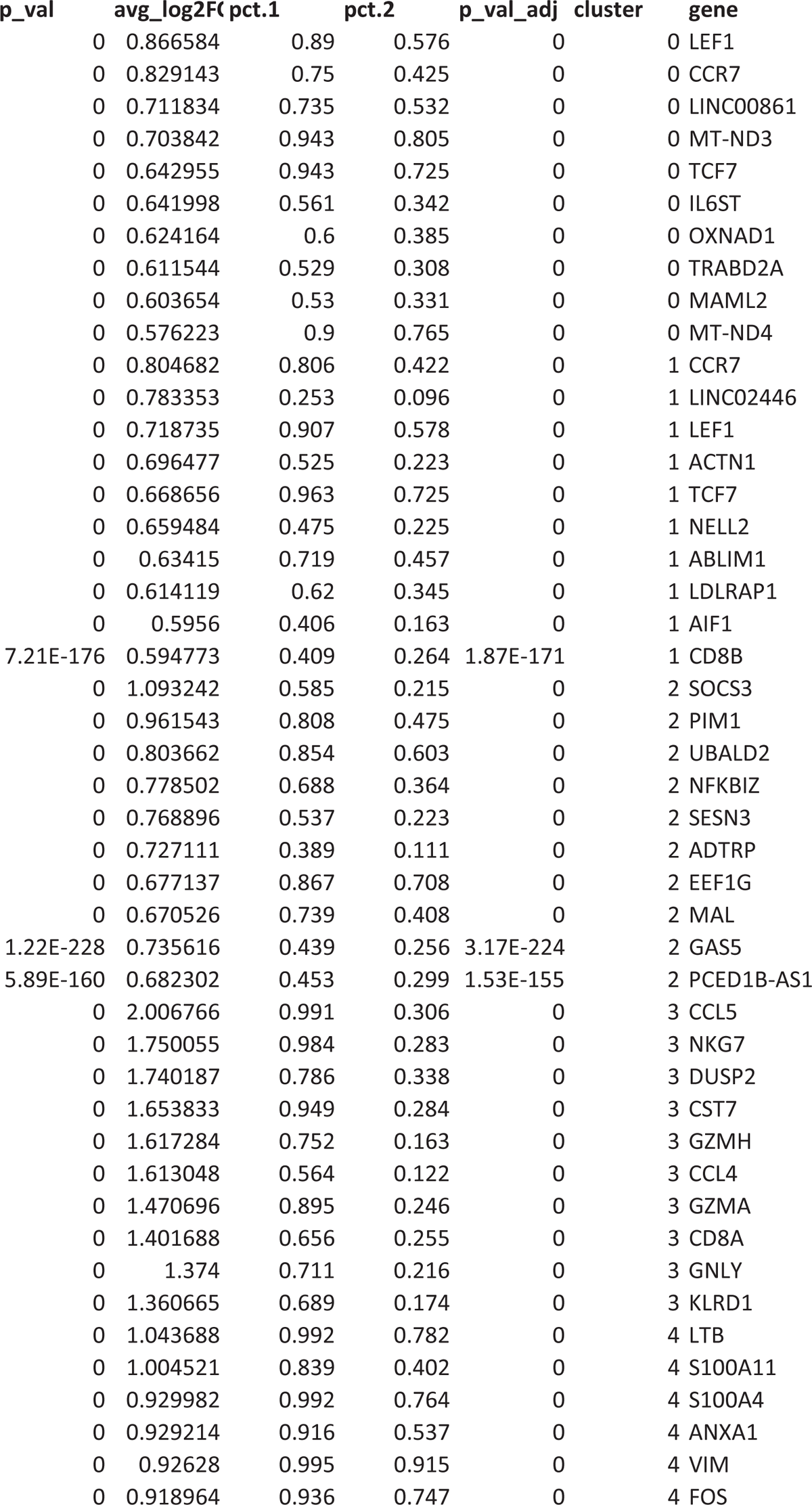
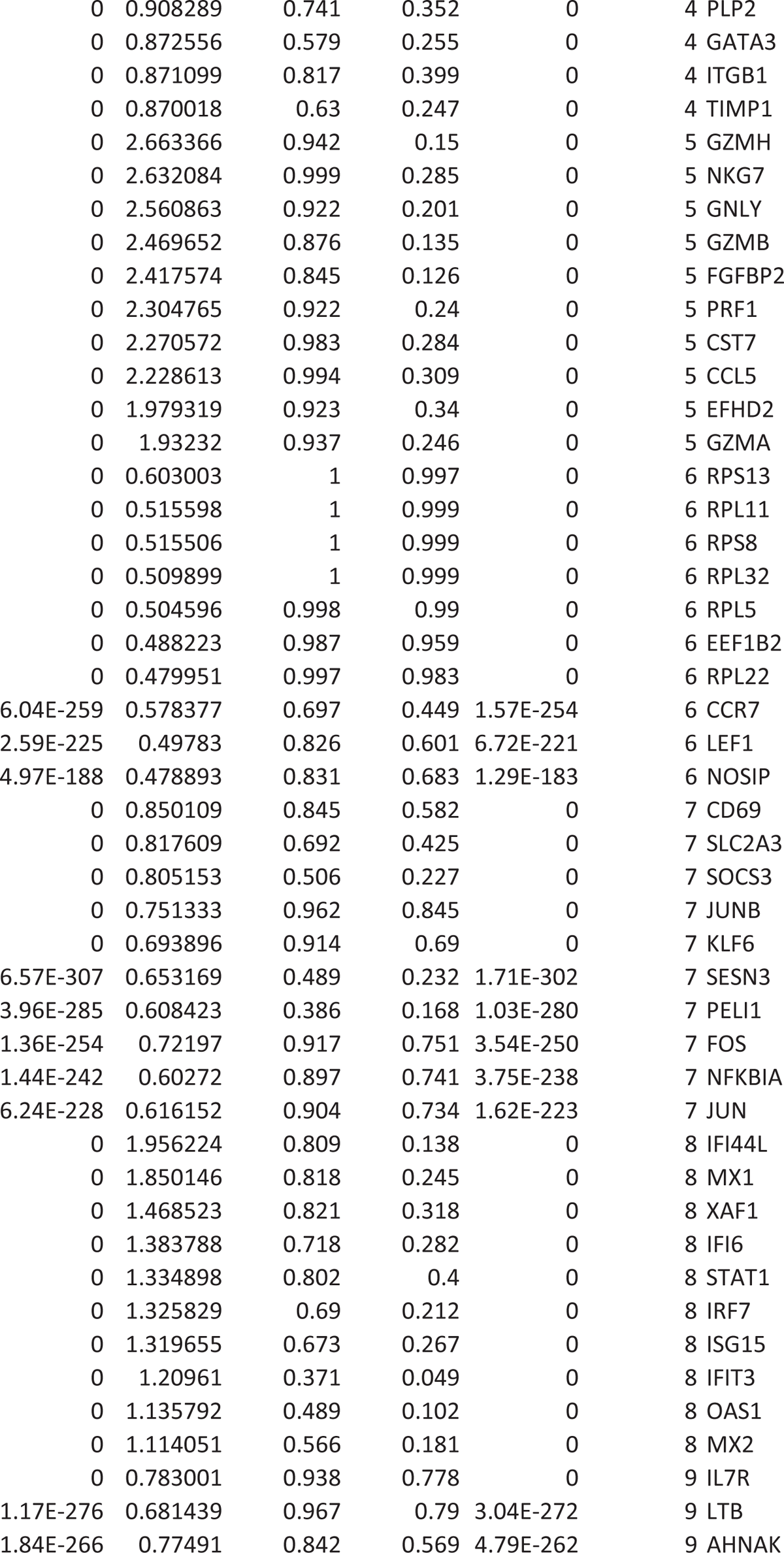
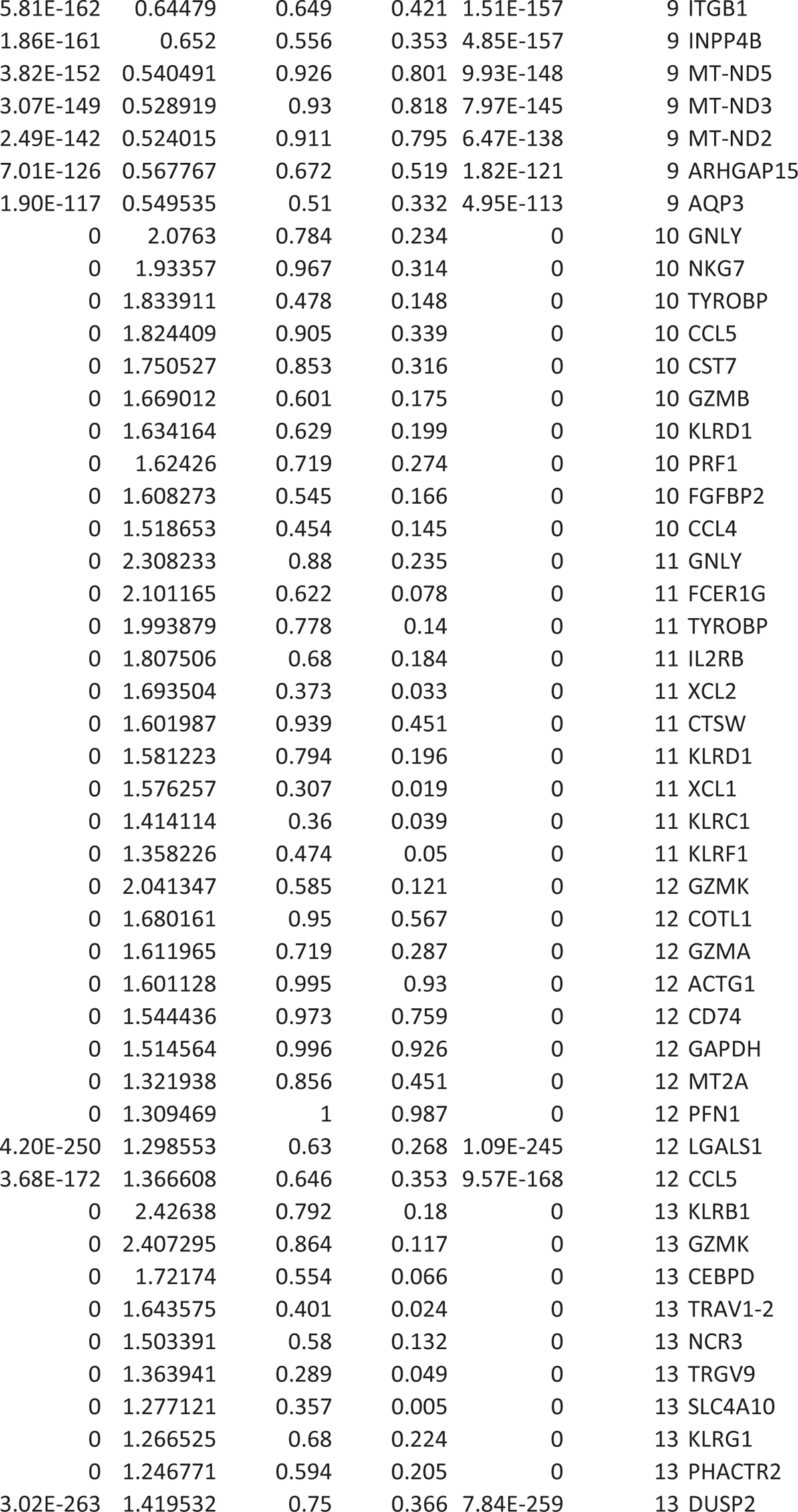
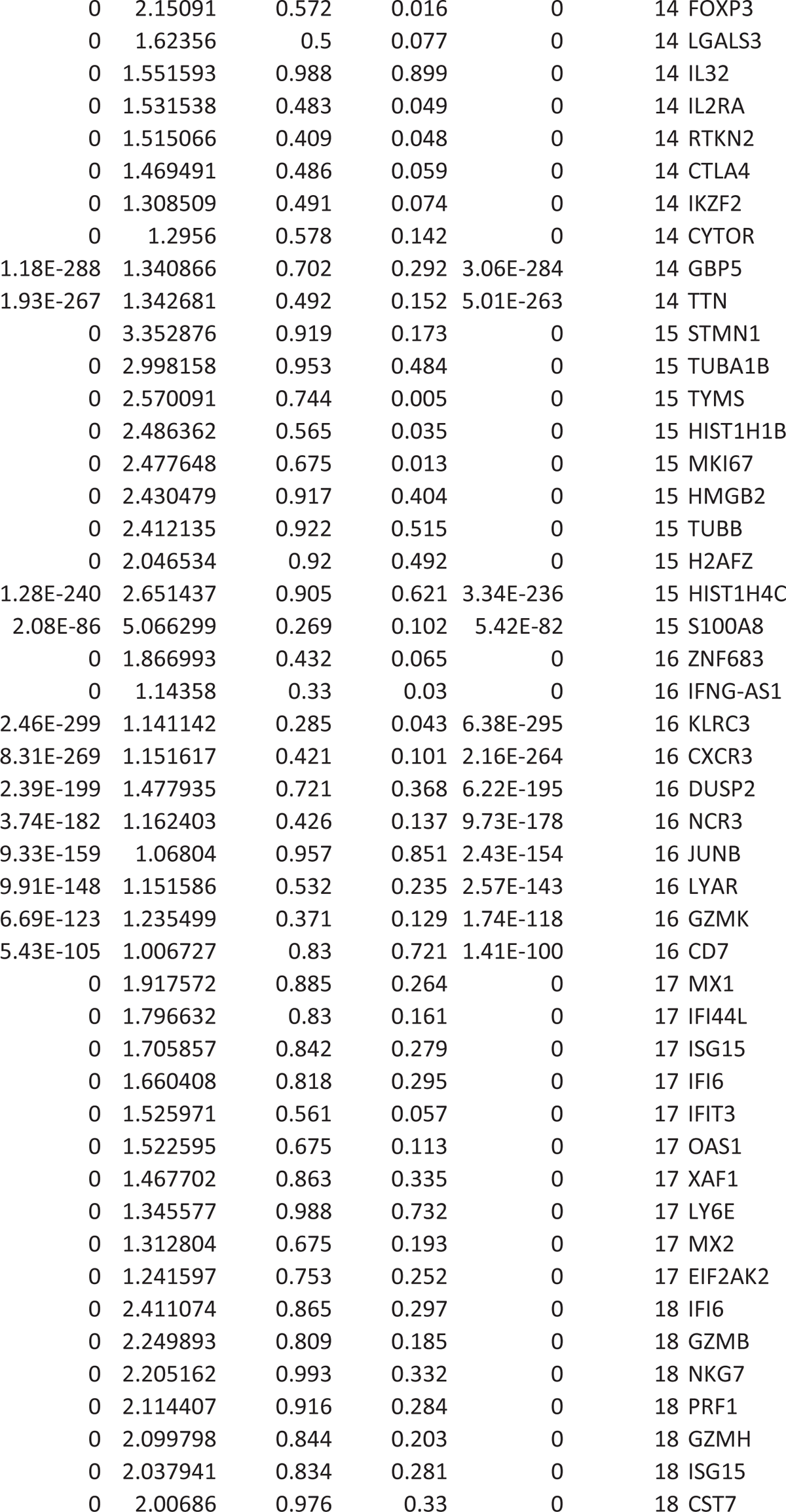
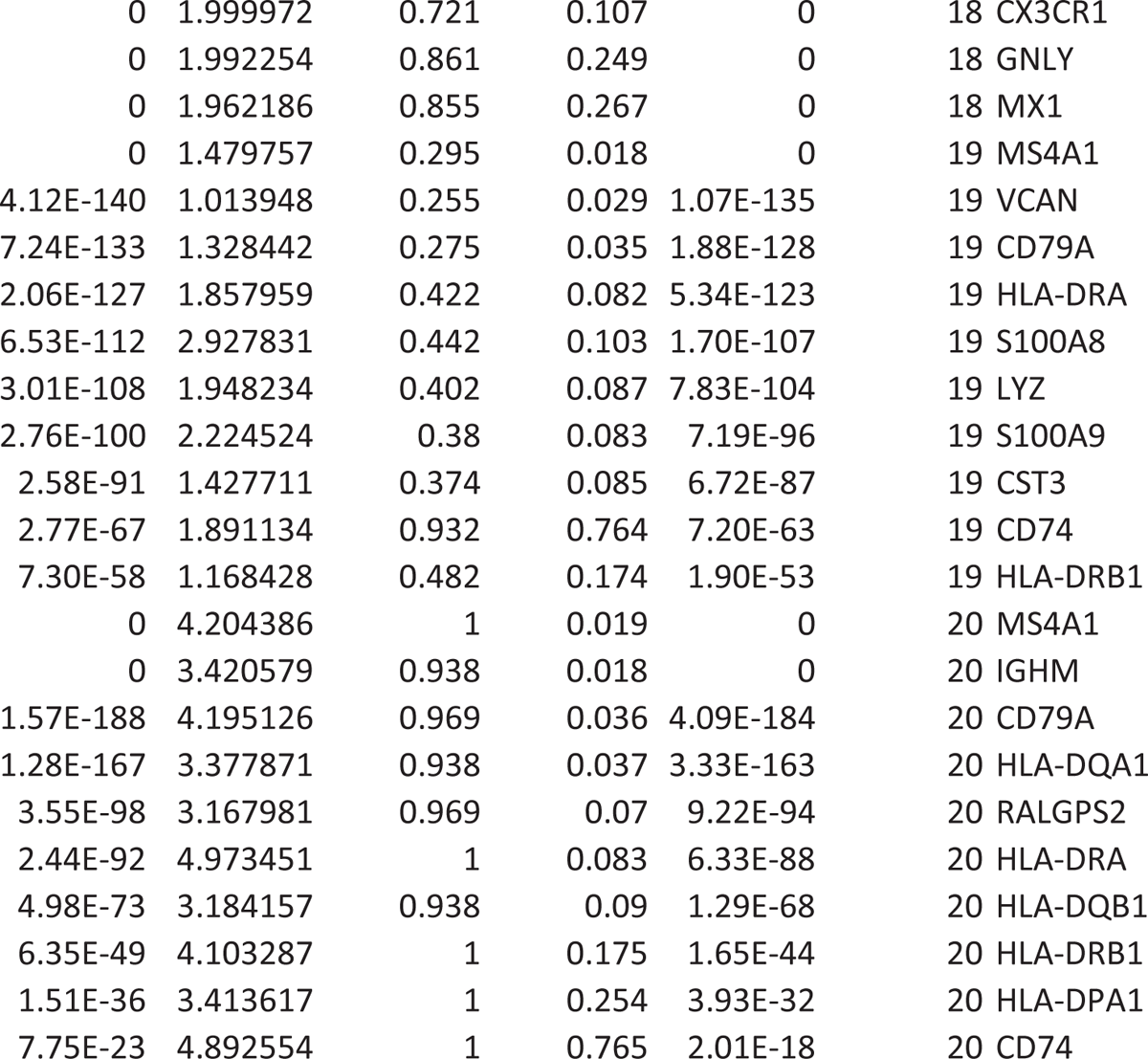
PBMC T CELLS CLUSTER MARKERS DE.

**TABLE S4:**
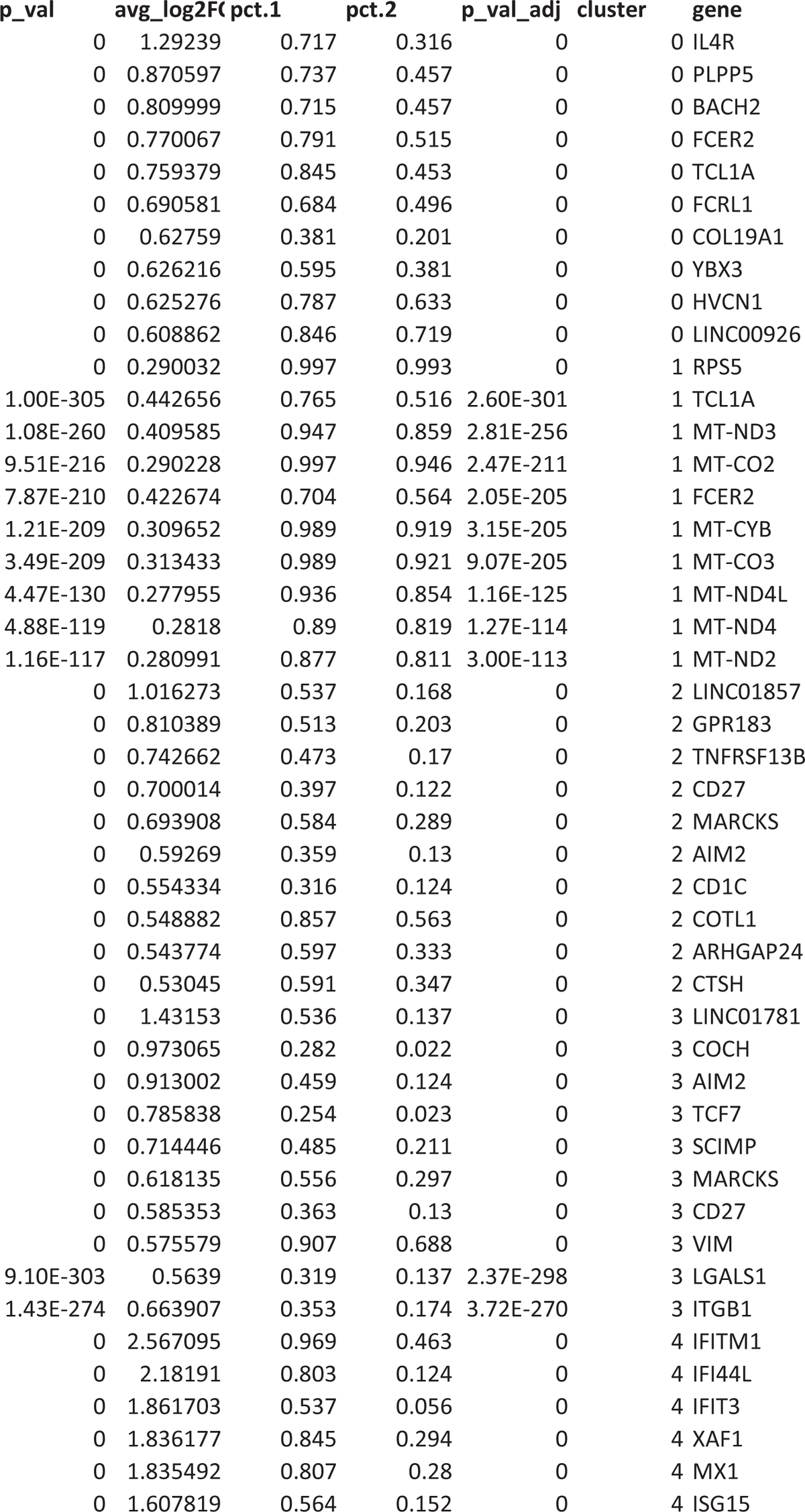
PBMC B CELLS CLUSTER MARKERS DE.

