## Supplementary Table 2 for "Rewired type I IFN signaling is linked to age-dependent differences in COVID-19"

|  | **Uninfected children** | **Uninfected adults** | **Infected children** | **Infected adults** |
| --- | --- | --- | --- | --- |
| Total number | 22 | 49 | 58 | 98 |
| - RECAST | 22 | 35 | 58 | 40 |
| - PA-COVID | 0 | 0 | 0 | 58 |
| - COVIMMUNIZE, EICOV | 0 | 14 | 0 | 0 |
| Age, median (range) | 8.5  (2–16) | 52  (25-86) | 8  (1-17) | 47  (18-84) |
| Sex |  |  |  |  |
| - Male, N (%) | 10 (45%) | 24 (49%) | 30 (52%) | 51 (52%) |
| - Female, N (%) | 12 (55%) | 25 (51%) | 28 (48%) | 47 (48%) |
| COVID-19, WHO scale, N (%) |  |  |  |  |
| - Asymptomatic (1) | 0 (0%) | 0 (0%) | 11 (19%) | 0 (0%) |
| - Mild (2-4) | 0 (0%) | 0 (0%) | 47 (81%) | 68 (69%) |
| - Severe (5-8) | 0 (0%) | 0 (0%) | 0 (0%) | 30 (31%) |
| DPSO first visit, median±SD, mean (range) | n/a | n/a | 4.5±3.3  5  (-1-11) | 8±4.5  7.7  (-3-19) |
| DPSO second visit, median±SD, mean (range) | n/a | n/a | 20±3.8  20  (12-30) | 22±5.3  22  (12-41) |
| DPSO third visit, median±SD, mean (range) | n/a | n/a | 147±16.2  153  (129-183) | 181±26.4  176  (122-245) |
| CCI w/o age factor > 0, N (%), median CCI w/o age factor (range) | 0 (0%)  0 (n/a) | 9 (18.4%)  0 (0-4) | 2 (3.5%)  0 (0-1) | 30 (30.6%)  0 (0-7) |
| - Myocardial infarction, N (%) | 0 (0%) | 0 (0%) | 0 (0%) | 3 (3.1%) |
| - Chronic heart failure, N (%) | 0 (0%) | 0 (0%) | 0 (0%) | 0 (0%) |
| - Peripheral artery disease, N (%) | 0 (0%) | 1 (2%) | 0 (0%) | 2 (2%) |
| - Cerebrovascular accident / transient ischemic stroke, N (%) | 0 (0%) | 2 (4.1%) | 0 (0%) | 1 (1%) |
| - Dementia, N (%) | 0 (0%) | 1 (2%) | 0 (0%) | 0 (0%) |
| - Chronic obstructive pulmonary disease, N (%) | 0 (0%) | 2 (4.1%) | 0 (0%) | 7 (7.1%) |
| - Connective tissue disorder, N (%) | 0 (0%) | 2 (4.1%) | 0 (0%) | 2 (2%) |
| - Peptic ulcer disease, N (%) | 0 (0%) | 0 (0%) | 0 (0%) | 0 (0%) |
| - Chronic liver disease, N (%) | 0 (0%) | 0 (0%) | 0 (0%) | 4 (4.1%) |
| - Diabetes, N (%) | 0 (0%) | 0 (0%) | 2 (3.5%) | 17 (17.3%) |
| - Chronic kidney disease, N (%) | 0 (0%) | 0 (0%) | 0 (0%) | 6 (6.1%) |
| - Tumor, N (%) | 0 (0%) | 5 (10.2%) | 0 (0%) | 3 (3.1%) |
| - Leukaemia, N (%) | 0 (0%) | 0 (0%) | 0 (0%) | 0 (0%) |
| - Lymphoma, N (%) | 0 (0%) | 0 (0%) | 0 (0%) | 0 (0%) |
| - AIDS, N (%) | 0 (0%) | 0 (0%) | 0 (0%) | 1 (1%) |
| Other COVID-relevant comorbidities |  |  |  |  |
| - Asthma, N (%) | 1 (4.5%) | 3 (6.1%) | 1 (1.7%) | 5 (5.1%) |
| - Obesity, N (%)   *only 40 uninf. adult patients with BMI available | 1 (4.5%) | 2 (4.1%)* | 1 (1.7%) | 23 (23.5%) |
